# scFLAME: a unified generative model for interpretable clustering, hierarchical structure discovery and marker-gene identification in single-cell RNA-seq data

**DOI:** 10.64898/2026.09.04.749363

**Authors:** Jackie Rao, Muntadher Jihad, Giulia Biffi, Paul D.W. Kirk

## Abstract

Identifying cell types from single-cell RNA sequencing (scRNA-seq) data typically requires several separate and often uninterpretable steps: dimensionality reduction, batch-correction, clustering, marker-gene identification and the discovery of finer-grained structure. Here we introduce scFLAME (single-cell Factor Latent Analysis with Mixture Embeddings), a probabilistic generative model that unifies these tasks: a negative binomial factor analysis of the raw counts - which can be adjusted for batch - is coupled to a Gaussian mixture prior over the latent space, learning the embedding and clustering jointly, while a shared linear decoder provides cluster-specific marker genes directly from the fitted model, and a merging procedure recovers a probabilistic hierarchy of finer-grained partitions. On simulated and real data, scFLAME matches or exceeds state-of-the-art clustering accuracy, is robust across sequencing platforms, and scales near-linearly to hundreds of thousands of cells. scFLAME thus replaces a chain of separate tools with a single, interpretable model for single-cell analysis.

## Introduction

Single-cell RNA sequencing (scRNA-seq) profiles transcription in individual cells, enabling cellular heterogeneity to be resolved and the development of reference atlases of whole tissues and organisms [1, 2]. The identification of cell types is a fundamental step in most downstream analyses [3], and is typically posed as an unsupervised clustering problem. The data are high-dimensional, sparse integer counts that exhibit overdispersion [4], for which the negative binomial (NB) distribution has become the model of choice [5, 6]. Their noise characteristics also vary across sequencing platforms, motivating count-based methods that are robust across technologies. However, biological variation in scRNA-seq data is not solely driven by discrete cell-type differences: other sources of variation, such as pathway activity, cell-cycle state and batch effects arising from differences in collection time, protocol or sequencing platform, can confound or mask the signal that clustering aims to recover.

Integrating data across batches offers substantial gains in biological insight but is complicated by systematic differences in gene expression as a result, including differences in cell type composition between batches, and expression differences within the same cell type across batches [7]. Such integration is often essential rather than incidental: studies frequently combine data from multiple donors to capture population-level variation, or from multiple timepoints to resolve dynamic processes such as disease progression. Batch correction methods are widely used to address this, ranging from approaches that directly adjust the count or expression matrix (e.g. ComBat [8]) to those that correct a lower dimensional embedding (e.g. Harmony [9], LIGER [10]). Batch and cell-type structure are often confounded, and recent benchmarks suggest current methods remain poorly calibrated [11]. Correcting for batch prior to clustering can also remove or mask rare cell populations [12], motivating methods that instead integrate batch correction directly into the cell-type discovery problem, for example by incorporating batch as a parameter within the count-generating model itself [13].

Dimensionality reduction is typically applied to improve statistical and computational tractability, and to overcome the curse of dimensionality. Most pipelines perform dimensionality reduction and clustering as separate, sequential steps: linear or non-linear embeddings (e.g. Principal Components Analysis (PCA) [14], Uniform Manifold Approximation and Projection (UMAP) [15]) are followed by *k*-means or graph-based community detection such as the Louvain and Leiden algorithms [16, 17]. Since the embedding is learned independently of the clustering objective (and may also assume Gaussian noise on transformed counts), information that distinguishes cell populations can be lost. Count-based factor models (e.g. ZINB-WaVE [18]) and deep generative models (e.g. scVI [19]) instead model counts directly, and a growing class of methods optimise embedding and clustering jointly, including Gaussian-mixture-prior approaches such as VaDE [20]. A comparison of some current methods is given in Supplementary Table S1. Notably, few methods combine count-based modelling, joint clustering, and interpretable gene programme discovery. The identification of marker genes - subsets of genes whose expression profiles distinguish specific cellular subpopulations - is essential for cell-type annotation. Conventionally, many pipelines rely on a separate post-hoc differential expression step to identify these markers, and methods such as the Wilcoxon rank-sum test remain top performers [21]. Only a handful of current models, such as the linearly-decoded LDVAE [22], recover gene programmes directly during dimensionality reduction process.

We introduce scFLAME (single-cell Factor Latent Analysis with Mixture Embeddings), which unifies these goals in a single probabilistic generative model. A negative binomial factor analysis of the raw counts is combined with a Gaussian mixture prior over the latent space, performing dimensionality reduction and clustering jointly, while a shared linear decoder makes cluster-specific marker genes identifiable directly from the fitted parameters, without post-hoc testing. Like VaDE [20], scFLAME places a mixture prior over the latent space, but instead of using deep encoder and decoder networks, scFLAME operates directly on counts through an interpretable linear decoder. We further introduce a merging procedure that yields a hierarchy of finer-grained partitions within the data, which may infer the existence of cell subtypes. Benchmarking against a broad range of methods on simulated and real data, we show that scFLAME matches or exceeds the state of the art on clustering accuracy, is robust across data-generating mechanisms and sequencing platforms, and scales near-linearly with cell number. scFLAME thus unifies a full single-cell workflow (dimensionality reduction, batch-aware clustering, sub-cluster discovery and marker-gene identification) within a single interpretable generative model.

## Results

### Overview of the scFLAME model

scFLAME is a probabilistic generative model that performs dimensionality reduction, clustering, batch correction and marker gene identification within a single framework (Figure 1a). It models raw counts directly using a negative binomial likelihood, a per-cell library-size offset to account for sequencing depth and an optional batch-correction term. Each cell is embedded as a continuous latent factor, and a Gaussian mixture prior over this latent space induces clustering, with a shared isotropic covariance across clusters (an Equal-Volume, Isotropic Gaussian Mixture Model; EII GMM). A loading matrix maps the latent space to the observed raw counts, providing a direct, interpretable link between the low-dimensional cell representation and gene programmes. Model fitting uses mean-field variational inference in two stages: a negative binomial factor analysis (NBFA) initialisation to learn the latent geometry, followed by joint optimisation of the mixture model (Figure 1b; see Methods).

**Figure 1.**
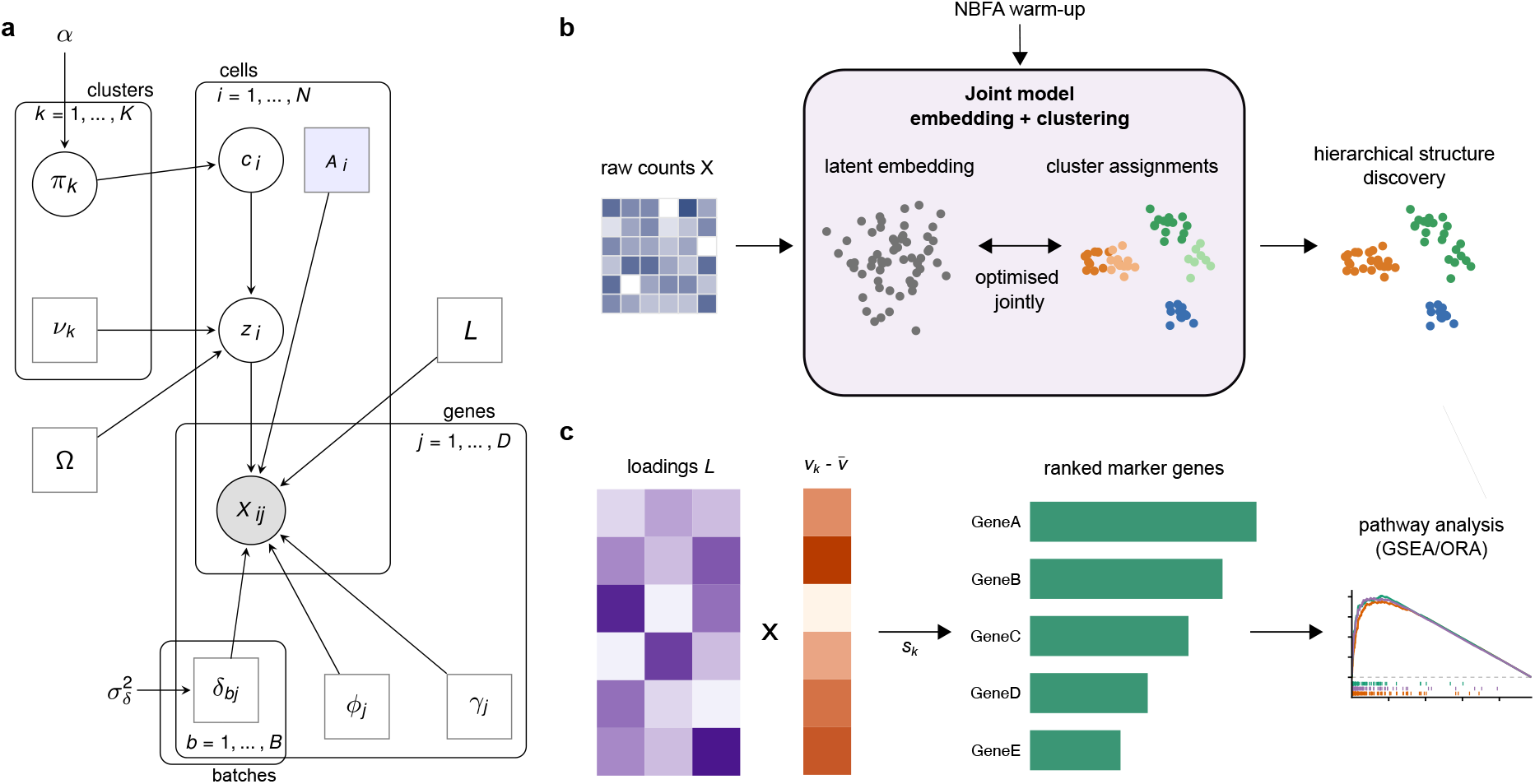
**a**, scFLAME couples a negative binomial factor analysis of the raw counts to a Gaussian mixture prior over the latent space. Mixture weights *π* are drawn from a Dirichlet prior with concentration *α*_0_; each cell *i* has a cluster indicator *c*_*i*_ *∼* Categorical(*π*) and a latent factor *z*_*i*_ | *c*_*i*_=*k ∼ N* (*ν*_*k*_, Ω), where *ν*_*k*_ is the mean of cluster *k* and Ω is a diagonal covariance shared across all clusters (an equal-volume, isotropic Gaussian mixture; EII GMM). Observed counts follow *X*_*ij*_ | *z*_*i*_ *∼* NB(*µ*_*ij*_, *ϕ*_*j*_) with log *µ*_*ij*_ = *γ*_*j*_ + (*Lz*_*i*_)_*j*_ + log *A*_*i*_ + *δ*_*bj*_, where *L* is a shared gene-loading matrix, *γ*_*j*_ a gene-specific intercept, *ϕ*_*j*_ a gene-specific dispersion, *A*_*i*_ a fixed, pre-estimated library-size offset and *δ*_*bj*_ a per-gene batch-correction term, with a 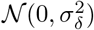 shrinkage prior. Shaded nodes are observed and solid dots denote fixed or pre-estimated quantities; plates index cells (*i* = 1, …, *N*), genes (*j* = 1, …, *D*), clusters (*k* = 1, …, *K*) and batches (*b* = 1, …, *B*). **b**, A negative binomial factor analysis (NBFA) warm-up is used only to initialise the model; the latent embedding and the cluster assignments are then optimised jointly, so that dimensionality reduction and clustering inform one another rather than being performed as separate sequential steps. **c**, Because the decoder is linear and shared across clusters, cluster-specific marker genes are read directly from the fitted model: combining the loadings *L* with the cluster offset 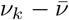 yields a per-gene score *s*_*k*_ (with 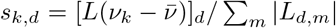 that ranks genes for each cluster and feeds downstream pathway analysis, without a separate differential-expression step.

Two notable features of scFLAME are: (1) cluster-specific marker genes can be identified directly from its fitted parameters as a *gene score* (Figure 1c, Methods), negating the need for a separate differential-expression step; and (2) holding the latent factors fixed, an optional merging procedure constructs a hierarchical tree of subtypes and their corresponding gene markers. We benchmark scFLAME against a broad range of methods on simulated and real data, and demonstrate its interpretability and recovery of cell-type hierarchies.

### scFLAME recovers latent structure in simulated data

We first benchmarked scFLAME on a series of synthetic and semi-synthetic datasets, for which the ground truth partition is known, thereby avoiding the circularity that can affect computationally-annotated real datasets (see below). We generated data from three sources: the scFLAME generative model itself, under settings of increasing difficulty, and the Splat and ZINB-WaVE simulators [18, 23], with parameters for both estimated from a real pancreas dataset from Baron et al. [24] (Methods). As a fully generative model, scFLAME can similarly simulate data based on real datasets, as we explore in the Supplementary Information.

scFLAME was compared against ten existing methods spanning graph-based (Louvain [16]), classical (PCA [14] + k-means [25]), deep-learning (scVI [19], LDVAE [22], scDeepCluster [26], scDSC [27]), count based factor (ZINB-WaVE [18]) and mixture model (VaDE [20], MFA [28]) approaches, plus the NBFA + EII GMM model used to initialise scFLAME (Methods). These methods capture commonly used pipelines as well as closely related generative modelling approaches. Each method was run three times on each of 20 datasets per scenario and evaluated using the adjusted Rand Index (ARI), normalised mutual information (NMI) and clustering accuracy (ACC) (Methods).

Across all scenarios, scFLAME ranked consistently among the best methods, whereas most competitors varied across data-generating models (Figure 2a,b). On the scFLAME-simulated datasets, several methods clustered the easy, well-separated scenario near-perfectly, but scFLAME dominated as difficulty increased: in the harder scenario it achieved a mean ARI of ≈ 0.88, against ≈ 0.66–0.69 for the deep-learning methods scVI, VaDE and scDSC. On Splat data, only scFLAME and NBFA + EII GMM approached the performance of manually-tuned Louvain (mean ARI *>* 0.75). MFA, PCA and VaDE all collapsed to ARI *≈* 0 and scDSC, scVI, LDVAE and scDeepCluster also exhibited poor performance. On ZINB-WaVE data, scFLAME and NBFA + EII GMM recovered the true structure perfectly (ARI = 1.0) despite having no explicit zero-inflation component, demonstrating a capacity for cross-model generalisation. Throughout, scFLAME generally improved on the NBFA + EII GMM baseline, validating the importance of the joint model over a two-step initialisation approach. Although Louvain with tuned *K* frequently ranked highly, this performance relied on manually tuning the resolution parameter; Louvain with its default resolution consistently ranked lower and led to over-clustering.

**Figure 2.**
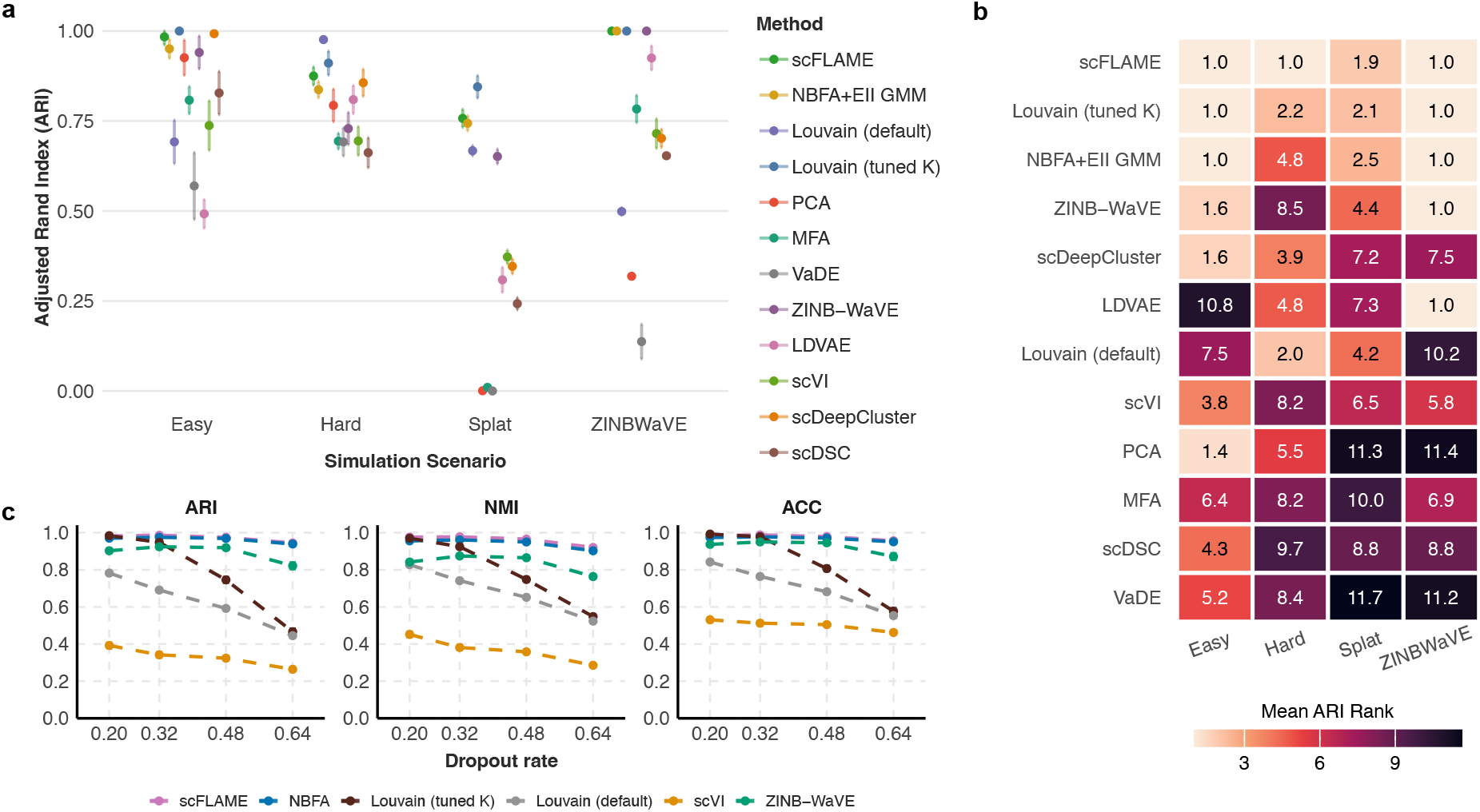
Comparative performance of scFLAME against competitor methods across simulated scenarios. **a**, Clustering accuracy measured by adjusted Rand Index (ARI) across four simulation scenarios: Easy, Hard, Splat, and ZINB-WaVE. Dots represent the mean ARI across 3 independent runs over 20 simulated datasets, with error bars indicating ± s.e. (standard error). **b**, Relative performance ranking of methods across simulation scenarios. The heatmap displays the mean rank (where 1 is the top-performing method) for the best performing replicate across 3 repeats for each method and each dataset. Lighter cell colours indicate superior performance (lower rank). **c**, Performance comparison of scFLAME against five baseline methods across increasing rates of simulated gene dropout in Splat simulated data with parameters from the Segerstolpe pancreas dataset (0.20 to 0.64; dropout.mid 1 to 4). Clustering performance is evaluated using three complementary metrics: ARI, NMI, ACC (Methods). Points represent the mean performance across *n* = 20 independent simulations with 3 repeats per method per dataset, with dashed lines indicating performance trends. Error bars indicate ± s.e.

scFLAME was also evaluated under increasing levels of technical sparsity and reduced clustering signal in Splat simulated data. We varied the technical dropout rate by changing the parameter ‘dropout.mid’, and generated simulations while deriving all other parameters from the Segerstolpe pancreas dataset Segerstolpe et al. [29]. We simulated datasets with dropout rates ranging from 19.8% to 63.6% and across this range, scFLAME and NBFA + EII GMM maintained ARI, NMI and ACC above 0.9 and outperformed every competitor at all levels, including ZINB-WaVE, which explicitly models dropout through a zero-inflation component (Figure 2c).

We additionally evaluated robustness to weaker biological cluster separation by reducing the de.facScale parameter in the Splat model. As cluster signal strength was weakened, scFLAME again outperformed all methods, and significantly exceeded NBFA + EII GMM at every setting, with the margin widening as noise grew (paired Wilcoxon signed-rank test, *p <* 0.001; Supplementary Figure S7). Together, these experiments show that scFLAME remains effective across a range of challenging simulation settings. Additional Splat and ZINB-WaVE settings including higher-sparsity regimes are reported in the Supplementary Information, with scFLAME maintaining a clear performance lead.

Finally, mini-batched stochastic optimisation allows scFLAME to scale to large datasets. On Splat simulated datasets ranging from 1,000 to 200,000 cells, wall-clock runtime scaled near-linearly with *N* (power-law exponent 0.89, *R*^2^ = 0.996), with the complete warm-up and training for scFLAME completing in approximately 39 minutes for 200,000 cells. Clustering accuracy stayed high (median ARI *>* 0.99; Supplementary Figure S8). Clustering runtimes relative to other methods on a real dataset are reported in Supplementary Table S3; we note that, unlike scFLAME, some baselines require a separate dimensionality reduction step and most do not natively support marker-gene identification, so their reported times do not reflect the cost of a full, comparable analysis.

### scFLAME identifies known cell types in benchmark single-cell datasets

We next applied scFLAME to eight publicly available scRNA-seq datasets spanning five sequencing platforms and a wide range of size, sparsity and biological complexity, from 1,700 to 68,000 cells (Table 1, Methods). Using the raw counts as input, we compared scFLAME to the same ten methods as above for the seven smaller datasets, each run five times starting at the reported number of cell types for each dataset and evaluated over 5 seeds. We took the best run for methods with an internal model-selection criterion, or took the first if this did not exist; some baselines (e.g. Louvain, VaDE) could not reliably attain the correct cluster count across seeds (Methods). We further applied scFLAME to two additional datasets comprising multiple batches and sequencing platforms to evaluate the batch-correction term under real technical variation.

**Table 1:** Summary of real datasets used for evaluation. All datasets were filtered to the top 5,000 most variable genes after any pre-processing described in Methods, except the Tabula Muris (TM) spleen datasets, which were filtered to 8,000 genes to compare methods on a higher-dimensional dataset. # Clusters refers to the number of groups in the ground truth clustering that we compare to. % zeros refers to the filtered dataset.

| Dataset | Sequencing platform | Filtered # cells | Original # genes | # Clusters | % zeros |
| --- | --- | --- | --- | --- | --- |
| PBMC 4K [30] | 10X Genomics | 3857 | 33694 | 8 | 93.0% |
| Baron human pancreas [24] | inDrop | 1895 | 20125 | 9 | 70.0% |
| Zeisel mouse brain cortex [31] | STRT-Seq (UMI) | 2816 | 20006 | 7 | 46.8% |
| Segerstolpe human pancreas [29] | Smart-seq2 | 1992 | 26179 | 7 | 33.1% |
| Mouse uterus [2] | Microwell-seq | 3392 | 21848 | 10 | 86.5% |
| Mouse spleen (10X) [32] | 10X Genomics | 7022 | 20138 | 6 | 91.6% |
| Mouse spleen (Smart-seq2) [32] | Smart-seq2 | 1693 | 22966 | 5 | 90.9% |
| PBMC 68K [30] | 10X Genomics | 68165 | 32738 | 10 | 97.6% |

Across the eight benchmark datasets (Table 1) and all clustering metrics, scFLAME was consistently among the top methods, achieving the best median rank for both ARI and accuracy, and coming second for NMI (Figure 3a; Supplementary Figure S10). Performance held across platforms and sparsity levels, whereas competitors varied widely. scFLAME and the NBFA+EII GMM baseline were near-perfect on Segerstolpe (*>* 0.98 on all metrics) and clearly best on the TM spleen SS2 data (ACC 0.78, versus 0.67 for Louvain). The strongest competitors were the other count-based latent-variable models: LDVAE was the top method on PBMC (ARI 0.768; scFLAME 0.757), and ZINB-WaVE performed well despite not being designed for clustering. The poorest were PCA, MFA and VaDE, which operate on transformed data (VaDE frequently collapsing to a single cluster), motivating the use of methods operating directly on count data. Notably, scFLAME’s accuracy was not degraded by its lack of a zero-inflation component, even on the sparsest data. For exploratory analyses where computational resources are limited, NBFA+GMM may provide a useful alternative, but across the same seed, scFLAME usually improves upon the initial clustering configuration.

**Figure 3.**
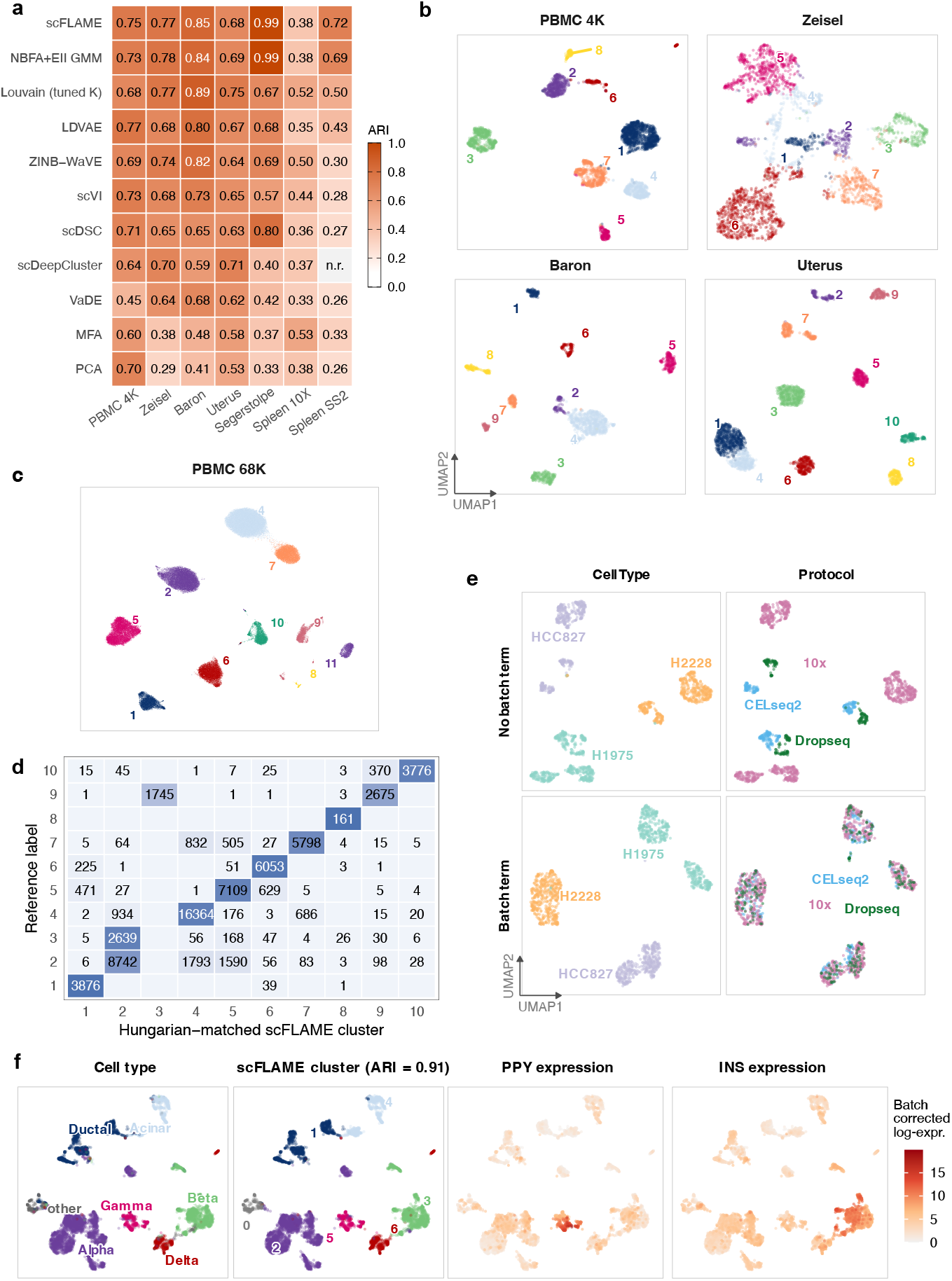
scFLAME recovers cell-type structure across diverse scRNA-seq datasets, including a large-scale 68k-cell dataset, and performs batch correction effectively. **a**, ARI between inferred and true cell-type labels for scFLAME and comparator methods across seven datasets where the ‘best’ of 5 or the first run is taken as detailed in Methods. ‘n.r’ indicates not run; the method ran into numerical errors. Rows are ordered by the median rank of the method across all datasets. **b**, UMAP projections of scFLAME latent embeddings for the PBMC 4K, Zeisel, Baron and Uterus datasets, coloured by scFLAME cluster. UMAP visualisations are labelled with scFLAME cluster numbers; for correspondence between these cluster numbers and reference cell type identities, see confusion matrices in Figure S12 and cell type mapping in Table 2.2. **c**, UMAP projection of scFLAME latent embeddings for the PBMC 68K dataset, coloured and numbered by scFLAME cluster assignment. **d**, Confusion matrix showing the proportion of cells in each reference cluster assigned to each Hungarian-matched scFLAME cluster for the PBMC 68K dataset. Cell counts are annotated in each tile. **e**, UMAP projections of scFLAME latent embeddings demonstrating batch correction across cell lines and sequencing protocols for the LUAD dataset. **f**, UMAP projections showing reference cell types, scFLAME cluster assignments (ARI = 0.91), and batch-corrected log-expression levels for marker genes *PPY* and *INS* for the Pancreas dataset, defined as the posterior mean of *µ*_*ij*_, evaluated at the reference batch level (*δ*_*bj*_ = 0).

Louvain achieved the best scores on the Baron and Uterus datasets, and generally performed well. This is expected, and should be read with care: graph-based clustering is the field standard, and the reference annotations are often generated by Louvain clustering, giving it an intrinsic advantage. More generally, since real-data cell type annotations are often derived computationally, these results are best considered in conjunction with the simulations above, for which the ground truth is unambiguously defined. On the highly imbalanced TM spleen 10X dataset (66% B cells), scFLAME’s lower ARI reflects biologically coherent subdivision of the dominant population rather than error, with its sub-clusters matching the finer annotations the authors themselves provide. In 3 of 5 runs, scFLAME achieved higher ARI values (0.527, 0.518, and 0.518), where the latter (with the second-highest ELBO) is visualised (Supplementary Figure S13).

Hungarian-matched confusion matrices and UMAP embeddings (Figure 3b,c) show that scFLAME recovers the annotated cell types across many datasets, and separates them more cleanly than the NBFA + EII GMM baseline (Supplementary Figure S11), indicating that the joint objective improves the embedding. In the PBMC dataset, scFLAME performs well relative to reference graph-based clustering, with minor mixing occurring only between NK and T-cell clusters, consistent with their closely related transcriptional programmes [33]. Similarly, in the Baron dataset, scFLAME recovers the majority of reference labels with high fidelity, while only mixing the activated (Cluster 2) and quiescent (Cluster 8) stellate populations two transcriptional states of the same underlying cell type. Although the mouse uterus dataset contains two batches, its ground-truth clusters are largely batch-specific, so it was evaluated without batch correction here; despite this, scFLAME successfully separates two stromal populations (Clusters 3 and 5) confounded by batch in other methods’ latent spaces. Applying batch correction did not improve performance due to the high confounding between reference labels and batch identity (Supplementary Figure S15). Although scFLAME did not separate microglia in the Zeisel dataset, these were not consistently resolved as a distinct cluster by any method, consistent with known difficulty in this complex atlas. We further identified characteristic failure modes of some of the deep methods: scDeepCluster over-separated cells into visually distinct islands, but these did not necessarily correspond to improved biological fidelity, where it was amongst the poorest performers on the Zeisel dataset despite achieving the most separated latent space. scDSC often merged distinct lineages, and split distinct populations such as B-cells in the PBMC dataset (Supplementary Figure S11). Additional per-dataset confusion matrices and embeddings for competing-method embeddings are provided in the Supplementary Information.

We applied scFLAME to the PBMC 68K dataset using mini-batching, recovering the annotated populations at competitive accuracy compared to analyses in the literature (ARI 0.66, NMI 0.73, ACC = 0.80; see e.g. [26]) and confirming that the method scales to large data (Figure 3c,d). At this scale we replaced the TMM library-size estimate with a simple total-count sum, with negligible loss of accuracy.

We next evaluate the effect of including a batch-correction term in scFLAME on two datasets where cell types are represented across multiple batches spanning multiple sequencing platforms; see Methods for further information about the datasets. In the LUAD dataset, three cancer cell lines (HCC827, H1975, H2228) were each profiled by three protocols exhibiting varying sparsity levels (CELseq2, 10x, Drop-seq), yielding a rare setting in which cell-type identity is known for all 1,401 cells [34]. Without the batch term, while scFLAME recovers an almost perfect clustering structure (ARI=0.98, best of 5 runs), the latent space separates almost entirely by protocol, with each cell type forming three distinct, protocol-specific sub-clusters (Figure 3e, top). Adding the batch term merges these sub-clusters into a single cluster per cell type, mixing all three protocols within each while still cleanly separating the three cell lines (Figure 3e, bottom) (ARI=0.99).

The Pancreas dataset combines four publicly available studies profiled by CEL-seq, CEL-seq2, and Smartseq2, comprising both islet cell types (alpha, beta, gamma, delta) and non-islet cell types (acinar, ductal) amongst others. Unlike LUAD, not every cell type is represented in every batch, though the dataset retains sufficient overlapping cell-type composition across studies for batch correction to be well identified (in contrast to the Uterus dataset above). With the batch term, scFLAME achieves an ARI of 0.91 against the annotated cell-type labels (Figure 3f), separating both the major islet/non-islet division and the finer cell-type structure within each. This outperforms scFLAME without batch correction, as well as a widely used batch-correction method - Harmony [9] - followed by Louvain (Supplementary Figure S14) and previously reported analyses [13]. Due to the interpretable nature of the negative binomial mean *µ*_*ij*_ in scFLAME’s generative model, batch-corrected posterior mean expression can be obtained directly from the fitted model by setting *δ*_*bj*_ to zero, without a separate post-hoc correction step. Batch-corrected marker gene expression in the latent space is concentrated within its corresponding cluster; for example, *PPY* in the Gamma cluster and *INS* in the Beta cluster (Figure 3f, Supplementary Figure S14). To assess whether batch correction improves separation of cell types independently of clustering accuracy, we computed the silhouette coefficient of each cell in the latent space with and without the batch term [35], using ground-truth cell-type labels as clusters. Average per-cell-type silhouette width increased with batch correction for all seven annotated cell types (paired t-test, *p* = 0.0025; Wilcoxon signed-rank, *p* = 0.016), indicating that the batch term reduces within cell-type, between-batch variance while preserving separation. Harmony correction [9] evaluated in a much higher-dimensional embedding (30 vs. 15 dimensions) produced lower silhouette widths than scFLAME’s batch-corrected latent space for all cell types (Supplementary Table S4).

Ablations confirmed that pre-estimating gene dispersions (with edgeR) and library sizes each sharpen the resolution of closely related populations. For example, gamma and delta pancreatic cells separate only when dispersions are pre-estimated (Supplementary Figure S18). A further study of cluster initialisation sensitivity from the NBFA latent space is provided in the Supplementary Information. Runtimes for all methods on the PBMC dataset are reported in Supplementary Table S3. For scFLAME, runtime scaled near-linearly with the number of genes (*R*^2^ *>* 0.99), while clustering accuracy plateaued beyond 5,000 genes; including more genes increased computation without improving accuracy, likely due to these genes having little biological signal (Supplementary Figure S19).

### Integrated marker gene identification via shared loading matrix

A key advantage of scFLAME is that interpretable, cluster-specific gene signatures emerge directly from the fitted model, without any post-hoc differential-expression step. For each cluster we define a *gene score s*_*k*_ that projects the cluster’s latent-mean offset *ν*_*k*_ through the shared loading matrix *L*, giving the scaled log fold-change of each gene relative to the population mean (Methods). Ranking genes by this score yields a marker list for every cluster, and the same construction applied to a pair of clusters ranks the genes that distinguish them.

Applied to the PBMC clusters, the top-scoring genes by *s*_*k*_ recover canonical cell-type markers and allow each cluster to be labelled directly (Figure 4a,b). For example, Cluster 3 was characterised by B-cell markers with *IGHM, CD79A, IGHD, MS4A1, VPREB3* making up the top 5 genes, while Cluster 4 showed enrichment of naive T-cell markers including *CCR7* and *LEF1*, consistent with established PBMC signatures [36]. While *KLRF1* and *FGFBP2* identified Cluster 5 as NK-cells, *NKG7* was shared with Cluster 0, where its association with cytotoxic activity supported the annotation of this cluster as CD8^+^ effector memory T cells [37]. Pairwise scores distinguishing two specific clusters are equally interpretable: contrasting the classical monocyte cluster (Cluster 2) with the CD16^+^-monocyte cluster (Cluster 7) recovers expected discriminating genes. *FCGR3A* (rank 1) cements the CD16^+^ monocyte annotation, whereas *S100A8, S100A12* and *S100A9* are known markers for classical monocytes. (Supplementary Figure S21b) [38].

**Figure 4.**
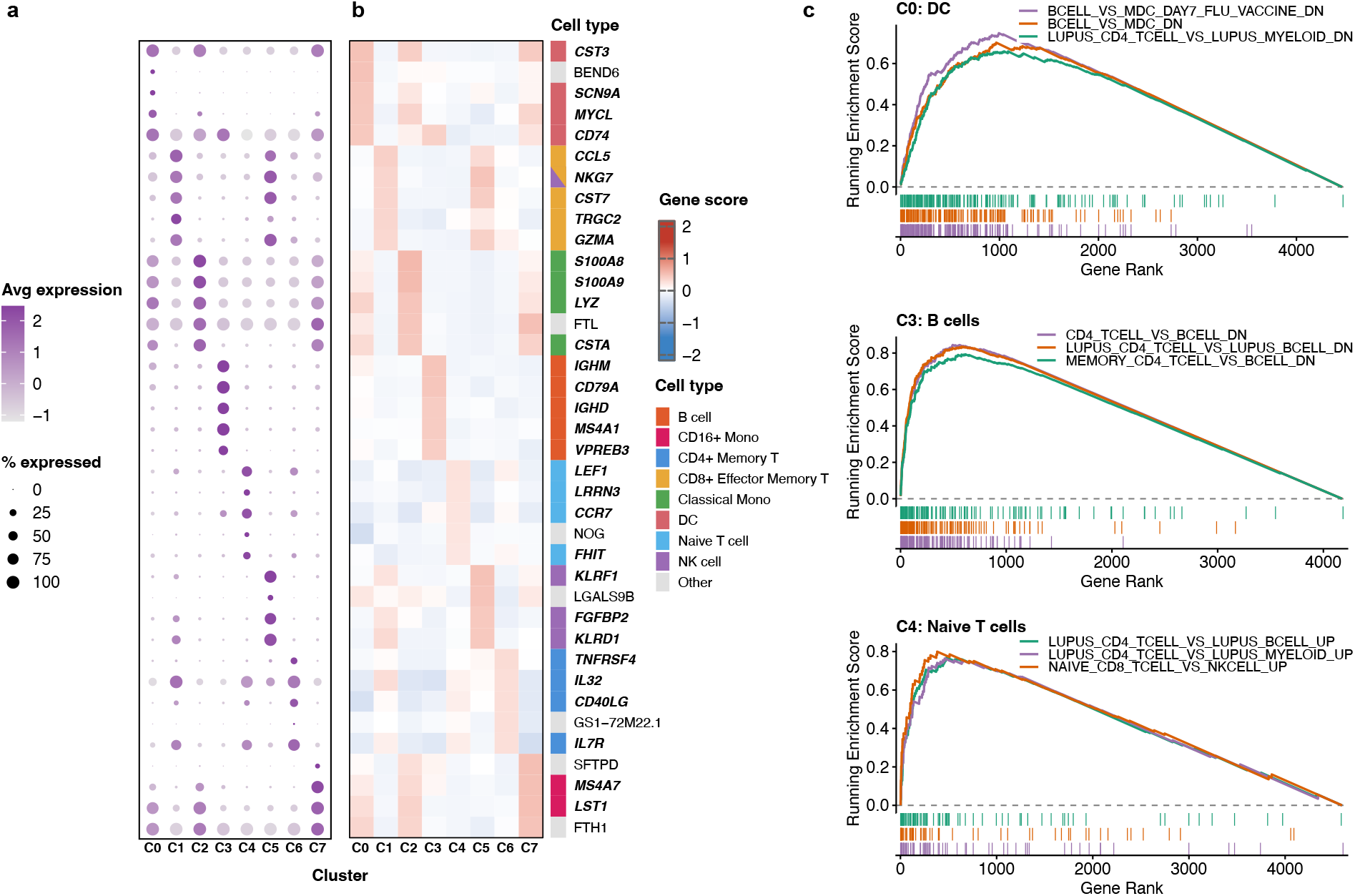
Characterisation of PBMC clusters identified by scFLAME via gene scores derived from the fitted scFLAME model. **a**, Dot plot showing expression of identified cluster marker genes across eight clusters (C0–C7). Dot size indicates the percentage of cells expressing each gene; colour indicates average scaled expression. **b**, Heatmap of gene scores for the top five marker genes per cluster, annotated by known cell-type identity (right). Bold italic gene names indicate canonical markers from prior literature [36]. Note poorly characterised RP11-annotated transcripts were excluded from visualisation, *NKG7* appears in the top 5 for both C1, C5 and FTL appears in the top 5 for C2, C7. **c**, Gene Set Enrichment Analysis (GSEA) enrichment plots for three selected clusters showing the top three enriched immunological gene sets by *p*.*adjust* (MSigDB C7), restricted to GEO-derived signatures (GSE). Running enrichment scores are shown as a function of gene rank, where genes were ranked for GSEA based on scFLAME gene scores *s*_*k*_.

Characterising the ranked score lists using gene set enrichment analysis with the MSigDB C7 gene set [39] supported these annotations at the pathway level, with each cluster’s top enriched immunological gene sets matching its assigned identity (Figure 4c). For example, Cluster 6 showed strong enrichment for CD4^+^ T-cell identity over B and myeloid lineages (NES *>* 3.0, *p*_*adj*_ *<* 10^−23^), and also captured sub-lineage maturation state of the population, where we saw positive enrichment for memory over naive phenotypes. Full enrichment results, Gene Ontology over-representation analysis (Supplementary Figure S21a), and the equivalent analysis for the Zeisel dataset are provided in the Supplementary Information.

To benchmark against conventional approaches, we assessed the concordance between our top-*N* ranked genes and the marker genes identified by Seurat’s post-hoc Wilcoxon rank-sum test on the top 5000 HVGs across a range of thresholds (*N ∈* [10, 100]). Our scores achieved high concordance (Jaccard similarity up to *∼*0.67; Supplementary Figure S21c), yet required no additional post-hoc computation once scFLAME had been fitted. Where concordance was lower, Seurat’s list was dominated by non-informative structural transcripts (e.g. ribosomal genes *RPS2, RPS18*), whereas scFLAME’s ranking recovered coherent, cell-type-specific programmes that simple fold-change thresholds overlooked.

### scFLAME reveals hierarchical structure and cellular sub-clusters

scFLAME incorporates a merging procedure which provides a hierarchical decomposition of the latent space, allowing coarser and finer structure - such as broad cell types and their constituent subtypes - to be explored at multiple resolutions. Beginning from an over-specified mixture with *K*_*init*_ components, clusters are iteratively merged by Integrated Classification Likelihood (ICL), leveraging our Bayesian framework, yielding a sequence of nested partitions. The resulting merge tree encodes a data-driven cell type hierarchy: fine-grained partitions at large *K* may capture transcriptional subtypes, while coarser partitions at small *K* recover broad cell types. Each level of the resulting hierarchy inherits the model’s interpretable gene scores *s*_*k*_, and partition stability along the merge path is assessed via the Davies–Bouldin index (DBI).

Applied to the PBMC 4K dataset, initialising at *K*_init_ = 15, the merge tree reveals a consistent hierarchical structure, with closely related sub-clusters merging early, while well-separated populations remain distinct until later stages (Figure 5a, b). At *K* = 8, the recovered clustering closely matches the annotated cell types from the marker-gene analysis above (ARI = 0.94) with consistent marker genes. The DBI is instead lowest at *K* = 9 (Figure 5f), where a non-classical monocyte/DC mixed cluster further splits into two populations (see below), suggesting a finer resolution captures additional, well-separated biological structure.

**Figure 5.**
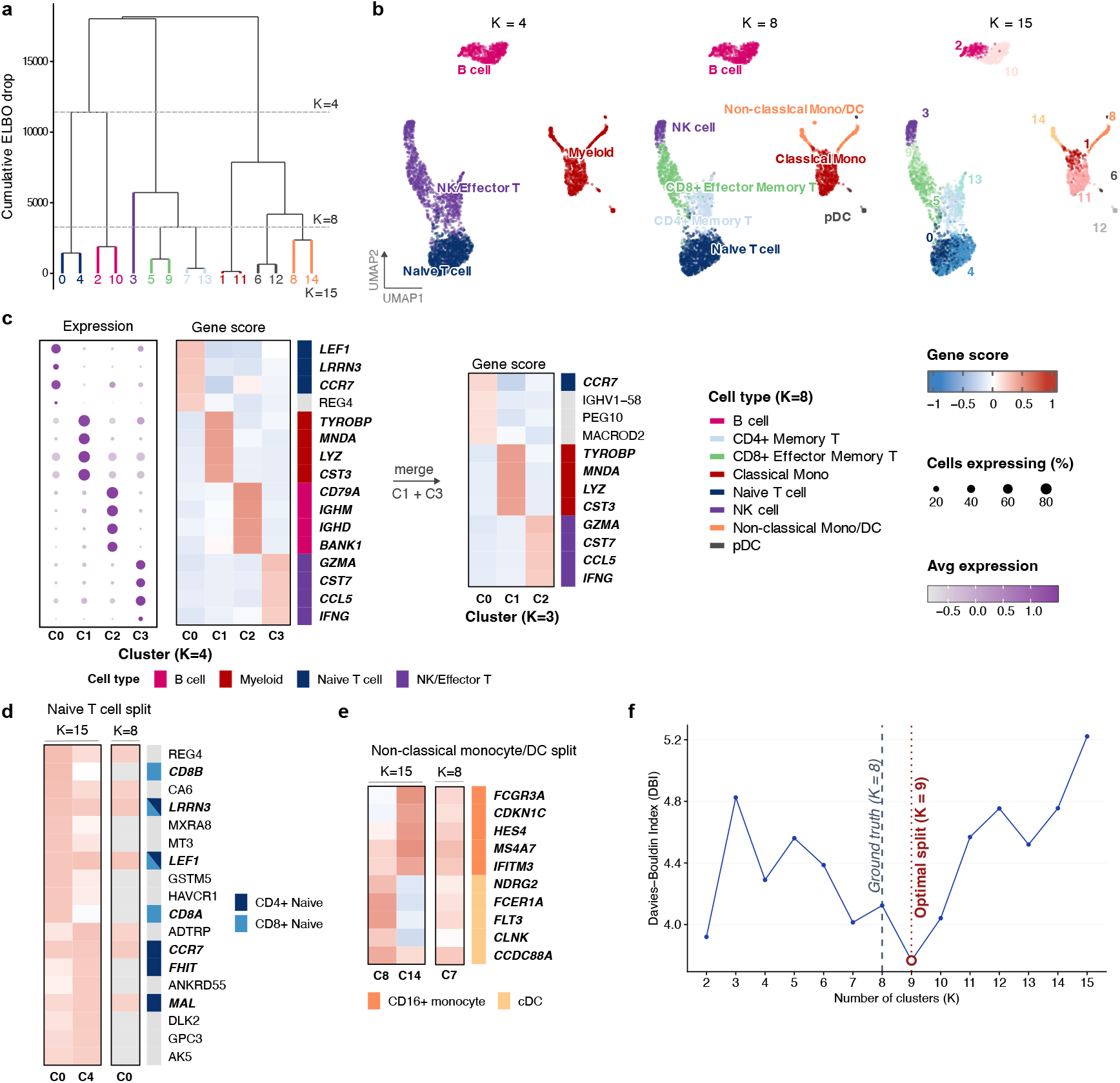
A greedy merge procedure recovers a biologically coherent hierarchy of sub-clusters in human PBMCs. **a**, Dendrogram showing the order of cluster merges. Branches are coloured by their *K* = 8 parent cell type; leaves are labelled with their *K* = 15 cluster index. Dashed line indicates *K* = 8. **b**, UMAP projections of the PBMC 4K dataset coloured by cluster assignment at three resolutions of the merge hierarchy for the first run of 3, starting from *K*_init_ = 15: *K* = 4, *K* = 8, and *K* = 15. Cluster identities were assigned post hoc using marker gene scores *s*_*k*_(Methods). Other runs were qualitatively consistent. **c**, Marker genes distinguishing broad clusters. Left: dot plot of gene expression for the top 4 marker genes per cluster at *K* = 4. Right: gene scores *s*_*k*_ for the same clusters before and after merging into *K* = 3, annotated by inferred cell type. **d**, Gene scores distinguishing naive CD4+ and CD8+ T cell clusters at *K* = 15 (clusters 0 and 4), showing the top 10 marker genes, and their merged counterpart at *K* = 8 (cluster 3), where genes appearing in the top 10 for Cluster 0 are highlighted with their score. *LEF1, LRRN3* are shared marker genes for both clusters at *K* = 15. **e**, Gene scores distinguishing non-classical monocyte and dendritic cell subclusters at *K* = 15 (clusters 8 and 14) and their merged counterpart at *K* = 8 (cluster 7). We show 5 key marker genes for each cell type in both cases. **f**, Davies–Bouldin index across merge steps from *K*_init_ = 15 down to *K* = 2. The minimum is at *K* = 9, where the reference labels have *K* = 8.

At coarser resolutions (*K* = 3, *K* = 4), NK cells and effector/memory T cells group together, separate from naive T cells (Figure 5c). This reflects their similarity through a shared cytotoxic transcriptional program, with the top 4 genes *GZMA, CST7, CCL5, IFNG* being key marker genes marking the cytotoxic lymphocyte response expressed by both NK and effector/memory T cells [40]. *KLRG1* at rank 6 and *NKG7* at rank 8 are also known to mark both cell types [41, 42]. At *K* = 3, B cells merge with naive T cells. The top-scoring gene, *CCR7* reflects a shared naive/quiescent lymphocyte transcriptional state common to naive T and naive B cells [43] (Figure 5c). The top genes retain lineage-specific transcripts from both populations (e.g. *TRBV13* (rank 16) for T cells, *CR2* (rank 22) for B cells) amongst other non-canonical genes, and the DBI score is relatively high for this partition, indicating this merge reflects a partial, rather than fully coherent, shared signature.

At finer resolution up to *K* = 15, the procedure reveals biologically interpretable subtypes. A naive T-cell cluster present at broader clustering levels splits into two sub-clusters at *K* = 11. High *CD8A* (*s*_0,*d*_ = 0.23) and *CD8B* (*s*_0,*d*_ = 0.27) identifies cluster 0 as naive CD8^+^ T cells. Scores for these genes are *<* 0.01 in Cluster 4, while marker genes such as *FHIT* (*s*_4,*d*_ = 0.22) and *MAL* (*s*_4,*d*_ = 0.21) identify this population as naive CD4^+^ T cells. Both clusters score highly for *LEF1* and *LRRN3*, consistent with a shared regulatory programme across the CD4^+^/CD8^+^ lineage split [44].

Within the myeloid compartment at *K* = 8, the merge procedure distinguished three populations: classical monocytes (*S100A8, S100A9, LYZ*), non-classical monocytes and other DC cells (*CST3, HLA-DRA, LST1*), and plasmacytoid dendritic cells (pDCs) (*LILRA4, CLEC4C*) [36]. At *K* = 9 and finer resolution, the non-classical monocyte/DC cluster splits into CD16+ monocytes (Cluster 14; *MS4A7, s*_14,*d*_ = 0.46; *FCGR3A, s*_14,*d*_ = 0.44) and a DC population (Cluster 8; *HLA-DRA, s*_8,*d*_ = 0.54; *CST3, s*_8,*d*_ = 0.52) (Figure 5e); the structure at *K* = 9 achieves the lowest DBI, suggesting this split is biologically stable. This is further supported by the fixed-*K* benchmarking analysis earlier, where at *K* = 8, pDCs instead combined with other DC populations separate from a non-classical monocyte cluster which matched reference cell-type labels despite a distinct pDC signature being clearly recoverable at finer resolution (Supplementary Figure S24). This indicates that *K* = 8 cannot simultaneously capture both the pDC/DC and DC/CD16^+^ monocyte distinctions, and that *K* = 9 may better reflect the underlying myeloid substructure. A small platelet contaminated subset of 20 cells separates from the pDC population as Cluster 6 at *K* = 13, with genes such as *PPBP* (rank 6, *s*_6,*d*_ = 0.48) and *PF4* (rank 9, *s*_6,*d*_ = 0.47) likely reflecting platelet-associated ambient RNA or doublets [45]. The core pDC population (Cluster 11) is clearly defined by genes such as *LILRA*4, *IRF* 7, *SERPINF* 1.

Unlike Louvain and Leiden, which require tuning an arbitrary resolution parameter over a neighbourhood graph, our merge procedure is grounded directly in the probabilistic model: users specify only an initial, over-specified number of components *K*_init_, and merges are selected by the ICL of the resulting fit, so the hierarchy reflects structure in the latent space rather than parameter tuning. Because the procedure begins from an over-specified mixture, *K*_init_ need not reflect prior knowledge of the true number of cell types; in the supplementary material we show that initialising at *K*_init_ = 20 instead of 15 recovers a qualitatively similar hierarchy including the same split of the naive T cell cluster to CD4^+^ and CD8^+^ naive T cells. However, setting *K*_init_ far higher than the expected number of populations may increase the risk of over-fragmented, spurious components that do not correspond to real biological structure, and a longer, more fragmented merge path is more prone to suboptimal local ICL decisions. In practice, we recommend setting *K*_init_ to moderately exceed the anticipated number of cell types.

Applied to the Baron pancreas dataset, initialising at *K*_init_ = 15 (Supplementary Figure S26), we see broader cell types align with known pancreatic biology; at *K* = 3, cells split into a main beta cell cluster, other endocrine cells (alpha, delta, gamma, some beta), and a non-endocrine group [24]. Beta cells remain distinct at every resolution and sub-cluster at *K* = 6 and above, consistent with their status as the most abundant and transcriptionally distinct endocrine population [46]. Marker genes at *K* = 9 are consistent with the cell type structure we get when setting *K*_init_ = 9. Across 3 runs, the activated and quiescent stellate subpopulations - two phenotypes of the same lineage [47] - are separate at *K* = 15 but merge at *K* = 11 − 12. Further marker gene analysis is provided in the Supplementary Information.

## Discussion

As single-cell technologies develop and datasets grow in both scale and complexity, the methodological toolkit for their analysis has expanded rapidly. However, many existing approaches for cell-type identification still treat dimensionality reduction and clustering as separate steps, where the low-dimensionality embedding is often found without reference to the discrete cluster structure it is subsequently expected to reveal. Even though many recent approaches use deep-learning to learn complex structures, many of these sacrifice interpretability and rely on post-hoc methods for the annotation of cell types. Here, we introduce scFLAME, a probabilistic generative model that performs dimensionality reduction and clustering jointly within a unified framework, built upon a factor analytic framework with a shared gene-loading matrix across clusters.

Across both simulated and real datasets, scFLAME remains competitive with established clustering methods, while retaining an interpretable latent space in which up- and down-regulated genes for each cluster can be identified and related back to the underlying biology.

In particular, marker genes identified for clusters align well with canonical marker genes reported in the literature for corresponding cell types, as well as supporting novel insights, and are consistent with those identified by standard approaches. This identifiability extends to our hierarchical cell-typing procedure: at each level of the merging hierarchy, we are able to obtain a list of marker genes allowing coarser and finer partitions to be interpreted. Inspection of smaller sub-clusters is biologically informative; when a cluster is numerically dominated by one constituent cluster, its marker genes can obscure the existence of rarer, minority populations within the group. The resulting dendrogram reveals the hierarchical relationships underlying these sub-clusters, showing which broader cell-type family each sub-cluster belongs to, and at what level of resolution that relationship emerges. Few existing methods in the single-cell clustering literature explicitly surface this hierarchical structure; our greedy merging strategy provides one route to finding this, where we saw that finer and coarser cell types identified were in line with known cell-type biology. However, scFLAME still requires the user to specify the (maximum) number of mixture components *K* in advance; although the Dirichlet prior is designed to prune unsupported components automatically, we find this rarely occurs in practice, and instead rely on the post-hoc merging procedure to recover a suitable resolution. Extending the mixture component itself - for example via a nonparametric Bayesian formulation - may allow the number of cell types to be inferred more directly from the data.

scFLAME currently relies on full-batch variational inference, which produces high-fidelity latent representations, and we have shown it extends to datasets of hundreds of thousands of cells. However, the memory requirements present a computational bottleneck if atlas-scale datasets (millions of cells) are required to be analysed. A promising extension would adapt the model to a distributed stochastic variational inference framework, in which local, per-cell variational parameters are optimised asynchronously across multiple GPUs, while global parameters such as the shared loading matrix *L* are periodically synchronised across devices.

scFLAME could also be extended beyond transcriptomic clustering alone. The generative framework could be extended to spatial transcriptomic data, for example via a spatial smoothness prior over cluster assignments, or to multi-omic integration, incorporating paired modalities such as chromatin accessibility or protein abundance through additional shared or modality-specific loading structures.

Finally, semi-supervised methods have shown promise in dimensionality reduction and clustering tasks for single-cell data, allowing existing biological knowledge to guide the learned representation. While scFLAME performs strongly in a fully unsupervised scenario, augmenting scFLAME with prior information e.g. through a graph-based prior over genes or gene sets may improve its performance in scenarios where rich prior knowledge exists. Beyond improving cluster performance, this extension may also enrich the interpretability of the learned gene loadings, and could open a route towards modelling dependencies between genes within the latent structure.

## Methods

### The scFLAME model

We introduce scFLAME, a probabilistic model for single-cell RNA sequencing data that jointly performs dimensionality reduction and clustering while retaining interpretable gene programs through a shared loading matrix over the latent space. Let *X*_*ij*_ denote the observed count for gene *j* = 1, …, *D* in cell *i* = 1, …, *N* . We model counts *X*_*ij*_ using a negative binomial likelihood to account for overdispersion common in single-cell data.

We associate each cell *i* with a continuous latent factor *z*_*i*_ *∈* ℝ^*q*^ and a discrete latent cluster indicator *c*_*i*_ *∈* {1, …, *K*}, which are jointly updated in our inference scheme. Clustering is induced via a mixture prior over the latent factors, which represent cell positions in a low-dimensional embedded space; an optional batch coefficient *δ*_*b*_ allows the model to additionally correct for batch effects. The generative model is given by

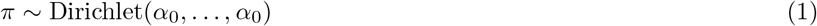

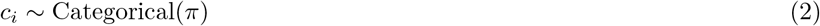

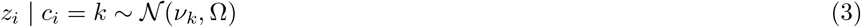

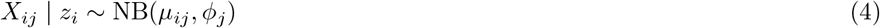

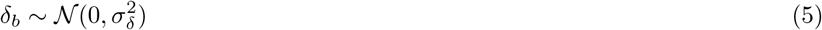

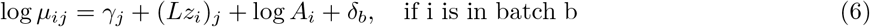

*π* = (*π*_1_, …, *π*_*K*_) are mixture weights with 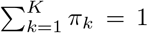 We place a Dirichlet prior on the mixture weights to encourage sparse cluster usage, allowing redundant components to be pruned during inference. *ν*_*k*_ *∈* R^*q*^ is the mean vector for cluster *k* in latent space, and Ω *∈* ℝ^*q*×*q*^ is a shared diagonal covariance matrix across all clusters. *L ∈* ℝ^*D*×*q*^ is a factor loading matrix representing gene loadings on the latent space, and *γ ∈* ℝ^*D*^ are gene-specific intercepts. *A*_*i*_ *>* 0 is a library-size offset which may be pre-estimated with TMM effective library sizes [48], and *ϕ*_*j*_ *>* 0 is a gene-specific dispersion parameter which may be pre-estimated using `edgeR` [49]. The library-size offset is fixed during training, while *ϕ*_*j*_ is updated as with other parameters (see ‘Initialisation and warm-up’). *δ*_*b*_ may be included in the log-mean to correct for batch effects, where cell *i* belonging to batch *b ∈* {0, …, *B* − 1} receives the corresponding offset *δ*_*b*_ *∈* ℝ^*D*^, with *δ*_0_ = 0 fixed for identifiability. A Gaussian shrinkage prior is placed on *δ*, equivalent under MAP estimation to *L*_2_ (ridge) regularisation, with smaller 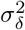 giving stronger shrinkage towards zero; we set 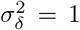 throughout, which performed well in practice.

We use the negative binomial distribution under (mean, dispersion) parameterisation. The log-probability mass function is given by:

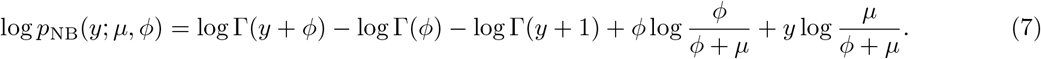

scFLAME generalises Negative Binomial Factor Analysis (NBFA), which provides a low-dimensional representation of each cell via latent factors *z*_*i*_ *∈* ℝ^*q*^ while modelling the overdispersed count likelihood. NBFA is a special case of our model where the latent factors follow a single Gaussian prior *z*_*i*_ *∼ N* (0, *I*_*q*_) rather than a mixture. Existing approaches such as ZINB-WaVE [18] and NewWave [50] perform similar, linearly interpretable dimensionality reduction under comparable likelihoods, followed by separate clustering in the latent space; scFLAME instead incorporates clustering directly through a mixture prior in a joint model. This formulation is conceptually related to models including VaDE [20], but with a linearly interpretable decoder (6) in place of neural encoder/decoder networks.

A shared diagonal covariance Ω between clusters improves stability and performance; cluster-specific covariance structures degrade performance on real single-cell data, consistent with overfitting or instability in variance estimation for small clusters. The isotropic prior in the NBFA latent space assumes unit variance and no correlation across dimensions in the latent space, and encourages approximately spherical embeddings. Consequently, clusters in this space are more likely to be of similar size and roughly spherical in shape. We exploit this architectural property by adopting an Equal Volume, Isotropic (EII) Gaussian Mixture Model hereafter referred to as EII GMM.

### Variational inference

Posterior inference over latent factors and cluster assignments is intractable due to the non-conjugate negative binomial likelihood. We therefore employ a mean-field variational approximation:

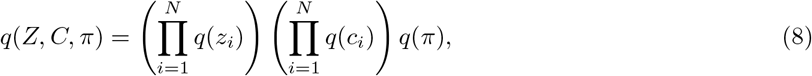

and we have that the variational families are given by:

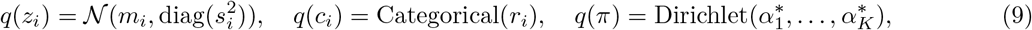

where *r*_*ik*_ = *q*(*c*_*i*_ = *k*). All remaining parameters are treated as model parameters and have no variational distribution associated. The evidence lower bound (ELBO) for scFLAME is given by:

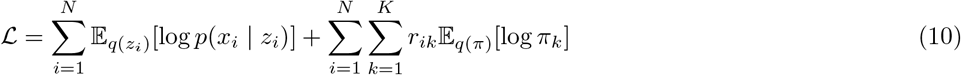

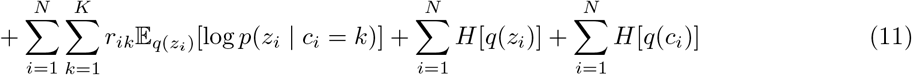

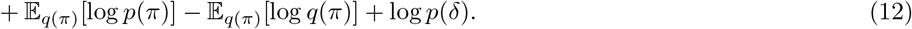

where *H*[·] denotes entropy. Parameters are updated via a coordinate ascent scheme: cluster assignment probabilities (known as ‘responsibilities’) *r*_*ik*_, cluster means/variances (*ν*_*k*_, Ω), and Dirichlet parameters 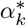 admit closed-form updates under the mean-field approximation. Responsibilities *r*_*ik*_ are computed via a temperature-annealed softmax to delay hard cluster assignments during early optimisation and reduce poor local optima. Parameters associated with the negative binomial likelihood - including variational parameters for the latent factors 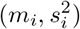, loading matrix *L* and dispersion parameters *ϕ*_*j*_, batch effects *δ* - are updated by stochastic gradient descent on the ELBO using Adam and the reparameterisation trick for expectations under *q*(*z*_*i*_). The parameter *γ* is held fixed in scFLAME to mitigate identifiability issues, as jointly updating *L* and *γ* while simultaneously updating the clustering can lead to redundant parameterisations that destabilise optimisation. The loading matrix is additionally regularised towards its initial value, and the batch-effect parameters are regularised through their zero-centred Gaussian prior. Further details of the variational updates, regularisation and annealing schedule are provided in the Supplementary Material.

### Initialisation and warm-up

scFLAME is initialised with a Negative Binomial Factor Analysis (NBFA) model to obtain initial low-dimensional embeddings *z*_*i*_, loading matrix *L* and gene-specific intercepts *γ*. This warm-up stage addresses an identifiability issue in early training: the model must simultaneously learn a low-dimensional representation of the count data and a Gaussian mixture structure defined on this latent space. As these components are mutually dependent, joint optimisation from random initialisation results in unstable optimisation as neither component provides useful signal to the other. To mitigate this, an NBFA model with a single Gaussian prior *z*_*i*_ *∼ N* (0, *I*) is fitted, and *L, γ*, {*m*_*i*_, *s*_*i*_} are optimised jointly. This yields a stable and informative initialisation for the full scFLAME model. If including batch effects, *δ* is optimised jointly with *L, γ*, {*m*_*i*_, *s*_*i*_} via gradient ascent on the ELBO and we do not include the Gaussian shrinkage prior for simplicity in the warm-up stage.

Library-size offsets *A*_*i*_ and initial dispersion parameters *ϕ*_*j*_ can optionally be pre-estimated prior to training using R, specifically via TMM effective library sizes [48] and `edgeR` tagwise dispersions, respectively. While pre-estimation is not strictly necessary - scFLAME can estimate both parameters end-to-end or rely on raw total UMI counts per cell — pre-estimating these values provides better empirical performance (Supplementary Material). We use `edgeR’s` tagwise dispersions directly, without inverting them to match the parameterisation in Equation (6): although this is the mathematically corresponding value, we found that it collapses the latent embedding on sparse data. Genes with many zero counts receive high `edgeR` dispersion, which maps to a near-zero dispersion under our parameterisation, flattening the likelihood surface with respect to the mean. Using uninverted estimates forces high-noise genes to be explained through the mean, driving better separation in the latent space. In the joint scFLAME clustering stage, *ϕ*_*j*_ is updated via gradient ascent, as the latent space is well-initialised so residual variance can be safely attributed to true dispersion to improve joint training.

The library-size offset *A*_*i*_ enters as a fixed additive term in log *µ*_*ij*_ (Equation (6)), ensuring differences in sequencing depth are absorbed directly into the mean rather than the latent representation. Generally, we set *A*_*i*_ to TMM-normalised effective library sizes, which are more robust to compositional biases than raw total counts [48]. For larger datasets, TMM estimation is often computationally prohibitive, so each cell’s raw total UMI count may be used as a substitute. Dispersion pre-estimation in `edgeR` can be run on a representative downsample of cells (e.g., 5000 cells).

All other variational and generative parameters were initialised via small-scale random draws (loadings, local means) or fixed constants (gene intercepts, local log-variances); full initialisation details are available in the accompanying code.

Variational inference remains a non-convex optimisation problem despite this warm-start initialisation, and typically converges to different local optima. We therefore perform *R* independent initialisations (using *R* = 5 in real-world simulations throughout this study) and retain the model maximising the final ELBO. The ELBO is a lower bound to the marginal log-likelihood and so serves as a proxy for model fit [51]. Many existing single-cell clustering frameworks, such as scDeepCluster and Louvain, lack a comparable formal objective for adjudicating between solutions.

Beyond the default EII GMM initialisation of the mixture prior, fitted to the empirical latent means *m* over 10 restarts and selecting the highest-likelihood configuration, which we use throughout the study, we also explore two random initialisations of the cluster means, assessing their effect on clustering robustness in the Supplementary Information.

## Implementation

scFLAME is implemented in PyTorch, with all variational parameters, cluster parameters, and gradient based updates computed on GPU to exploit batched tensor operations across cells and genes. All models were trained on a NVIDIA A100-SXM-80GB GPU (6,912 FP32 CUDA cores) with PyTorch version 2.4.1+cu121 and Python 3.11.13. Data preprocessing for library size and dispersion estimation was performed in R (version 4.6.0) using the `edgeR` package (version 4.10.1); pre-processed values were passed to scFLAME (in Python) via standard tabular files (e.g., CSV), avoiding external Python-R language bindings to simplify deployment. Stand-alone R scripts for the (optional) pre-processing step are provided.

We use 10 latent dimensions for simulations, and 15 for real data to capture the additional biological complexity. For NBFA, the learning rate for global parameters (*L, ϕ*_*j*_, *γ, δ*) is set at 10^−2^, with 10 Monte Carlo samples per gradient step, while local per-cell parameters 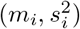 are updated at twice this rate, reflecting that per-cell embeddings should adapt more quickly to align with the current latent structure. For scFLAME, each epoch alternates (i) a closed-form E-step update of the latent posterior (3 gradient refinement steps at learning rate 10^−3^) and cluster responsibilities, (ii) closed-form M-step updates of the cluster means, variances, weights, and Dirichlet concentration parameters, and (iii) 3 gradient-ascent steps updating the factor loadings *L* and dispersions *ϕ* regularised with a penalty of strength 0.01 towards their initial values. For simulations, we run 350 epochs of NBFA warm-up and 250 epochs of scFLAME, while for real data, we run 500 epochs of NBFA warm-up and 300 epochs of scFLAME.

Gradient-based updates and updates for local variational parameters *m*_*i*_, *s*_*i*_ can be mini-batched for large *N* (in our experiments, datasets of *N >* 10000), whereas closed-form CAVI updates for cluster assignment remain cheap as they operate on the low-dimensional latent space and depend only on small *q, K*. Wall clock run-times can be seen in Supplementary Figure S8. 500 epochs of NBFA warm-up and 300 epochs of scFLAME clustering takes 9-10 seconds and 12-13 seconds respectively on a dataset of *N* = 1000, *D* = 4000, up to 10 minutes and 30 minutes respectively on a dataset of *N* = 100000, *D* = 4000.

### Identifying a tree of subtypes

scFLAME requires the user to specify a number of mixture components *K*. Under an overfitted Dirichlet prior, unsupported components are, in principle, asymptotically emptied by the posterior [52]. In practice, we find this rarely occurs due to the complexity of transcriptomic data coupled with the flexibility of the latent representations; specifying a larger *K* yields finer sub-clusters rather than automatic pruning of redundant components.

Following convergence of the full scFLAME model, we introduce a greedy agglomerative cluster merging procedure for discovering hierarchical structure in single-cell data. Starting from an over-specified mixture model, our procedure iteratively merges components, yielding a sequence of partitions across decreasing values of K. This process naturally defines a dendrogram over latent components. We can interpret the merge trajectory as a hierarchy of cell states, rather than relying on one optimal partition; cellular identities are often organised hierarchically without a single ‘true’ number of clusters. Merges of clusters have been seen in other variational inference frameworks; see [53] for an early proposal.

Latent means *m*_*i*_ and latent variances *s*_*i*_ are held fixed at their converged values throughout, so that merging operates entirely on the mixture structure without changing the learned cell representations. The EII GMM constraint is relaxed at this stage: rather than enforcing a single shared isotropic variance Ω across all components, we transition to a VII covariance structure in which each component is assigned its own scalar variance, 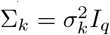,estimated via a closed-form variational M-step applied to the responsibilities *r*_*ik*_. At each step, all 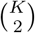 pairs of components are considered as merge candidates; to reduce computation when *K* is very large, candidates can optionally be pre-filtered to the top *M* pairs ranked by a similarity metric such as Bhattacharyya distance.

For a candidate merge of components *l* and *m* where we currently have *K* clusters in the model, the merged responsibilities are formed as 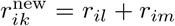, and the remaining *K* − 1 component parameters 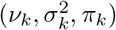 are re-estimated via the usual M-step on the updated responsibilities. The quality of the resulting (*K* − 1) component mixture is evaluated using the Integrated Completed Likelihood (ICL), with free parameter count (*K* − 1)(*q* + 1) +(*K* − 2) reflecting the VII structure. The pair whose merger yields the highest ICL is selected and applied. This is repeated greedily until *K* = 2 (or terminated earlier). We note that the ELBO usually drops as we merge clusters; forcing two clusters to share a single mean and covariance constrains the variational family, and increasing *K* improves the flexibility of the latent prior even when extra components do not correspond to distinct biological populations. A larger ELBO drop at a given merge may indicate that the collapsed components were more separated, reflecting a more costly loss of genuinely distinct structure. Alongside ICL, we compute the Davies-Bouldin Index (DBI) [54] at each step of the merge path, measuring the average ratio of within-cluster scatter to between-cluster separation, with lower values indicating more compact and well-separated clusters. Where ICL guides the merge decisions themselves, the DBI is used post-hoc to identify stable plateaus along the merge path - steps at which the cluster structure is particularly well-resolved - providing a complementary summary of the hierarchy and highlighting biologically meaningful resolutions.

### Interpretability of genes

The loading matrix *L ∈* ℝ^*D*×*q*^ provides a direct and interpretable link between the latent space in which clustering is performed and gene expression. For the purpose of cluster interpretation and gene scoring, we evaluate expression in the reference batch, for which the batch effect is defined to be zero, i.e., (*δ*_*b,j*_ = 0). Thus, the resulting scores characterise cluster-specific expression patterns independently of batch-specific shifts in gene expression. Since the generative model specifies

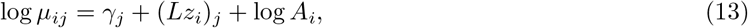

the expected log-expression of a cell in cluster *k*, relative to the population mean, is given analytically by

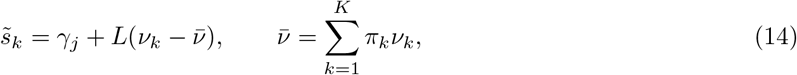

The *d*-th element of *s*_*k*_ gives the log fold-change in expected expression of gene *d* in cluster *k* relative to the population mean, propagated through the shared latent structure encoded in *L*. Since the batch effect is held fixed at its reference value, the contrast between a cluster and the population mean depends only on the shared latent expression structure.

To normalise for global abundance and penalise genes with broad, non-specific factor loading profiles, the final interpretability score *s*_*k,d*_ for the *d*-th gene in cluster *k* is scaled by the total absolute magnitude of its loadings across all *q* latent dimensions:

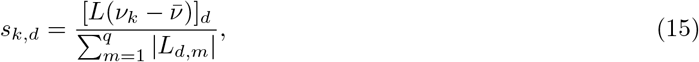

where 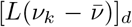 denotes the *d*-th element of the propagated latent contrast vector. This formulation prioritises genes whose loading weights are tightly concentrated on cluster-specific latent axes rather than distributed globally.

This quantity is a direct consequence of the generative model and requires no post-hoc statistical testing as *s*_*k*_ can easily be calculated from the final parameters of the fitted model, and allows our cluster interpretation to be joint with the generative model behind the clusters. As *L* is shared across all clusters, the scores can capture coordinated gene expression programs rather than gene-by-gene noise. Both upregulated (*s*_*kd*_ *>* 0) and downregulated (*s*_*kd*_ *<* 0) genes can be naturally identified through our model set-up, providing a complete picture of each cluster’s transcriptional identity. By evaluating the scores in the reference batch, cluster interpretation is disentangled from systematic batch-specific expression effects.

Genes can be ranked by |*s*_*kd*_| to produce a ranked list per cluster, with the sign of *s*_*kd*_ indicating direction of regulation. The ranked vectors *s*_*k*_ also serve as direct input to gene set enrichment analysis (GSEA), providing pathway-level interpretation of each cluster’s transcriptional programme. We perform this with the `fgsea` package [55]. Gene Ontology (GO) Biological Process over-representation analysis (ORA) of the top 50 positively scored marker genes per cluster was performed with `clusterProfiler` [56], using annotations from the Gene Ontology Consortium [57].

For pairwise comparison between clusters *k* and *j*, the score vector

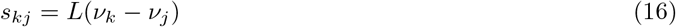

gives the log fold-change between the two cluster centroids.

### Competing methods

All methods were initialised with the true number of clusters (from simulated or ‘ground truth’ labels), and datasets filtered to the same number of highly variable genes as scFLAME, except where stated.

**NBFA + EII GMM** served as a baseline representing scFLAME’s initialisation quality. NBFA was run for 300 epochs and an EII GMM was fit to the latent factors.

**Louvain** is a graph-based community detection algorithm [16] and is the default clustering mechanism in the widely adopted Seurat [58] workflow. While newer iterations such as the Leiden algorithm improve on this [17], Louvain remains a foundational baseline. Cells are embedded via PCA on normalised data, a shared-nearest-neighbour graph is built and community detection takes place. Default settings were used for normalisation, variable feature selection, scaling, PCA, graph construction, and clustering resolution (0.8) initially. As Louvain does not allow direct specification of *K*, we also ran a ‘tuned *K*’ version where we ran a resolution sweep (0.1 to 2, in steps of 0.1 for simulated data, and 0.05 to 1, in steps of 0.05 for real data), selecting the resolution whose resulting number of clusters was closest to the known number of cell types *K*_true_ (stopping early on an exact match), for a comparable result to other methods at matched *K* (as ARI is biased to solutions with a similar *K*).

**PCA +** *k***-means** [14, 25]: raw counts were log(1 + *x*)-transformed, embedded with 20 principal components via prcomp in R, and clustered with *k*-means.

**MFA** (Mixture of Factor Analysers) [28] assumes that each observation arises from one of *K* latent components, each with its own low-rank factor structure. We implement a standard variational EM version of MFA (code provided) with a mix of closed-form updates and gradient-based updates (learning rate 0.01 for factor loadings and noise variances). We used library-size-normalised, log-transformed expression data, with 10 latent factors per cluster, initialised responsibilities with an even random split across clusters, and trained for 200 epochs.

**VaDE** [20] was implemented by adapting a third-party PyTorch reproduction of the original (Keras based) method (https://github.com/GuHY777/VaDE-pytorch), as the original code was outdated. VaDE combines a variational autoencoder with a GMM prior over the latent space, jointly learning a low-dimensional representation and cluster assignments by optimising a clustering-oriented ELBO. VaDE was designed for bounded continuous image pixel intensities; we replaced the sigmoid output and binary cross-entropy loss with a linear output and MSE loss appropriate for log-normalised expression data. Settings: batch size of 128, a fixed learning rate of 2e-4 (no decay schedule), 200 epochs, 10 latent dimensions and the same encoder/decoder layer sizes (500, 500, 2000).

**ZINB-WaVE** [18] fits a ZINB factor model to raw counts to obtain a low-dimensional representation, which allows for gene-level and cell-level covariates to be optionally included. We fit ZINB-WaVE (via the`zinbwave R` package) to the top 1000 highly variable genes, as in the original paper, as the package is computationally heavy. We used 20 latent dimensions, no covariates and set `epsilon` = 1000 (a regularisation parameter; package default is the number of genes). We applied *k*-means to the resulting latent space.

**scVI** [19] uses a variational autoencoder with a ZINB likelihood to learn a low-dimensional representation. We use 10 latent dimensions, as in the original paper, along with the default number of epochs (400), learning rate (0.001), and number and width of layers, and apply *k*-means to the resulting latent space. Training/validation ELBO tracking on a subset of simulation runs indicated convergence around epoch 200.

**LDVAE** [22] is a variant of scVI with a linearly interpretable decoder, enabling the identification of gene programs. We implement this via the `scVI-tools` package, with 10 latent dimensions as in the original paper, and run for 400 epochs with the same default learning rate, number of layers and layer width. ELBO tracking implied that this converged much faster than scVI.

**scDeepCluster** [26] combines a ZINB-based autoencoder with a deep embedded clustering objective, jointly learning a latent representation and cluster labels. We implemented scDeepCluster via the authors’ PyTorch implementation (https://github.com/ttgump/scDeepCluster_pytorch/tree/main); the original TensorFlow implementation was incompatible with our cluster. Pre-training was run for 600 epochs for both simulated and real data, while all other parameters followed the ‘Implementation’ section of the paper, including a bottleneck layer (latent space) of size 32.

**scDSC** [27] is a structural deep clustering network combining a ZINB-based autoencoder with a graph neural network module and a mutual-supervised module. We implemented scDSC via the authors’ repository (https://github.com/DHUDBlab/scDSC). In line with example code, we set encoder/decoder layer size as (500, 500, 2000) and 10 latent dimensions. We use the same batch size, epochs and learning rate for pre training as stated by the authors, and a learning rate of 0.0001 for the main training loop as in the example code. We run the main training for 200 epochs (full-batch, Adam).

### Evaluation metrics

Clustering results are evaluated against ‘true’ labels (either simulated or expert annotated) with three metrics - Adjusted Rand Index (ARI), Normalised Mutual Information (NMI), and Clustering Accuracy (ACC). Let *y*_*i*_ denote the true label of sample 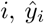 denote its predicted cluster assignment, and *n* be the total number of samples. Let *n*_*ck*_ denote the number of samples assigned to true class *c* and predicted cluster *k*, with row and column sums: 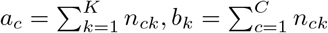

The Rand Index (RI) [59] calculates the proportion of pairs of objects that are assigned to either the same or different clusters in both partitions. The ARI [60] adjusts the RI for chance:

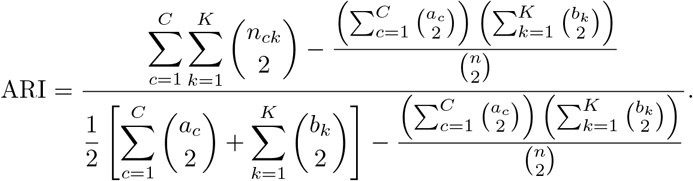

An ARI of 1 indicates identical cluster assignments; ARI close to 0 indicates that the clustering is no better than random assignment. ARI values can also be negative.

NMI [61] takes the mutual information between the two cluster partitions and normalises this by dividing by the Shannon entropy of the clusterings. An NMI of 1 indicates perfect clustering, while 0 means there is no mutual information between the two clusterings:

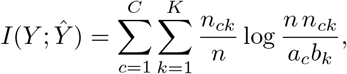

and

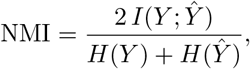

where 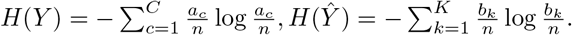.

ACC measures the proportion of correctly assigned samples after using the Hungarian algorithm [62] to optimally match clustering labels:

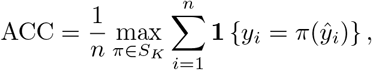

where *π* is the optimal label permutation, given by the Hungarian algorithm. ACC ranges from 0 to 1, where a higher value indicates better clustering performance.

To assess the separation of cell types in a given low-dimensional representation to evaluate batch correction in the Pancreas dataset, we computed the silhouette width of each cell using the `cluster R` package [35], treating reference cell-type labels as cluster assignments. For a cell *i*, let *a*(*i*) denote the mean distance to all other cells of the same annotated type, and *b*(*i*) the mean distance to cells in the nearest neighbouring type (the type with the lowest mean distance to *i*). The silhouette width is given by 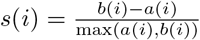 and ranges -1 to 1, with larger values indicating that a cell lies closer to cells of its own annotated type than to any other type. We report the mean silhouette width per cell type, computed directly on Euclidean distances within each method’s latent or embedding space (scFLAME latents, Harmony embeddings).

### Data simulation

For the simulation experiments, we simulated data directly from the scFLAME generative model (‘The scFLAME model’). Two scenarios of varying difficulty were constructed by increasing the number of clusters *K* and latent dimensions *q*, while decreasing cluster separation and the dispersion parameter *r* (higher overdispersion). Latent factors *z*_*i*_ were sampled from cluster-specific Gaussian distributions *N* (*ν*_*k*_, 0.8^2^*I*), where cluster centres *ν*_*k*_ were spaced according to a separation parameter. Observed counts *X*_*ij*_ were generated using a Negative Binomial model via the Gamma-Poisson parameterisation. The ‘Easy’ scenario used (*N* = 800, *D* = 1000, *r* = 2.5, sep = 2.5) and ‘Hard’ used (*N* = 1500, *D* = 2000, *r* = 0.5, sep = 1.5). Each scenario was evaluated across three independent replicates across 20 simulated datasets.

We also simulated data from Splat [23] and ZINBWaVE [18], two well-established data-generation methods for single-cell data from the literature [63]. We derived parameters from the first donor of the Baron pancreas dataset [24] (inDrop). When examining different levels of dropout and clustering signal strength, we derived parameters from the Segerstolpe pancreas dataset [29] (Smart-Seq2). As before, we simulated 20 datasets for all scenarios and evaluated each clustering method three times per dataset.

For Splat (`Splatter R` package), cell types with *<* 50 cells (*<* 2% of data) were removed, genes with *<* 10 total counts removed, and the top 5000 highly variable genes (HVGs) selected. Global parameters were estimated with `splatEstimate`, and 1,000 cells across 4,000 genes were simulated with `splatSimulate`, setting `group.prob = c(0.4, 0.25, 0.15, 0.1, 0.1)`, `de.prob = c(0.3, 0.2, 0.1, 0.2, 0.2)` and `de.facLoc = 0.2` to simulate clusters. Dropout was varied via `dropout.mid` between val-ues (1, 2, 3, 4) corresponding to dropout rates 19.8 ± 0.2%, 32.4 ± 0.2%, 47.8 ± 0.2%, 63.6 ± 0.2% (default 0).

Clustering signal strength was varied with `de.facScale` between values [0.2, 0.25, 0.3, 0.35] (default 0.4).

Dropout rate, *d*, was calculated as:

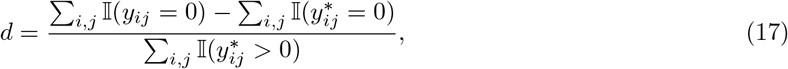

where *y*_*ij*_ represents the observed simulated counts for gene *j* in cell *i*, and 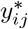 represents the latent ground-truth counts prior to dropout.

For ZINB-WaVE (`zinbwave R` package), we filtered similarly to Splat except used 2,000 HVGs due to the higher computational cost. The design used intercept and cell-type labels in *X* to model the original group structure, and an intercept-only gene matrix *V* (no gene-level covariates). We included a library-size offset for the mean model and *K* = 0 latent factors. `zinbFit` was run with common dispersion and zero inflation both set to true. Synthetic counts were sampled using `zinbSim`, preserving original dimensions and group labels.

For Splat and ZINB-WaVE datasets, to facilitate downstream benchmarking with scFLAME and NBFA+EII GMM, TMM effective library sizes and edgeR dispersion estimates were calculated as in ‘The scFLAME model’. We filtered all datasets to the top 3000 HVGs for benchmarking, except ZINB-WaVE where we used the top 1000 due to computational cost. Further simulation scenarios and data generation details are provided in the Supplementary Information.

### Real-world data and preprocessing

For the eight real-data experiments without batch correction, all datasets were filtered to 5000 HVGs, except the mouse spleen datasets, which were filtered to 8000 HVGs.

The **PBMC 4K** dataset was obtained from 10X Genomics (https://www.10xgenomics.com/datasets/fresh-68-k-pbm-cs-donor-a-1-standard-1-1-0) as a filtered UMI count matrix generated using the Chromium Single Cell 3’ platform [30]. We compared to authors’ default cluster labels (graph-based clustering on PCA-reduced data). After removing cells with *>* 5% mitochondrial RNA content, we removed genes expressed in *<* 3 cells and cells expressing *<* 200 genes.

The **Baron pancreas** dataset, consisting of pancreatic islet cells, was subset to a single human donor (GSM2230757) [24]. The dataset was accessed through the `scRNAseq` Bioconductor package. Cell-type labels provided by the original authors were derived from recursive hierarchical clustering with Ward’s criterion on correlations of highly variable/highly expressed genes, followed by manual merging of over-split clusters. Cells were filtered via `scater::quickPerCellQC` on library size and detected features. Genes expressed in *<* 10 cells and cell types comprising *<* 2% of all cells were excluded.

The **Zeisel mouse brain cortex** STRT-Seq dataset consists of 3005 cells from the primary somatosensory cortex and hippocampal CA1 region, where cells were isolated using the Fluidigm C1 microfluidics system followed by Illumina sequencing (UMI-based) [31]. The dataset was accessed through the scRNAseq Bioconductor package. We benchmarked against the authors’ broad `level1class` annotations (derived via BackSPIN recursive biclustering, merged from 77 fine clusters): interneurons, pyramidal SS neurons, pyramidal CA1 neurons, oligodendrocytes, microglia, endothelial-mural cells, and astrocytes-ependymal cells. Cells were filtered via `scater::quickPerCellQC` on library size, detected genes, ERCC spike-in proportion, and mitochondrial transcript proportion following [64].

The **mouse uterus** dataset of the Mouse Cell Atlas project [2] was obtained via https://figshare.com/s/865e694ad06d5857db4b. We downloaded the digital expression matrix (batch gene background removed) of all 400,000 single cells and filtered to uterus cells. Cell types were manually labelled by the original authors via Seurat-based clustering [58], first at atlas-scale followed by organ-scale, using differentially expressed markers. After removing cells with *>* 5% mitochondrial RNA content, we removed cells expressing *<* 50 genes, genes expressed in *<* 50 cells and cell types making up *<* 3% of total cells, following [65]. This dataset is made up of two batches with filtered sizes 1954 and 1216. Of the 12 annotated cell types, 10 are strongly batch-skewed (*>* 80% of cells from a single batch), leaving little shared biology across the two batches to anchor a correction against; applying correction may lead to conflating batch and cell-type signal. Given this strong confounding between batch and biology, we evaluated this dataset without batch correction in the main text.

The **Segerstolpe pancreas** dataset consists of 2133 human pancreas endocrine and exocrine cells from 7 donors, sequenced by Smart-Seq2 on an Illumina HiSeq 2000 [29] and was accessed via the `scRNAseq` Bioconductor package. Cell-type annotations were obtained by identifying visually distinct groups in a tSNE embedding of HVGs and assigning biological identities based on marker genes. This revealed major populations; a cluster of 1554 endocrine cells were isolated and independently projected using a new set of HVGs to resolve endocrine subtypes. Following the OSCA quality-control workflow [64], we removed cells previously flagged as low quality by the original authors, then applied a further per-donor outlier-based QC step (`scater::quickPerCellQC`). Donors H5, H6 were excluded from the outlier-detection step due to substantially different quality-metric distributions. Cell types representing *<* 2.5% of the dataset after filtering were excluded.

The **TM mouse spleen** datasets were obtained from the `TabulaMurisSenisData` Bioconductor package, which provides the Tabula Muris Senis atlas of mice single-cell transcriptomes [66]. To align with the original Tabula Muris atlas, we filtered to 3-month old mice [32] and spleen cells. We analysed the Smart-seq2 and 10x Chromium datasets separately. Reference cell-type annotations were derived by the authors for each organ via Louvain clustering in Seurat v2.2.1 and annotated via differentially expressed genes; mixed clusters were iteratively refined by increasing the clustering resolution or reclustering subsets of cells. We compared to the ‘Cell Ontology’ annotations; multiple computational clusters were often represented by a single Cell Ontology class. Cells with *>* 5% mitochondrial RNA content were removed. For the Smart-seq2 dataset, cells expressing *>* 500 genes were retained; for the 10X Genomics dataset, cells expressing *>* 250 genes were retained to account for the lower sequencing depth. Genes detected in *>* 3 cells were retained; cell populations representing *<* 1% of total cells after filtering were excluded.

The **PBMC 68K** dataset was obtained from 10X Genomics (https://www.10xgenomics.com/datasets/fresh-68-k-pbm-cs-donor-a-1-standard-1-1-0) as a filtered UMI count matrix generated using the Chromium Single Cell 3’ platform [30]. We reproduced and compared to the authors’ cluster labels analysed in the paper, derived from *k*−means clustering (*k* = 10) run on the top 50 principal components from the 1000 top HVGs (code is available at http://www.github.com/10XGenomics/single-cell-3prime-paper). After removing cells with *>* 5% mitochondrial RNA content, we removed genes expressed in *<* 3 cells and cells expressing *<* 200 genes.

For the two datasets used to evaluate scFLAME’s batch-correction term, preprocessing generally followed Song, Chan, and Wei [13], using all genes retained after pre-processing.

The **LUAD** dataset (GEO accession number GSE118767) was obtained from Tian et al. [34], comprising three lung adenocarcinoma cell lines (HCC827, H1975, H2228), where all cell lines mixed and separated into three batches which were profiled on a different protocol (CELseq2, 10x Chromium, Drop-seq), yielding 1401 cells with known ground-truth cell-line identity. Following the preprocessing approach of [13, 67], the raw count data was obtained from https://github.com/LuyiTian/sc_mixology. Each protocol’s dataset was independently normalised using `scran`, followed by `logNormCounts`), and the top 6,000 highly variable genes were selected per protocol using `modelGeneVar`. The intersection of highly variable genes across all three protocols (2344 genes) was retained, and raw counts restricted to this common gene set were combined across protocols to form the final dataset, with protocol identity retained as the batch label.

The **Pancreas** dataset combines four publicly available human pancreas scRNA-seq studies, profiled by CEL-seq (GSE81076) [68], CEL-seq2 (GSE85241) [69], Smart-seq2 (GSE86473) [70], and Smart-seq2 (E-MTAB-5061) [29]. Our processing broadly follows the approach of Haghverdi et al. [67] and Song, Wang, and Dunson [71], broadly using code from https://github.com/MarioniLab/MNN2017/blob/master/Pancreas with several modifications described below. After removing genes expressed in *>* 90% of cells as zero and cells with *>* 80% zero expression, size factors were estimated per dataset using `scran’s` deconvolution method (`quickCluster/computeSumFactors`, followed by `logNormCounts`); this differs from the original pipeline of Haghverdi et al. [67], whose spike-in- and quality-control-aware implementation relies on `scran/scater` functions since deprecated, preventing an exact reproduction. Highly variable genes were identified independently within each dataset and combined across studies via meta-analysis of per-gene *p*-values (geometric mean of — log_10_(*p*) across datasets, combined threshold ≤ 0.001). Cell-type labels for GSE86473 and E-MTAB-5061 were taken directly from the original publications’ metadata. For GSE81076 and GSE85241, which lack pre-assigned labels, we deviated from the original PAM-on-all-HVGs approach of Haghverdi et al. [67]: using these *k*-medoids clusters did not correspond well to the visual structure of the t-SNE embedding or to marker gene expression, so cell types were instead assigned via PAM clustering (*K* = 9) restricted to a panel of established pancreatic marker genes (GCG for alpha, INS for beta, SST for delta, PPY for gamma, PRSS1 for acinar, KRT19 for ductal, COL1A1 for mesenchymal cells), with each resulting cluster labelled by its corresponding highly expressed marker. We took the intersection of genes across all 4 datasets as the genes in the final raw counts matrix, giving us 6630 genes profiled across 7099 cells.

## Code availability

Code for running scFLAME is available at https://github.com/j-ackierao/scFLAME_code.

