## Supplementary Information for "scFLAME: a unified generative model for interpretable clustering, hierarchical structure discovery and marker-gene identification in single-cell RNA-seq data"

#### 1 Comparison of related methods

Table S1: Representative methods for single-cell dimensionality reduction and/or clustering, classified by whether they model raw counts, jointly optimise clustering with the embedding, and yield interpretable gene programmes.

| Method | Class | Count based | Joint clustering | Interpretable gene programmes |
|---|---|---|---|---|
| PCA + k-means | Linear DR + clustering | No | No | Partial |
| Louvain/Leiden (Seurat) | Graph clustering | No | No | No |
| SIMLR/MPSSC | Multi-kernel similarity | No | Yes | No |
| ZIFA | ZI factor analysis | Yes | No | Partial |
| ZINB-WaVE/NewWave | NB factor model | Yes | No | Partial |
| scVI | VAE | Yes | No | No |
| LDVAE | Linear-decoder VAE | Yes | No | Yes |
| scDeepCluster/scDCC | Deep embedded clustering | Yes | Yes | No |
| scSMD | Attention autoencoder | Yes | Yes | Partial |
| VaDE | GMM-prior VAE | No | Yes | No |
| DR-SC | HMRF joint model | No | Yes | No |
| scDSC/scGNN/scGAC/scHNTL | Graph neural network | Yes | Yes | No |
| **scFLAME (this work)** | NB factor + GMM prior | Yes | Yes | Yes |

### 2 Supplementary Methods

#### 2.1 Variational Updates: detailed

We assume a mean-field variational family:

$$q(z_i, c_i, \pi) = q(z_i)\, q(c_i)\, q(\pi) = \mathcal{N}(m_i, \mathrm{diag}(s_i^2)) \times \mathrm{Categorical}(r_i) \times \mathrm{Dirichlet}(\alpha_1^*, \ldots, \alpha_K^*) \tag{1}$$

where $q(z_i) = \mathcal{N}(m_i, \mathrm{diag}(s_i^2))$, $q(c_i)$ is categorical with parameters $r_i$, and $q(\pi)$ is Dirichlet.

##### 2.1.1 E-step (latent factors): Update $m_i$, $s_i$

For each cell $i$, we optimise the variational parameters $m_i$ and $s_i$ by gradient ascent on the ELBO. The relevant terms are:

$$\mathcal{L}_i(m_i, s_i) = \mathbb{E}_{q(z_i)}[\log p(x_i \mid z_i)] + \sum_{k=1}^{K} r_{ik} \mathbb{E}_{q(z_i)}[\log p(z_i \mid c_i = k)] + H[q(z_i)]. \tag{2}$$

The expected log-likelihood is approximated using the reparameterisation trick. We sample:

$$z_i^{(s)} = m_i + s_i \odot \varepsilon_i^{(s)}, \quad \varepsilon_i^{(s)} \sim \mathcal{N}(0, I_q), \quad s = 1, \ldots, S, \tag{3}$$

and approximate:

$$\mathbb{E}_{q(z_i)}[\log p(x_i \mid z_i)] \approx \frac{1}{S} \sum_{s=1}^{S} \sum_{j=1}^{D} \log p_{\mathrm{NB}}(X_{ij}; \mu_{ij}(z_i^{(s)}), \phi_j), \tag{4}$$

where $\log \mu_{ij}(z_i^{(s)}) = \gamma_j + (L z_i^{(s)})_j + \log A_i$.

We use stochastic gradient ascent with the Adam optimiser alongside the reparameterisation trick to update $m_i$ and $\log s_i$ for a fixed number of steps (typically 3-5 steps per E-step iteration). Gradients are clipped to prevent instability.

##### 2.1.2 E-step (responsibilities): Update $r_{ik}$

Given the current values of $m_i$, $s_i$, $\nu_k$, $\Omega$, and $\pi_k$, we update the responsibilities $r_{ik} = q(c_i = k)$ using:

$$r_{ik} \propto \exp\left(\mathbb{E}_{q(\pi)}[\log \pi_k] + \mathbb{E}_{q(z_i)}[\log p(z_i \mid c_i = k)]\right), \tag{5}$$

normalised so that $\sum_{k=1}^{K} r_{ik} = 1$. Explicitly:

$$\log r_{ik} = \log \pi_k - \frac{1}{2} \sum_{j=1}^{q} \left( \log(2\pi\omega_j^2) + \frac{(m_{ij} - \nu_{kj})^2 + s_{ij}^2}{\omega_j^2} \right) + \mathrm{const.} \tag{6}$$

These responsibilities correspond to soft cluster allocations for the latent representations. To improve convergence and avoid local optima, we apply temperature annealing during the early phases of training. We modify the responsibility update to:

$$r_{ik} \propto \exp\left( \frac{1}{T} \left[ \log \pi_k + \mathbb{E}_{q(z_i)}[\log p(z_i \mid c_i = k)] \right] \right), \tag{7}$$

where $T > 1$ initially and linearly decreases to $T = 1$ for a given number of training epochs. Higher temperatures encourage more exploratory assignments early in training. Throughout our experiments, we set $T = 1.5$, and anneal for $epochs/3$.

##### 2.1.3 M-step (cluster parameters): Update $\nu_k$, $\Omega$, $q(\pi)$

Given the current responsibilities $r_{ik}$ and variational posteriors $q(z_i)$, we update the cluster parameters in closed form. These updates have closed-form solutions due to the conjugacy between the Gaussian (mixture) prior and the Gaussian variational posterior, and the conjugacy between the Dirichlet prior on the weights and the categorical variational distribution for cluster allocations $c_i$.

**Mixture weights (Dirichlet prior):** We place a symmetric Dirichlet prior on the mixture weights:

$$\pi \sim \mathrm{Dir}(\alpha_0, \dots, \alpha_0), \tag{8}$$

where we typically have $\alpha_0 < 1$, encouraging sparse solutions with small redundant clusters emptied. When $\alpha_0 < 1$, the Dirichlet prior concentrates mass near the boundary of the simplex, encouraging solutions in which many mixture components have negligible weight.

The relevant ELBO terms are:

$$\mathcal{L}_\pi = \sum_{i=1}^{N}\sum_{k=1}^{K} r_{ik} \log \pi_k + \sum_{k=1}^{K} (\alpha_k - 1) \log \pi_k. \tag{9}$$

Under the mean-field approximation, the optimal variational factor for $\pi$ is Dirichlet:

$$q^*(\pi) = \mathrm{Dirichlet}(\alpha_1^*, ..., \alpha_K^*) \tag{10}$$

where for $k = 1, ..., K$,

$$\alpha_k^* = \alpha_k + \sum_{i=1}^{N} r_{ik} \tag{11}$$

**Cluster means:** For each cluster $k$, we maximise the expected log-prior with respect to $\nu_k$. The relevant ELBO terms involving $\nu_k$ are:

$$\mathcal{L}_{\nu_k} = \sum_{i=1}^{N} r_{ik} \mathbb{E}_{q(z_i)}[\log p(z_i \mid c_i = k)]. \tag{12}$$

Under the Gaussian prior $p(z_i \mid c_i = k) = \mathcal{N}(\nu_k, \Omega)$ with $\Omega = \mathrm{diag}(\omega_1^2, \dots, \omega_q^2)$, this becomes:

$$\mathcal{L}_{\nu_k} = \sum_{i=1}^{N} r_{ik} \mathbb{E}_{q(z_i)} \left[ -\frac{1}{2} \sum_{j=1}^{q} \left( \log(2\pi\omega_j^2) + \frac{(z_{ij} - \nu_{kj})^2}{\omega_j^2} \right) \right] \tag{13}$$

$$= -\frac{1}{2} \sum_{i=1}^{N} r_{ik} \sum_{j=1}^{q} \frac{(m_{ij} - \nu_{kj})^2 + s_{ij}^2}{\omega_j^2} + \text{const.} \tag{14}$$

Taking the derivative with respect to $\nu_{kj}$ and setting to zero:

$$\frac{\partial \mathcal{L}_{\nu_k}}{\partial \nu_{kj}} = \sum_{i=1}^{N} r_{ik} \frac{m_{ij} - \nu_{kj}}{\omega_j^2} = 0 \tag{15}$$

$$\nu_{kj} = \frac{\sum_{i=1}^{N} r_{ik} m_{ij}}{\sum_{i=1}^{N} r_{ik}}. \tag{16}$$

In vector form:

$$\nu_k = \frac{\sum_{i=1}^{N} r_{ik} m_i}{N_k}. \tag{17}$$

This is equivalent to a responsibility-weighted mean of the variational means $m_i$ for cluster $k$.

**Shared diagonal covariance:** The shared diagonal covariance is updated by pooling information across all clusters:

The relevant ELBO terms involving $\Omega$ are:

$$\mathcal{L}_\Omega = \sum_{i=1}^{N}\sum_{k=1}^{K} r_{ik} \mathbb{E}_{q(z_i)}[\log p(z_i \mid c_i = k)]. \tag{18}$$

Expanding the Gaussian log-density:

$$\mathcal{L}_\Omega = \sum_{i=1}^{N}\sum_{k=1}^{K} r_{ik}\left[-\frac{1}{2}\sum_{j=1}^{q}\left(\log(2\pi\omega_j^2) + \frac{(m_{ij}-\nu_{kj})^2 + s_{ij}^2}{\omega_j^2}\right)\right] \tag{19}$$

$$= -\frac{1}{2}\sum_{j=1}^{q}\left[N\log(2\pi\omega_j^2) + \frac{1}{\omega_j^2}\sum_{i=1}^{N}\sum_{k=1}^{K} r_{ik}\left((m_{ij}-\nu_{kj})^2 + s_{ij}^2\right)\right]. \tag{20}$$

Taking the derivative with respect to $\omega_j^2$ and setting to zero:

$$\frac{\partial \mathcal{L}_\Omega}{\partial \omega_j^2} = -\frac{1}{2}\left[\frac{N}{\omega_j^2} - \frac{1}{(\omega_j^2)^2}\sum_{i=1}^{N}\sum_{k=1}^{K} r_{ik}\left((m_{ij}-\nu_{kj})^2 + s_{ij}^2\right)\right] = 0 \tag{21}$$

$$\omega_j^2 = \frac{1}{N}\sum_{i=1}^{N}\sum_{k=1}^{K} r_{ik}\left((m_{ij}-\nu_{kj})^2 + s_{ij}^2\right). \tag{22}$$

Note that the sum $\sum_{k=1}^{K} r_{ik} = 1$ for each $i$, so we can write:

$$\omega_j^2 = \frac{1}{N}\sum_{i=1}^{N}\left[\sum_{k=1}^{K} r_{ik}(m_{ij}-\nu_{kj})^2 + s_{ij}^2\right]. \tag{23}$$

This update has an intuitive interpretation: for each latent dimension $j$, we compute the responsibility-weighted squared distance from each cell's variational mean $m_{ij}$ to each cluster centre $\nu_{kj}$, plus the variational uncertainty $s_{ij}^2$, and average over all cells. Because the same variance $\omega_j^2$ is shared across all clusters, the estimate pools information from the entire dataset. We clamp the variance estimates to a minimum value to prevent numerical instability.

##### 2.1.4 Update $L, \phi, \delta$

We update $L$, $\phi$, and $\delta$ via stochastic gradient ascent on the ELBO, as there are no closed-form updates available for these parameters. All other parameters are held fixed. Gradients are estimated using Monte Carlo samples and the reparameterisation trick, analogously to the latent factor updates. During training, we keep $\gamma$ fixed to mitigate identifiability issues. Jointly updating $L$ and $\gamma$ while simultaneously updating the clustering can lead to redundant parameterisations that yield equivalent likelihoods, which in practice can destabilise optimisation; see 'Initialisation and warm-up' in the main text.

To stabilise estimation of the loading matrix, we introduce a quadratic regularisation term that penalises deviations from its initial value $L_{\text{init}}$:

$$\mathcal{R}_L = L_{\text{reg}}\,\|L - L_{\text{init}}\|_F^2\,, \tag{24}$$

where $L_{\text{reg}} > 0$ controls the strength of the regularisation and $\|\cdot\|_F$ denotes the Frobenius norm.

As we impose the zero-centred Gaussian prior on the batch offsets, this contributes the following negative log-prior to the objective:

$$\mathcal{R}_\delta = \frac{1}{2\sigma_\delta^2}\,\|\delta_{\text{free}}\|_2^2\,. \tag{25}$$

, where $\delta_{\text{free}}$ represents the unconstrained batch coefficients, with $\delta_0 = 0$ for identifiability. Thus, the gradient-based parameters are optimised using the ELBO together with the loading-matrix regularisation term, with the Gaussian prior on $\delta$ contributing directly through its log-prior term.

#### 2.2 Data preprocessing: cell type mapping

The PBMC 4K dataset was provided with graph-based clustering assignments represented as numerical cluster labels (via the 10X Genomics website), without corresponding predefined cell-type annotations by the authors. We therefore used these numerical labels directly to represent the original data-driven cluster structure.

Table S2: Mapping of numerical cluster identifiers to reference cell-type labels across datasets. Overview of numerical cluster IDs used in visualisation plots and their corresponding published reference cell-type annotations for each dataset.

| **Dataset** | **Cluster ID** | **Reference cell-type label** |
|---|---|---|
| Zeisel | 1 | Astrocytes-ependymal |
| | 2 | Endothelial-mural |
| | 3 | Interneurons |
| | 4 | Microglia |
| | 5 | Oligodendrocytes |
| | 6 | Pyramidal CA1 |
| | 7 | Pyramidal SS |
| Baron | 1 | Acinar |
| | 2 | Activated stellate |
| | 3 | Alpha |
| | 4 | Beta |
| | 5 | Delta |
| | 6 | Ductal |
| | 7 | Endothelial |
| | 8 | Quiescent stellate |
| | 9 | Gamma |
| Uterus | 1 | Stromal cell Hsd11b2 high |
| | 2 | Dendritic cell |
| | 3 | Muscle cell Mgp high |
| | 4 | Stromal cell Ccl11 |
| | 5 | Stromal cell Cxcl14 |
| | 6 | Stromal cell Has1 |
| | 7 | Macrophage |
| | 8 | Endothelial Tm4sf1 |
| | 9 | Glandular epithelium Sprr2f |
| | 10 | NK cell |
| Segerstolpe | 1 | Acinar |
| | 2 | Alpha |
| | 3 | Beta |
| | 4 | Delta |
| | 5 | Ductal |
| | 6 | Gamma |
| | 7 | PSC |
| TM Spleen 10X | 1 | B cell |
| | 2 | Macrophage |
| | 3 | Mature NK T cell |
| | 4 | NK cell |
| | 5 | Proerythroblast |
| | 6 | T cell |
| TM Spleen SS2 | 1 | B cell |
| | 2 | $CD4^{+}$ alpha-beta T cell |
| | 3 | $CD8^{+}$ alpha-beta T cell |
| | 4 | NK cell |
| | 5 | Proerythroblast |

### 3 Generating single-cell data with scFLAME

An advantage of the probabilistic framework behind scFLAME is that it serves as a fully generative model, allowing us to simulate realistic *in silico* single-cell RNA-sequencing (scRNA-seq) datasets using parameters fitted from real datasets. Simulations of scRNA-seq data are valuable for benchmarking computational tools where experimental ground truth is unattainable; we saw that simulations from Splat and ZINBWaVE were useful in our assessment of scFLAME versus other methods for clustering count data. To assess whether scFLAME captures the statistical structure of real transcriptomic data, we simulated new datasets from parameters fitted to the Baron pancreas dataset. After filtering to the top 5000 HVGs, and running scFLAME with fixed cluster allocations (as per our known 'true' labels), we could sample cluster assignments from the learned Dirichlet posterior and latent states $z$ directly from the learned Gaussian mixture prior given these cluster assignments ($z \sim \mathcal{N}(\nu_k, \Omega)$). This then allowed us to sample gene expression counts from the learned generative model, analogous to simulation frameworks such as ZINB-WaVE and Splatter.

$$q(z_i) = \mathcal{N}(m_i, \mathrm{diag}(s_i^2)), \quad q(c_i) = \mathrm{Categorical}(r_i), \quad q(\pi) = \mathrm{Dirichlet}(\alpha_1^*, \ldots, \alpha_K^*), \tag{26}$$

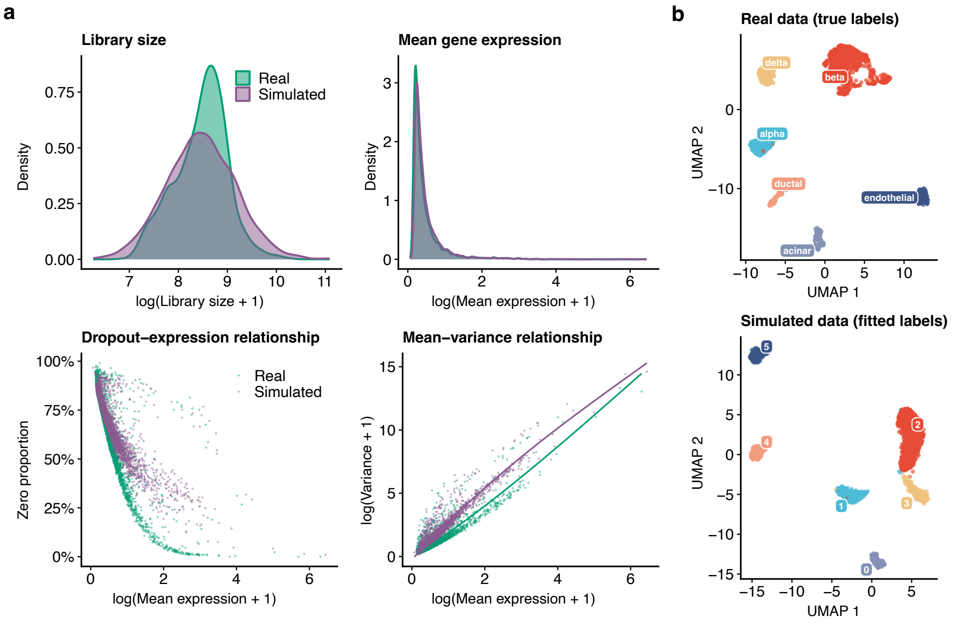


Figure S1: Evaluation of one example of scFLAME simulated data against Baron pancreas dataset. **a,** Comparison of common quality control metrics between real (green) and simulated (purple) datasets. Metrics include the probability density distributions of library sizes per cell (top left) and mean gene expression levels per gene (top right). Bottom panels display the dropout-expression relationship (bottom left) and the feature-level mean-variance relationship with trend-lines (bottom right). **b,** UMAP embeddings of the real Baron pancreas data annotated by true biological cell-types, and the simulated dataset annotated by predicted labels by running scFLAME on the simulated data.

We see in Figure S1 that scFLAME successfully replicates some key statistical features of single-cell data. While the simulated library size shows a slight shift compared to the real data, the mean gene expression profile is captured well. The model also generates a similar proportion of structural zeros observed in the real data across varying expression levels, suggesting that the negative binomial parameterisation can successfully account for single-cell sparsity without any need for explicit modelling of zero-inflation. This behaviour was consistent across all 3 simulated datasets, where we saw zero proportions of 0.715, 0.712, and 0.714, which was closely aligned with the real data zero proportion of 0.695. This supports the use of the NB model in our framework. The dropout-expression relationship is relatively well-replicated, with simulated data producing a slightly higher proportion of zeroes as mean expression increased.

However, the mean-variance relationship is less faithfully reproduced compared to other dedicated simulation frameworks such as Splat. We attribute this to two parts of our model: firstly, sampling latent factors from the variational posterior $q(z_i)$ rather than the true underlying posterior tends to underestimate posterior variance in latent variable models - leading to simulated cells which are over-concentrated around cluster means. [1] This is an inherent limitation of VI-based simulation (especially from mean-field approximations) but is not a fundamental constraint of the generative model itself used to perform the clustering and dimensionality reduction. However, we see overestimation in our resulting mean-variance plot. Using the inverse dispersions to initialise the model encourages the model to capture gene variability through the latent mean structure, enabling more accurate cell-type identification, but is likely to inflate variance in simulated data. A more faithful replication of the mean-variance relationship may require a richer posterior approximation or a dedicated simulation framework.

UMAP visualisations reveal that the simulated dataset maintains distinct, well-separated populations analogous to those in the real data. To validate the ground truth of our generative process, we applied scFLAME to re-cluster the newly simulated data. The model successfully recovered the latent cluster assignments it originally generated, achieving ARIs of [0.999, 0.995, 0.998] across 3 different simulated datasets.

Figures S2 and S3 showcase full-genome simulations of the Baron dataset alongside the less sparse, Smart-Seq2-derived Segerstolpe pancreas dataset. While primarily designed for clustering and dimensionality reduction, scFLAME features a generative architecture allowing users to produce synthetic data that mimics the complexity and structural heterogeneity of real single-cell tissues. Many popular single-cell simulation methods such as Splat do not have the capability to both estimate and simulate from multiple groups, but scFLAME naturally characterises individual clusters. The model also offers users control over simulation parameters, including cell counts, the Dirichlet($\alpha$) cluster proportion prior, and gene configurations (via subsetting or repetition). However, the mean-variance relationship is less faithfully reproduced compared to dedicated simulation frameworks such as Splat, and we recommend users treat scFLAME's generative capabilities as a useful auxiliary feature rather than a replacement for dedicated simulation tools.

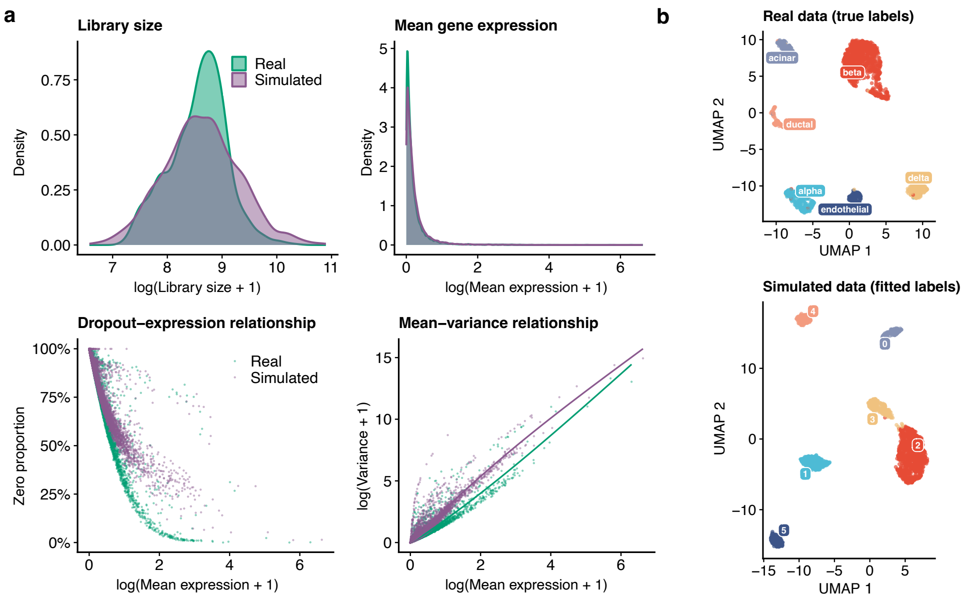


Figure S2: Evaluation of one example of scFLAME simulated data of the full genome against the full genome of the Baron pancreas dataset. **a,** Comparison of common quality control metrics between real (green) and simulated (purple) datasets. We see that simulating the whole genome shows similar results to filtering to the top 5000 HVGs. The average proportion of zeroes in 3 generated datasets was 0.855, versus 0.853 in the true data. **b,** UMAP embeddings of the real Baron pancreas data annotated by true biological cell-types, and the simulated dataset annotated by predicted labels by running scFLAME on the simulated data. The ARI of simulated to predicted labels was 0.993 in this simulation.

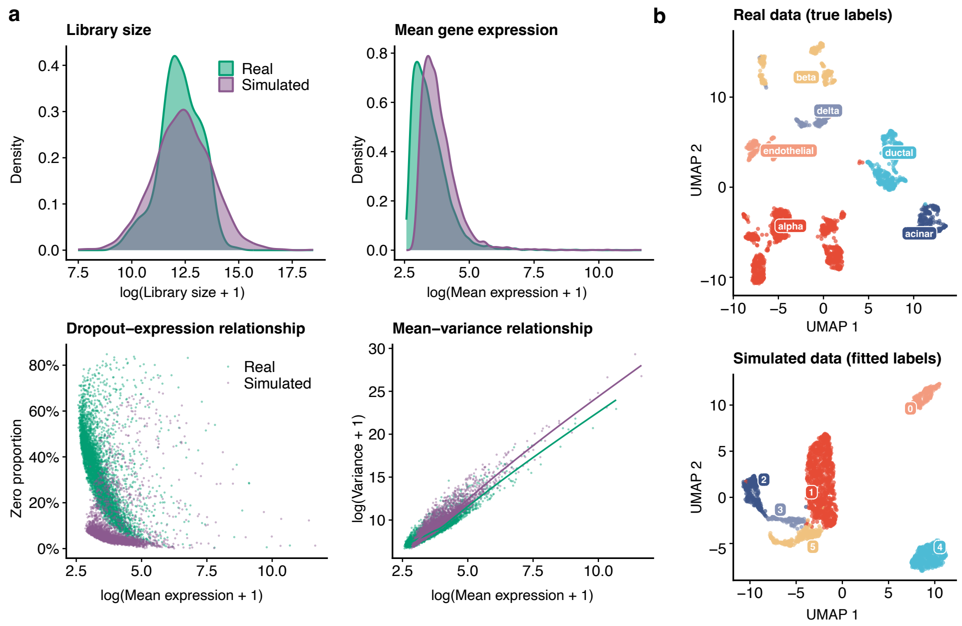


Figure S3: Evaluation of one example of scFLAME simulated data against Segerstolpe pancreas dataset (using Smart-seq2 technologies). **a,** Comparison of common quality control metrics between real (green) and simulated (purple) datasets. We see that in this less sparse data, the dropout rate is lower in the simulated data, although the mean-variance relationship is still captured. The average proportion of zeroes in 3 generated datasets was 0.077, versus 0.330 in the true data. **b,** UMAP embeddings of the real Segerstolpe pancreas data annotated by true biological cell-types, and the simulated dataset annotated by predicted labels by running scFLAME on the simulated data. The ARI of simulated to predicted labels was 0.997 in this simulation.

### 4 Supplementary Results

#### 4.1 scFLAME recovers latent structure in simulated data

Extended results for NMI and clustering accuracy, and additional latent-space visualisations, are shown in Supplementary Figures S4, S5 and S6.

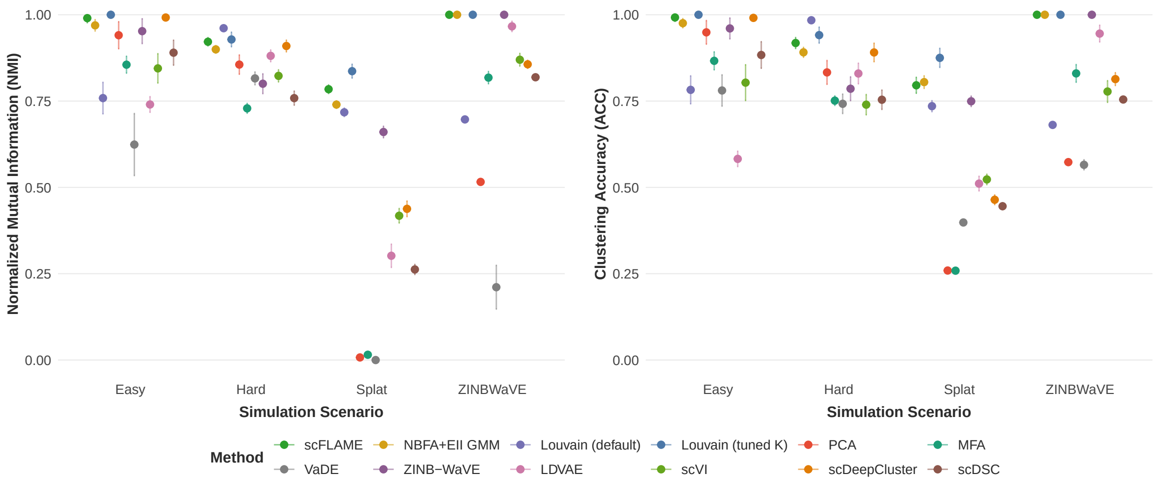


Figure S4: Comparative performance of scFLAME against competitor methods across simulated scenarios, comparing Normalised Mutual Information (NMI) and Clustering Accuracy (ACC) across four simulation scenarios: Easy, Hard, Splat, and ZINB-WaVE. Dots represent the mean NMI/ACC across 3 independent runs over 20 simulated datasets, with error bars indicating ± s.e. (standard error).

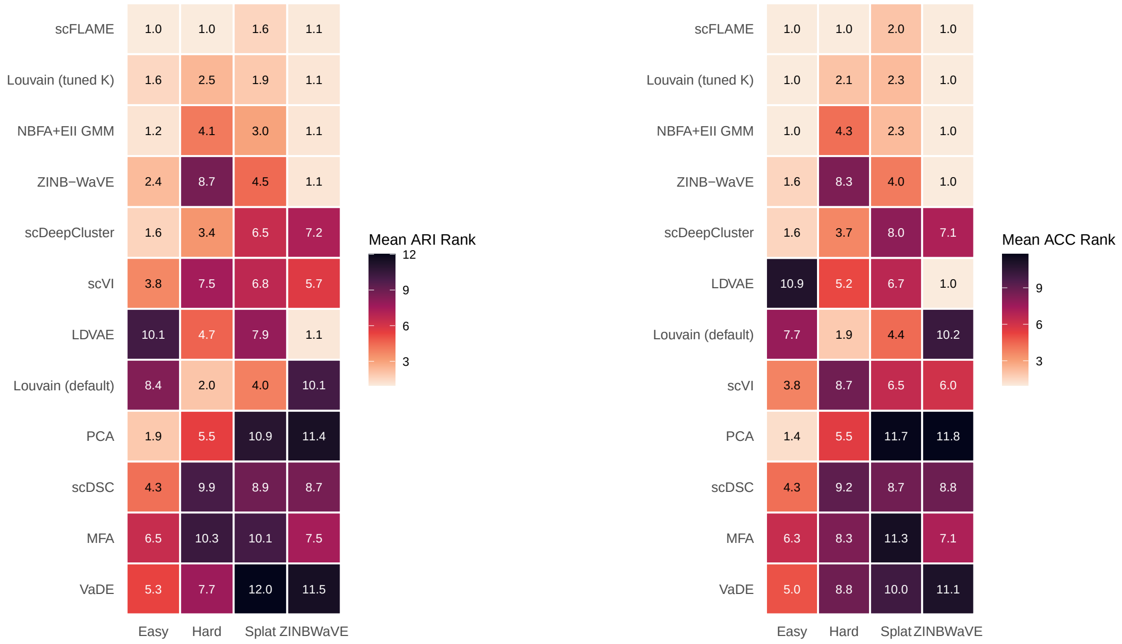


(a) (b)

Figure S5: Relative performance ranking of methods across simulation scenarios from the main text. Heatmaps display the mean rank (where 1 is the top-performing method) for the best result of 3 runs as measured by NMI and ACC. Cell values and colours represent the mean rank across replicates; lighter colours indicate superior performance (lower rank).

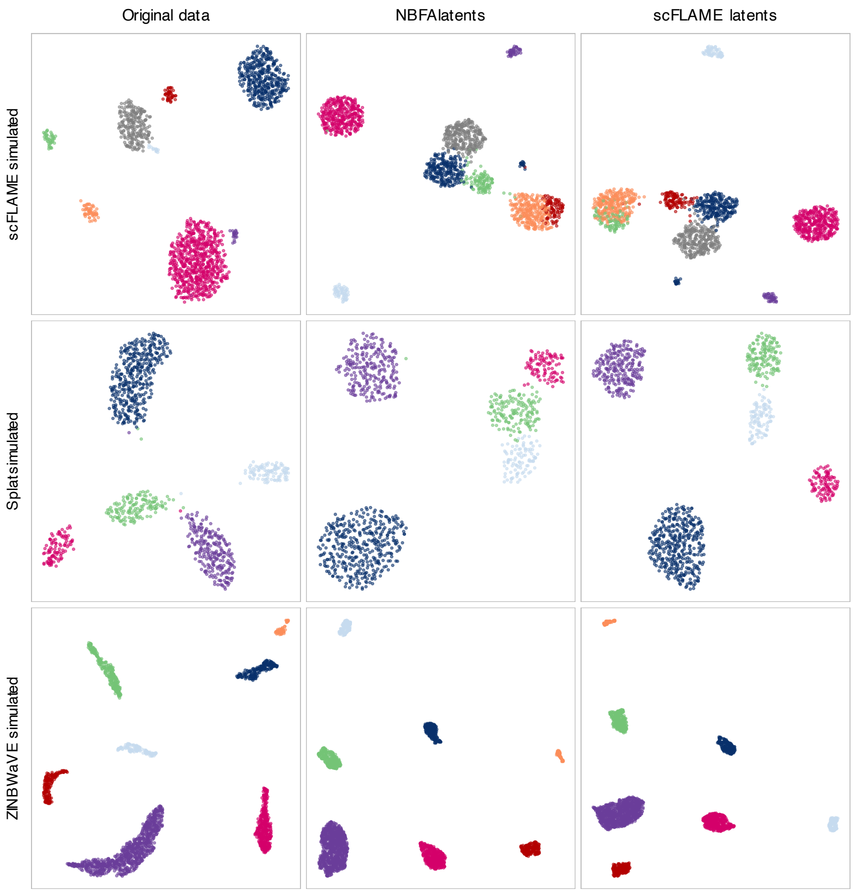


Figure S6: Comparative latent space visualisations across simulation methods. Panels show UMAP projections of original count data and latent representations. Column 1: Original data coloured by true cluster labels. Column 2: NBFA latent space with K-means clustering. Column 3: scFLAME latent space with derived cluster assignments. Rows represent different simulation methods: scFLAME, Splat, and ZINBWaVE. All visualisations use shared UMAP parameters (n_neighbors=15, min_dist=0.5) for comparability.

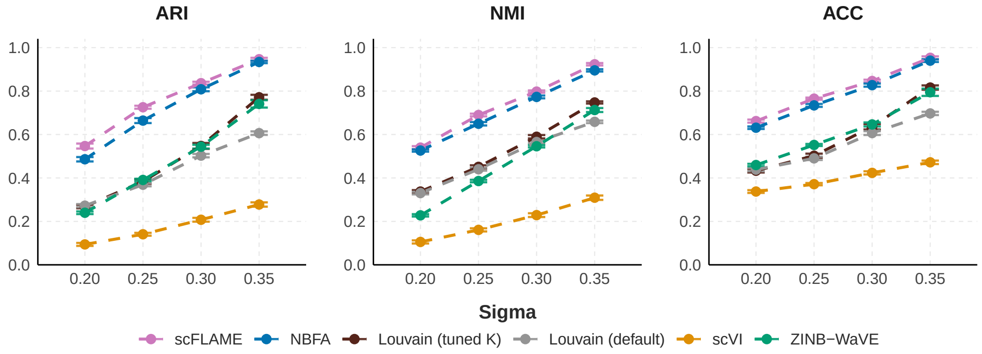


Figure S7: Performance comparison of scFLAME against five baseline methods across various clustering signal strengths, controlled via the 'de.facScale' parameter in Splatter - this is referred to as 'Sigma'. Higher Sigma indicates a stronger clustering signal. Clustering performance is evaluated using three complementary metrics: ARI, NMI, ACC. Points represent the mean performance across $n = 20$ independent simulations with 3 repeats per method per dataset, with dashed lines indicating performance trends. Error bars indicate $\pm$ s.e.m. (standard error of the mean)

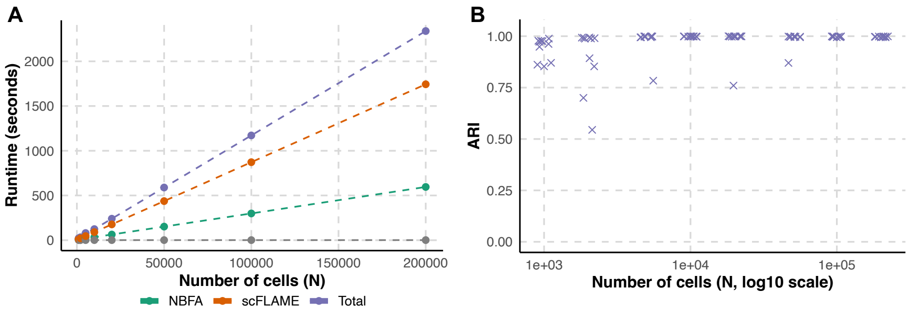


Figure S8: Wall-clock runtime and clustering performance of scFLAME as a function of dataset size. **a,** Wall-clock runtime for simulated datasets with 4,000 genes and increasing numbers of cells ($N$), using a fixed mini-batch size of 8,192 cells once $N$ exceeded the batch size. The NBFA warm-up and scFLAME clustering stages were run for 500 and 300 epochs, respectively. Lines show mean runtime across 10 independent simulated repeats for the NBFA initialisation stage, the scFLAME clustering stage, and the total runtime. **b,** Clustering accuracy measured by adjusted Rand index (ARI) across the same simulations.

##### 4.1.1 Additional simulation scenarios

Extra simulation scenarios considered here are Splat and ZINB-WaVE simulated data using parameters from the Segerstolpe pancreas data [2] with the same cluster parameters as used in the main text ('Methods'). We further examined manually simulating 50% extra dropout using the ZINB-WaVE model with the Segerstolpe parameters, and setting 'dropout.mid' = 2 in the Splat model with the Baron fitted parameters, which produces a very sparse dataset due to the high sparsity of the original data. We see that in these scenarios, scFLAME continues to be the strongest performer, and performs substantially better than other methods from the literature for the very sparse Baron data with extra simulated dropout.

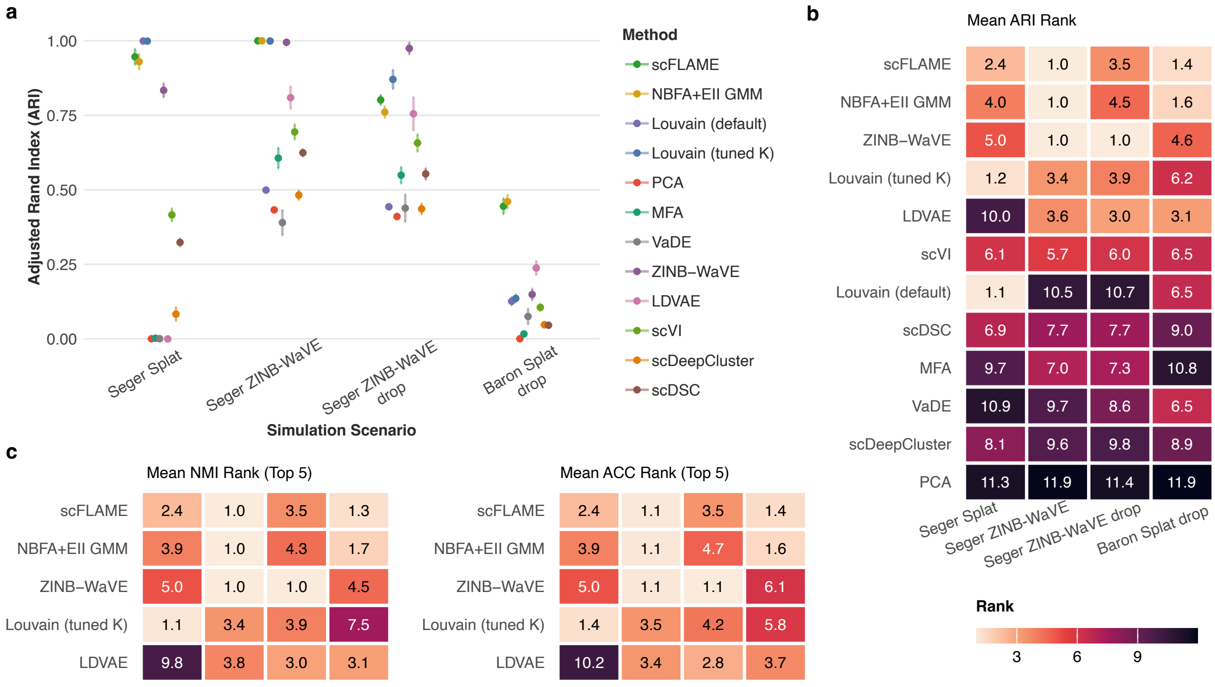


Figure S9: Comparative performance of scFLAME against competitor methods across simulated scenarios. **a,** Clustering accuracy measured by Adjusted Rand Index (ARI) across four simulation scenarios. Dots represent the mean ARI across 3 independent runs over 20 simulated datasets, with error bars indicating ± s.e. (standard error). **b,** Relative performance ranking of methods across simulation scenarios. The heatmap displays the mean rank (where 1 is the top-performing method) for the best performing replicate across 3 repeats for each method and each dataset as measured by ARI. Lighter cell colours indicate superior performance (lower rank). **c,** Relative performance ranking of methods across simulation scenarios, displaying the mean rank for the best perfoming replicate across 3 repeats for each method and each dataset as measured by NMI and ACC. Only the top 5 methods are shown

#### 4.2 scFLAME identifies known cell types in benchmark single-cell datasets

Table S3: Median wall-clock run-times in seconds for all methods on the PBMC 4K dataset across 5 independent runs, reported as median (min, max). Pre-train/warm-up time and main training time breakdowns are shown where applicable. Clustering with k-means for methods where required took negligible time. ZINB-WaVE was run on 4 CPU cores, PCA/Louvain was run on 2 CPU cores and other methods were run on 1 GPU. We note that scDeepCluster is usually run with Tensorflow, but the package version was incompatible with our computing cluster. We filtered to 1000 highly variable genes (HVGs) for ZINB-WaVE as the authors note the high computational complexity of running with more genes; all others were run with 5000 HVGs to match scFLAME. A subset of runs on all methods were subject to occasional GPU-scheduling lag.

| Method | Pre-train/warm-up (s) | Train (s) | Total (s) |
|---|---|---|---|
| scFLAME | 41.8 (41.8, 112.2) | 145.9 (145.8, 149.3) | 187.7 (187.7, 261.5) |
| NBFA+EII GMM | | | 42.9 (42.8, 48.0) |
| Louvain (tuned K) | | | 230.1 (229.7, 250.1) |
| PCA + k-means | | | 216.3 (215.9, 235.9) |
| MFA | | | 96.8 (96.8, 105.1) |
| VaDE | 3.3 (2.9, 35.7) | 54.1 (48.6, 56.6) | 57.6 (51.5, 92.3) |
| ZINB-WaVE + k-means | | | 1735.5 (1589.5, 1798.6) |
| LDVAE + k-means | | | 93.8 (93.3, 103.7) |
| scVI + k-means | | | 90.4 (83.6, 97.9) |
| scDeepCluster | 1143.4 (1142.1, 1152.8) | 94.6 (84.8, 130.9) | 1246.6 (1126.9, 1274.8) |
| scDSC | 11.7 (11.7, 41.8) | 14.0 (14.0, 27.9) | 25.7 (25.7, 69.7) |

Table S4: Average per-cell-type silhouette width in the Pancreas dataset latent/embedding space, comparing no batch correction, scFLAME with the batch-correction term, and Harmony correction. Ground-truth cell-type labels were used as cluster assignments. scFLAME latents are 15-dimensional; Harmony embeddings are 30-dimensional. Silhouette width was significantly higher for scFLAME with batch correction than without (paired $t$-test, $P = 0.0025$; Wilcoxon signed-rank, $P = 0.016$; $n = 7$ cell types).

| **Cell type** | **scFLAME (no batch correction)** | **scFLAME (batch-corrected)** | **Harmony** |
|---|---|---|---|
| Acinar | 0.354 | 0.424 | 0.346 |
| Alpha | 0.175 | 0.287 | 0.193 |
| Beta | 0.140 | 0.228 | 0.156 |
| Delta | 0.242 | 0.290 | 0.182 |
| Ductal | 0.236 | 0.274 | 0.166 |
| Gamma | 0.217 | 0.306 | 0.208 |
| Other | 0.115 | 0.127 | 0.096 |

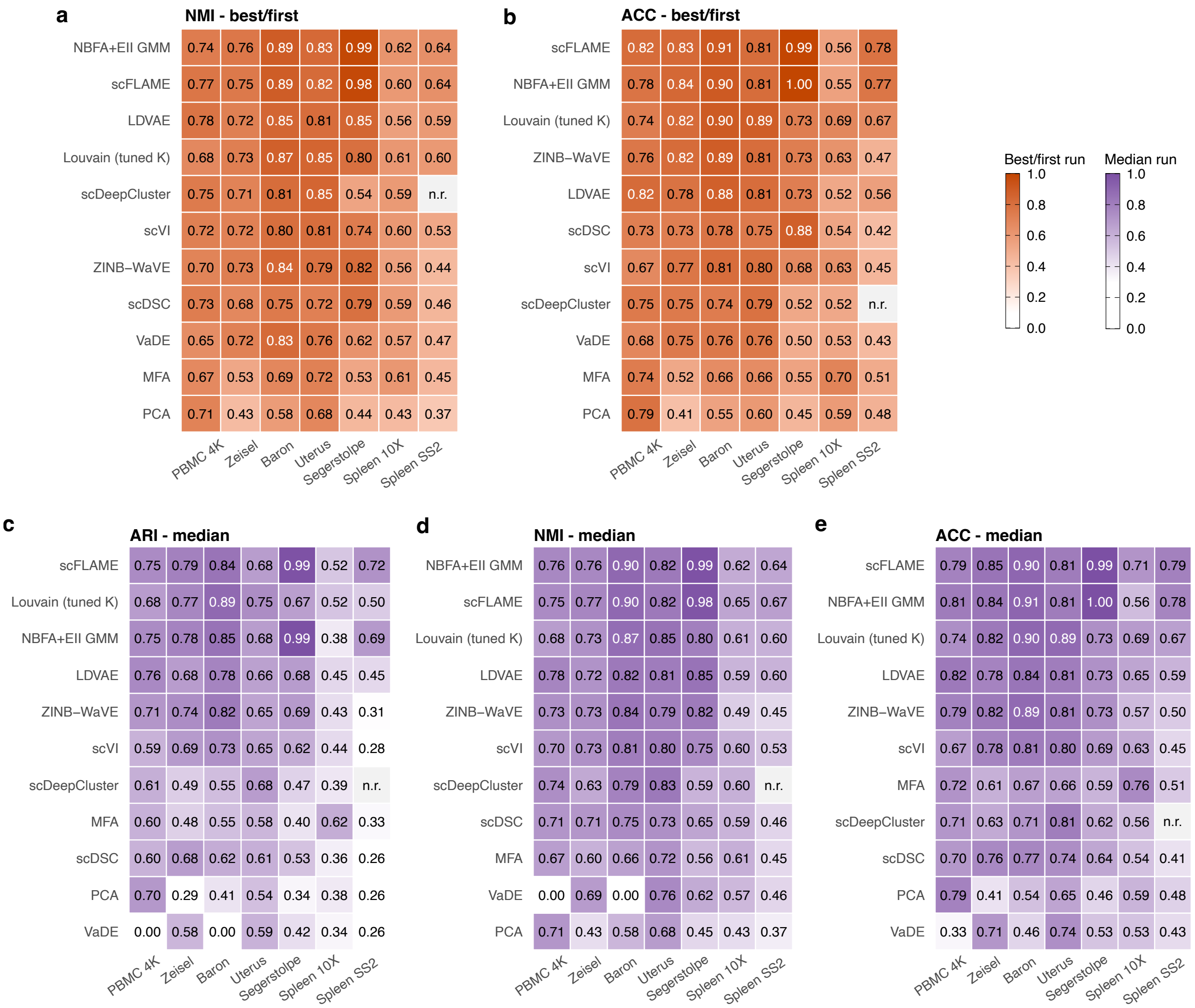


Figure S10: Summary of clustering metrics (ARI, NMI, ACC) between inferred and true cell-type labels for scFLAME and comparator methods across seven real scRNA-seq datasets (PBMC 4K, Zeisel, Baron, Uterus, Segerstolpe, Spleen 10x, Spleen SS2), looking at the 'best' of 5 or the first run in the top panels, and the median of 5 runs in the bottom panels. Missing entries indicate the method ran into numerical errors. Rows are ordered by the median rank of the method across all datasets. **a,** NMI between inferred and true cell-type labels where the 'best' of 5 or the first run is taken. **b,** ACC between inferred and true cell-type labels where the 'best' of 5 or the first run is taken. **c,** ARI between inferred and true cell-type labels where the median of 5 runs is taken. **d,** NMI between inferred and true cell-type labels where the median of 5 runs is taken. **e,** ACC between inferred and true cell-type labels where the median of 5 runs is taken.

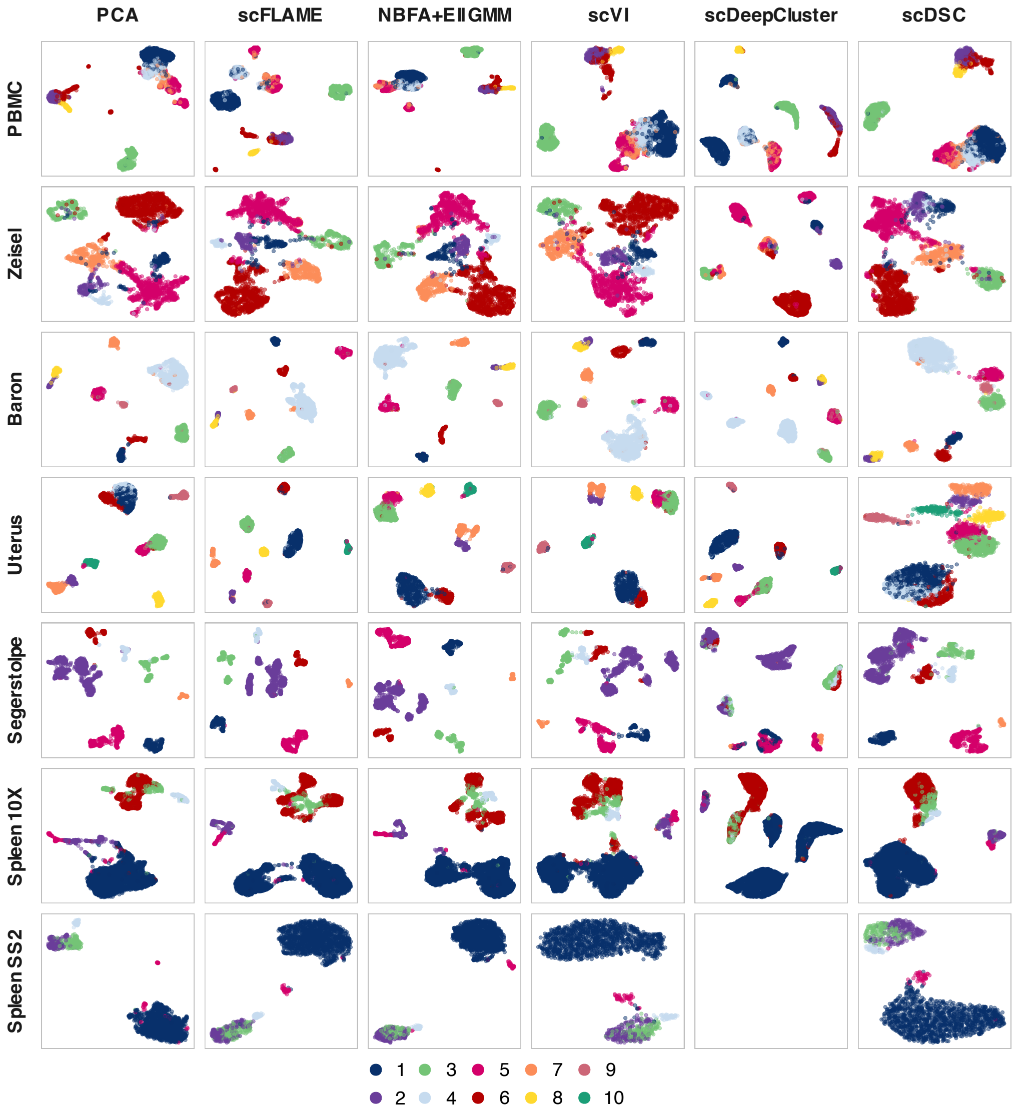


Figure S11: Two-dimensional UMAP projections of single-cell transcriptomes across seven moderately-sized datasets embedded into a shared latent space. Cells are colour-coded according to reference author-annotated labels, with cluster number to cell type mapping in Supplementary Table S2. Representative methods were selected to include the PCA baseline, proposed approaches, and established deep-learning-based methods, where scDSC and scDeepCluster both also perform joint dimensionality reduction and clustering, and scVI is a similar count-based variational dimensionality reduction approach. Projections show that scFLAME successfully finds a latent space where cell types are clearly separated, and latent embeddings separate clusters more than the NBFA baseline.

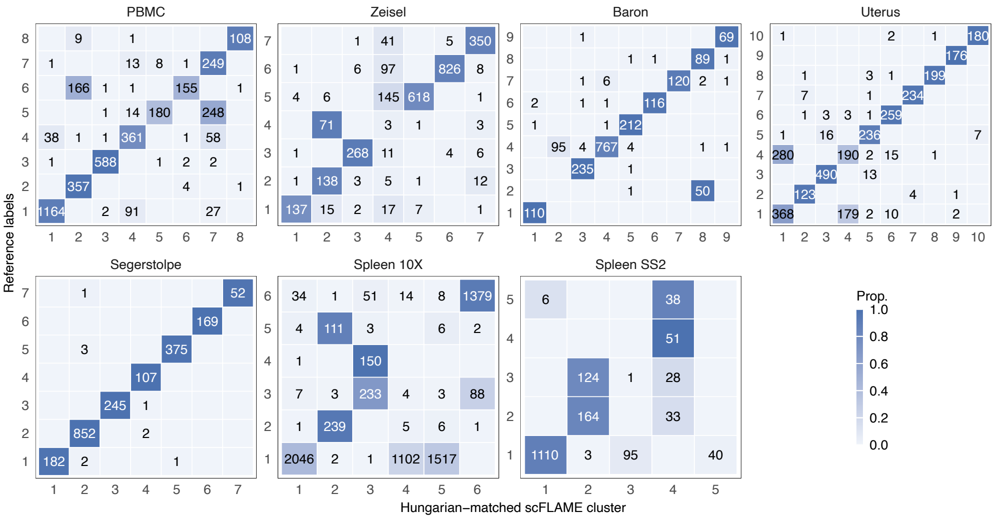


Figure S12: Confusion matrices showing Hungarian-algorithm-matched correspondence between scFLAME cluster assignments and author-provided reference cell type labels, for the PBMC 4K, Zeisel, Baron, Uterus, Segerstolpe, Spleen 10X, and Spleen SS2 datasets. The y-axis shows reference cluster labels (numbered per dataset; see Table/Section 2.2 for correspondence to named cell types); the x-axis shows the scFLAME cluster optimally matched to each reference label via the Hungarian algorithm. Tile colour indicates the proportion of cells from each reference cluster assigned to the matched scFLAME cluster (row-normalised), with raw cell counts annotated in each tile.

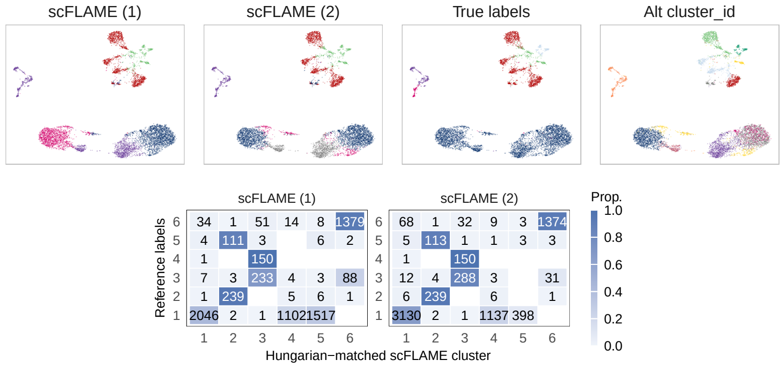


Figure S13: Top: Two-dimensional UMAP projections of scFLAME latent embeddings of Spleen 10X single-cell transcriptomes. Cells are colour-coded according to (1) scFLAME clusters, for the run with the best ELBO (ARI = 0.377) (2) scFLAME clusters, for the run with the second-best ELBO (ARI = 0.518) (3) Author-annotated cell types we benchmarked against (4) Alternative cluster IDs provided by the authors. Bottom: Hungarian-matched confusion matrices validating cross-dataset label correspondence for the two scFLAME runs. We see that scFLAME splits up a large cell type (B-cells) into sub-clusters, but clusters are in line with finer clusters provided by the authors.

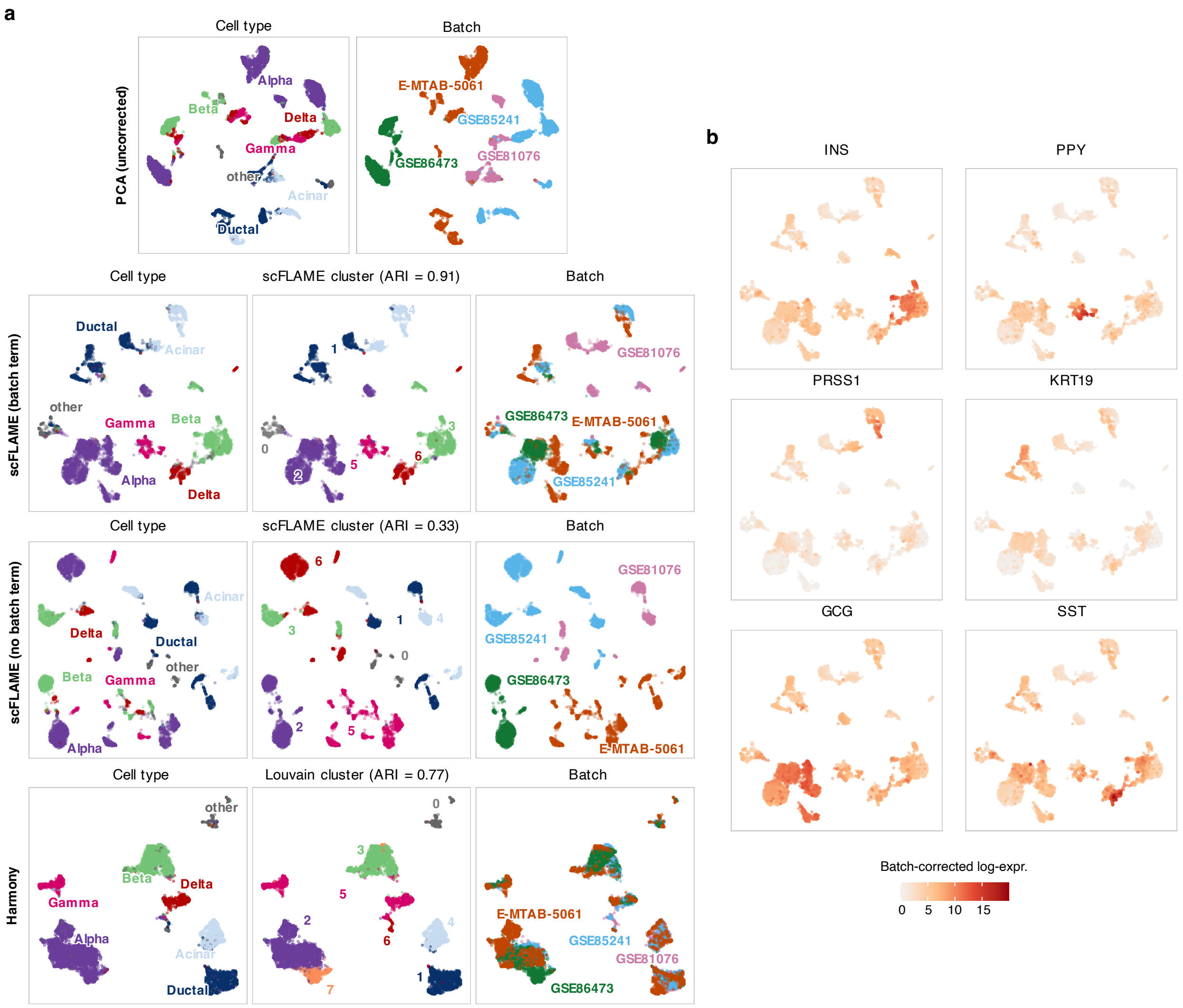


Figure S14: **a,** UMAP projections of the Pancreas dataset comparing reference cell type annotations, cluster assignments, and batch structure across four representations: PCA (uncorrected), scFLAME with a batch term, scFLAME without a batch term, and Harmony. Clustering agreement with reference cell types is reported as the Adjusted Rand Index (ARI) for scFLAME (batch term, ARI=0.91), scFLAME (no batch term, ARI=0.33), and Harmony (Louvain, ARI=0.77); UMAP of the PCA reduction of the original data is shown uncorrected for reference and has no associated clustering. As with other datasets, we performed a resolution sweep in steps of 0.05 to find a solution with as close to the reference number of clusters as possible; Louvain was ran with 'resolution = 0.2', resulting in 8 clusters. 6 clusters were obtained with 'resolution = 0.15'. Points in the batch panels are coloured by sequencing batch (E-MTAB-5061, GSE85241, GSE86473, GSE81076). **b,** Batch-corrected log-expression levels for marker genes *INS*, *PPY*, *PRSS1*, *KRT19*, *GCG*, and *SST*, defined as the posterior mean of $\mu_{ij}$, evaluated at the reference batch level ($\delta_{bj} = 0$), shown on the scFLAME (batch term) embedding.

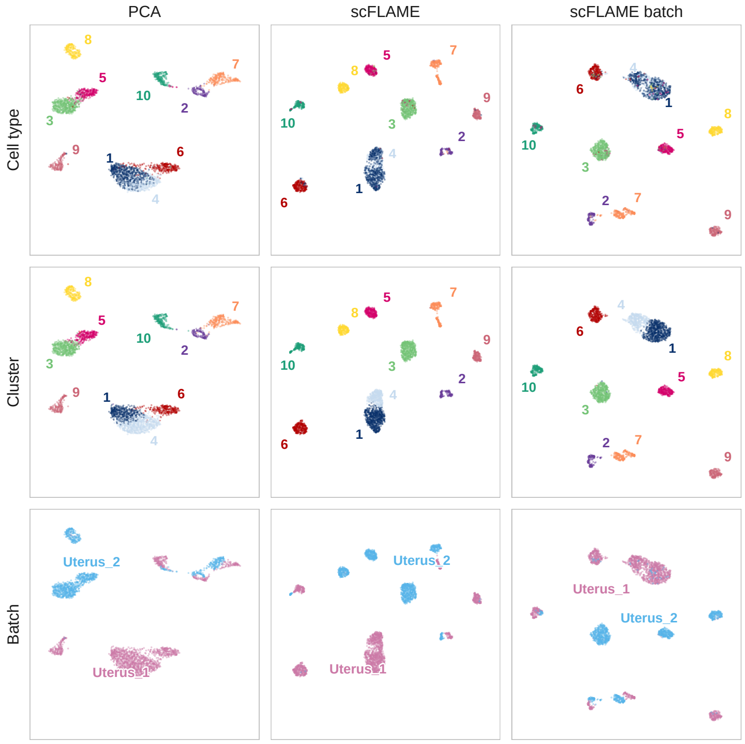


Figure S15: UMAP projections of the Uterus dataset comparing reference cell types, cluster assignments, and batch structure across three representations: PCA, scFLAME (no batch term), and scFLAME (batch term). Cluster assignments (middle row) are shown for Louvain clustering on PCA latents (ARI=0.75), scFLAME (no batch term, ARI=0.68), and scFLAME (batch term, ARI=0.64). Cluster labels are Hungarian-matched to reference cell types, with cluster number corresponding to reference label as given in Table 2.2. Points in the bottom row are coloured by sequencing batch (Uterus_1, Uterus_2). Despite not improving clustering performance against reference annotations, adding the batch term visually improves some mixing of batches e.g. in Cluster 10.

##### 4.2.1 Ablations

**Clustering initialisation** To assess sensitivity to initialisation, we compared the default EII GMM initialisation with two alternative initialisations of the mixture prior's cluster means ($\nu_k$) and variance ($\Omega$):

- **Random:** $\nu_k$ initialised by sampling $K$ unique observations without replacement from $m$, the latent mean vector. $\Omega$ is set to the global variance of $m$. $\pi_k$ is uniform $(1/K)$.
- **Random far:** $\nu_k$ drawn independently of the data from a broad isotropic Gaussian distribution $\mathcal{N}(0, 3^2 \cdot I)$. $\Omega$ is set to 1 globally. $\pi_k$ is uniform $(1/K)$.

Overall, EII GMM provides the most stable behaviour, generally yielding strong and consistent performance across runs in terms of ARI, NMI and ACC (Supplementary Figure **??**). We see less consistent behaviour in the TM Spleen 10X dataset due to the imbalance in cluster sizes mentioned previously. Standard data-informed random initialisations, which select starting component means directly from latent means $m_i$, produce comparable results to the EII GMM initialisation in many cases. This indicates that the scFLAME optimisation landscape is generally robust to local perturbations within the NBFA latent space. Occasional drops in performance with this initialisation likely stem from stochastic edge cases.

'Random_far' initialisations exhibit substantially higher variance. While in most runs they lead to lower or comparable performance due to initialisation in very sparse regions of the latent space, in some cases they converge to alternative local optima that substantially improve clustering quality with respect to authors' labels (e.g. ARI = 0.857 on Zeisel and ARI = 0.904 on TM spleen 10X) (Supplementary Figure S16). These cases correspond to different partitions of the latent space, suggesting that in noisier datasets the model may admit multiple plausible clusterings. In general, the random initialisations performed better for the TM Spleen 10X dataset, highlighting their utility in breaking free from symmetric structural constraints.

We recommend EII GMM as the default initialisation to capture the most stable and prominent structure in the data. ELBO-based model selection remains most reliable when restricted to stable, structurally congruent EII GMM initialisations, where this geometry matches the internal assumptions of the joint model prior. Other initialisations can be used as auxiliary exploratory tools. Another method of identifying subclusters with weaker signal is by looking at increasing the number of initialised clusters to identify rarer sub-populations and using our hierarchical approach.

**Pre-estimating dispersions and library sizes** We investigate the effects of pre-estimating the dispersions with `edgeR` and pre-computing library sizes, compared to having the model infer dispersions initialised from $\phi_j = 5$ for all dispersions, and the library sizes fixed at the same value for all cells. Figure S17 shows the best run of 3 for varying these settings on the Baron data. With dispersion pre-estimated for initialisation, scFLAME was able to clearly separate gamma and delta pancreatic cells - two rare but distinct endocrine pancreatic populations - whereas removing this pre-estimation caused the two populations to be blended into a single cluster in all 6 runs without `edgeR` dispersions. Excluding pre-estimated library size further degraded cluster resolution, muddling boundaries between additional cell types and lowering ARI to 0.77 in the best run as chosen by ELBO; omitting both components together produced the greatest loss of structure (ARI = 0.74). Both pre-estimation steps contribute complementary information toward resolving distinct cell populations. TMM-based library size estimates are widely used and have been shown to improve accuracy in various downstream differential expression analysis tasks [3, 4]; however, our results for the 68K dataset suggest that for large datasets where computing such normalisation factors may be computationally prohibitive, a simpler estimate of library size - for example, the total count sum per cell - may be sufficient.

**Number of genes** To examine how scFLAME's runtime and accuracy changed with the number of input HVGs, we ran the model on the PBMC dataset with HVGs ranging from 2000-16656 (the full gene set after filtering out e.g. genes with close to 0 counts across all cells), with three repeats per setting. Total runtime scaled near-linearly with gene count (linear fit $R^2 = 0.997$). Clustering accuracy (ARI) was generally stable across gene counts, but the mean accuracy across runs was lower when restricted to 2,000 genes, indicating a moderate gene set is required to capture sufficient biological signal for reliable clustering. Further increasing the gene count beyond around 5,000-8,000 genes yields no significant accuracy gain despite the added computational cost, as it is likely most of these genes have little signal. However, accuracy is not depleted, showing that our dimensionality reduction is still able to detect biological signal even with additional noisy genes.

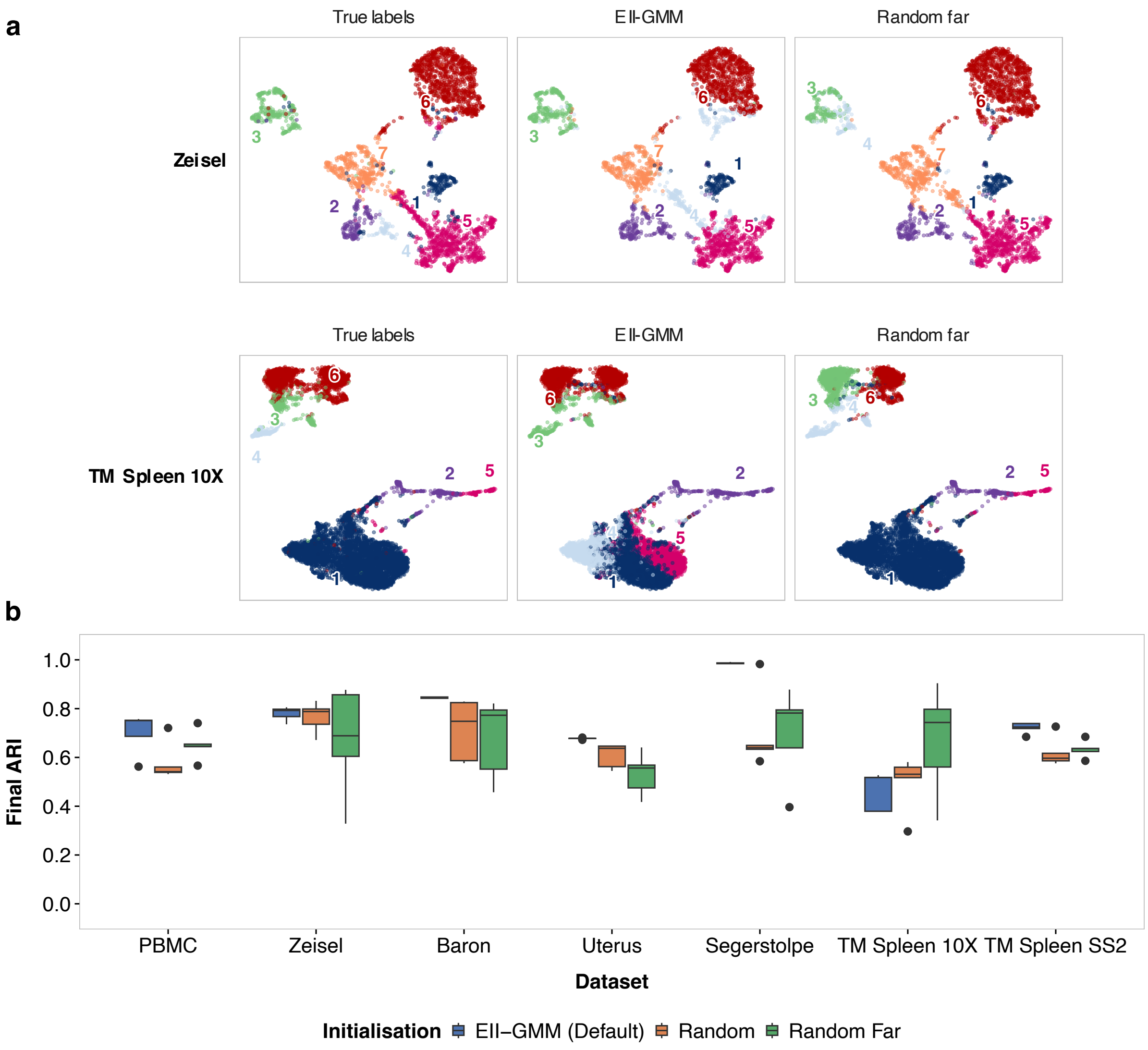


Figure S16: Evaluation of scFLAME clustering across different clustering initialisation strategies. **a,**UMAP visualisations contrasting EII GMM (middle) and 'random_far' (right) clustering initialisations against ground-truth cell annotations (left) for the Zeisel and TM spleen 10X datasets. We visualise the PCA-reduction of the original data in both cases. For both datasets, the underlying model corresponds to the specific seed replication that yielded the highest Evidence Lower Bound (ELBO) during NBFA + EII GMM warm-up, and this same latent space initialisation was used for the two different clustering initialisations. For Zeisel, the clustering gave an ARI of 0.768 and NMI of 0.755 under EII GMM initialisation, compared to ARI = 0.857 and NMI = 0.797 under 'random_far' initialisation. For TM spleen 10X, the clustering gave an ARI of 0.377 and NMI of 0.596 for EII GMM, and ARI = 0.904 and NMI = 0.790 for 'random_far'. **b,** Box plots illustrating the final ARI achieved by scFLAME across seven scRNA-seq datasets over five independent runs per strategy. Optimisation runs were initiated using three distinct regimes: the structural baseline matching the joint model prior (EII GMM, blue), data-informed random sampling from the latent coordinates (Random, orange), and out-of-distribution random generation (Random Far, green). Centre lines represent the median; box limits indicate the upper and lower quartiles; whiskers extend to 1.5× the interquartile range; and individual points represent statistical outliers.

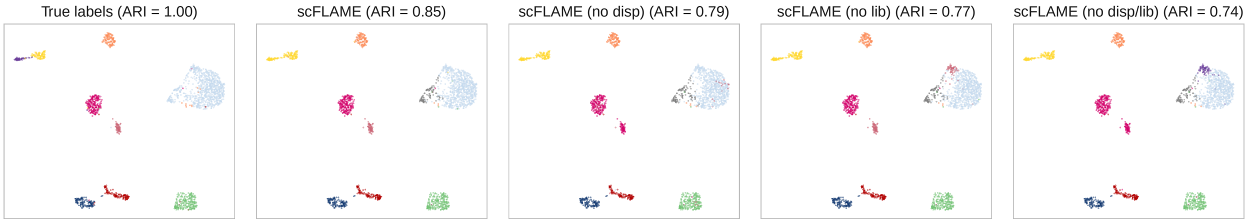


Figure S17: UMAPs of the Baron dataset coloured by Hungarian-matched clusters for true labels and scFLAME ran (i) with pre-estimated library sizes and dispersions (ii) without pre-estimated dispersions (iii) without pre-estimated library sizes (iv) without both, where the best run of 3 by ELBO is shown. UMAP projections are computed from a shared PCA embedding of the raw data. Removing dispersion and library-size components progressively degrades clustering accuracy.

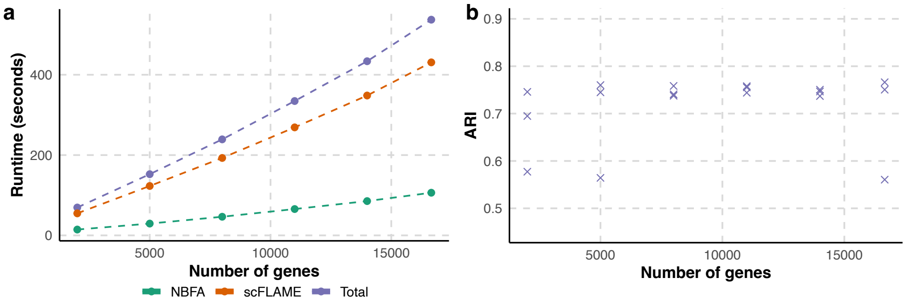


Figure S18: Wall-clock runtime and clustering performance of scFLAME applied to the PBMC 4K dataset as a function of the number of highly variable genes (HVGs) included. **a,** Wall-clock runtime for the PBMC 4K dataset with increasing numbers of genes ($D$), starting from 2000. The NBFA warm-up and scFLAME clustering stages were run for 500 and 300 epochs, respectively. Lines show mean runtime across 3 independent simulated repeats for the NBFA initialisation stage, the scFLAME clustering stage, and the total runtime. **b,** Clustering accuracy measured by adjusted Rand index (ARI) across the same runs.

#### 4.3 Integrated marker gene identification via shared loading matrix

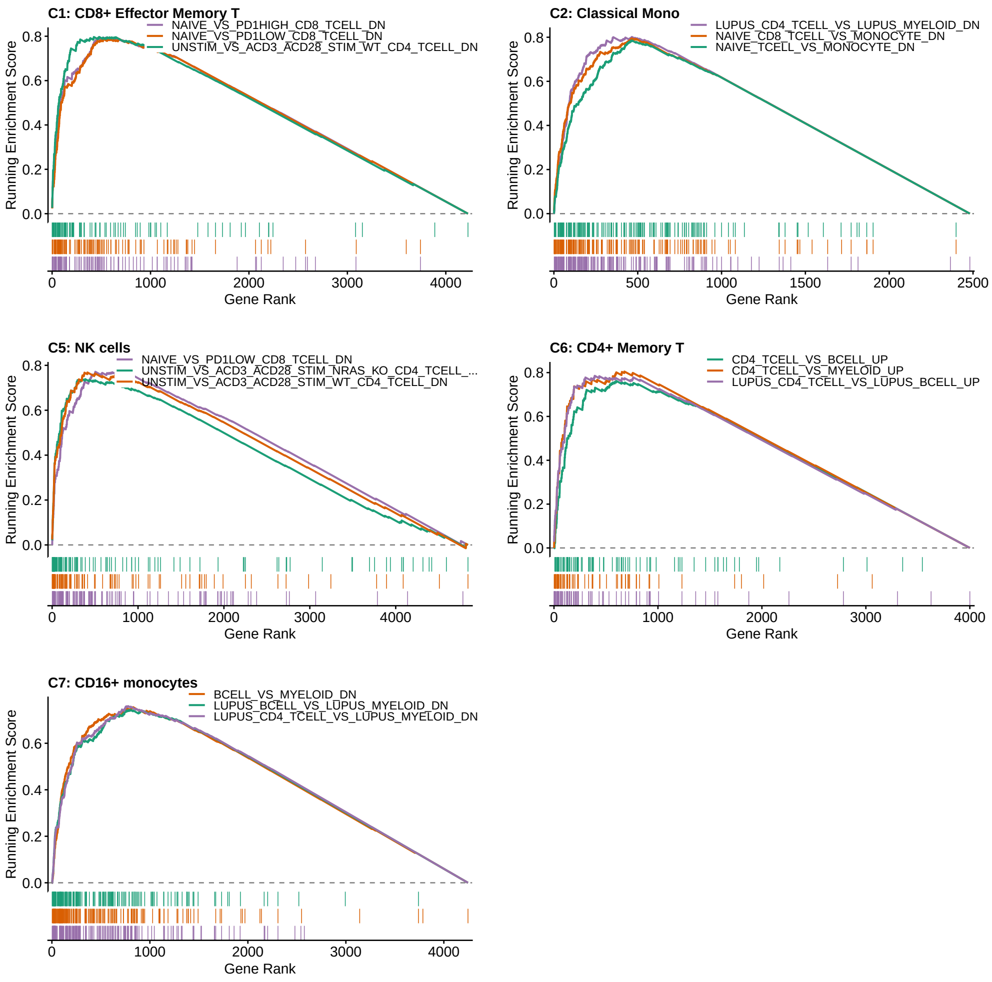


Figure S19: Gene Set Enrichment Analysis (GSEA) enrichment plots for clusters 1, 2, 5, 6, 7 for the PBMC data showing the top three enriched immunological gene sets (MSigDB C7), restricted to GEO-derived signatures (GSE). Running enrichment scores are shown as a function of gene rank.

Table S5: Top 10 enriched MSigDB C7 pathways by cluster in the PBMC data. Only pathways with NES > 0 are shown, ranked by $p_{\text{adj}}$.

| Cluster | Pathway | NES | $p_{\text{val}}$ | $p_{\text{adj}}$ |
|---|---|---|---|---|
| 0 | GSE29618_BCELL_VS_MDC_DAY7_FLU_VACCINE_DN | 3.42 | 1.65e-54 | 8.33e-51 |
| 0 | GSE29618_BCELL_VS_MDC_DN | 3.19 | 1.73e-40 | 4.37e-37 |
| 0 | HOEK_NK_CELL_2011_2012_TIV_3D_VS_0DY_ADULT_3D_DN | 3.02 | 2.51e-36 | 4.24e-33 |
| 0 | GSE10325_LUPUS_CD4_TCELL_VS_LUPUS_MYELOID_DN | 3.01 | 1.51e-34 | 1.91e-31 |
| 0 | GSE10325_CD4_TCELL_VS_MYELOID_DN | 2.95 | 3.79e-31 | 3.84e-28 |
| 0 | GSE22886_NAIVE_CD8_TCELL_VS_MONOCYTE_DN | 2.92 | 1.33e-30 | 1.12e-27 |
| 0 | FLETCHER_PBMC_BCG_10W_INFANT_BCG_STIMULATED_VS_UNSTIMULATED_10W_DN | 2.84 | 6.74e-29 | 4.87e-26 |
| 0 | HARALAMBIEVA_PBMC_M_M_R_II_AGE_11_22YO_VACCINATED_VS_UNVACCINATED_7YR_DN | 2.47 | 3.17e-28 | 2e-25 |
| 0 | GSE22886_NAIVE_TCELL_VS_MONOCYTE_DN | 2.84 | 2.44e-27 | 1.37e-24 |
| 0 | GSE29618_BCELL_VS_PDC_DAY7_FLU_VACCINE_DN | 2.88 | 3.25e-27 | 1.5e-24 |
| 1 | GSE26495_NAIVE_VS_PD1LOW_CD8_TCELL_DN | 3.33 | 8.99e-38 | 3.82e-34 |
| 1 | GSE26495_NAIVE_VS_PD1HIGH_CD8_TCELL_DN | 3.34 | 2.26e-37 | 3.82e-34 |
| 1 | GSE45739_UNSTIM_VS_ACD3_ACD28_STIM_WT_CD4_TCELL_DN | 3.26 | 6.06e-32 | 7.67e-29 |
| 1 | OSMAN_BLOOD_CHAD63_KH_AGE_18_50YO_HIGH_DOSE_SUBJECTS_24HR_DN | 2.61 | 2.02e-29 | 2.05e-26 |
| 1 | KAZMIN_PBMC_P_FALCIPARUM_RTSS_AS01_AGE_UNKNOWN_CORRELATED_WITH_PROTECTION_56DY_NEGATIVE | 3.17 | 1.77e-28 | 1.49e-25 |
| 1 | ZAK_PBMC_MRKAD5_HIV_1_GAG_POL_NEF_AGE_20_50YO_1DY_DN | 2.69 | 2.14e-24 | 1.36e-21 |
| 1 | GSE45739_UNSTIM_VS_ACD3_ACD28_STIM_NRAS_KO_CD4_TCELL_DN | 2.95 | 3.58e-23 | 2.01e-20 |
| 1 | GSE22886_NAIVE_CD8_TCELL_VS_MONOCYTE_UP | 2.97 | 6.33e-22 | 2.91e-19 |
| 1 | GSE22886_CD8_TCELL_VS_BCELL_NAIVE_UP | 2.93 | 1.22e-20 | 5.13e-18 |
| 1 | GSE11057_NAIVE_VS_EFF_MEMORY_CD4_TCELL_DN | 2.73 | 7.32e-19 | 2.65e-16 |
| 2 | HOWARD_PBMC_INACT_MONOV_INFLUENZA_A_INDONESIA_05_2005_H5N1_AGE_19_39YO_AS03_ADJUVANT_VS_BUFFER_1DY_UP | 3.22 | 2.63e-65 | 1.33e-61 |
| 2 | GSE10325_LUPUS_CD4_TCELL_VS_LUPUS_MYELOID_DN | 3.37 | 1.01e-62 | 2.56e-59 |
| 2 | HOEK_NK_CELL_2011_2012_TIV_3D_VS_0DY_ADULT_3D_DN | 3.32 | 5.79e-59 | 9.77e-56 |
| 2 | GSE22886_NAIVE_CD8_TCELL_VS_MONOCYTE_DN | 3.33 | 1.07e-57 | 1.35e-54 |
| 2 | GSE22886_NAIVE_TCELL_VS_MONOCYTE_DN | 3.29 | 2.83e-54 | 2.87e-51 |
| 2 | GSE22886_NAIVE_CD4_TCELL_VS_MONOCYTE_DN | 3.27 | 3.33e-53 | 2.81e-50 |
| 2 | GSE10325_LUPUS_BCELL_VS_LUPUS_MYELOID_DN | 3.24 | 6.43e-52 | 4.65e-49 |
| 2 | GSE29618_MONOCYTE_VS_PDC_UP | 3.26 | 1.61e-49 | 9.05e-47 |
| 2 | GSE29618_BCELL_VS_MONOCYTE_DAY7_FLU_VACCINE_DN | 3.22 | 5.92e-49 | 3e-46 |
| 2 | GSE10325_BCELL_VS_MYELOID_DN | 3.16 | 6.15e-48 | 2.83e-45 |
| 3 | GSE10325_CD4_TCELL_VS_BCELL_DN | 3.55 | 2.25e-59 | 1.14e-55 |
| 3 | GSE10325_LUPUS_CD4_TCELL_VS_LUPUS_BCELL_DN | 3.51 | 4.28e-55 | 1.08e-51 |
| 3 | GSE3982_MEMORY_CD4_TCELL_VS_BCELL_DN | 3.27 | 6.9e-39 | 1.16e-35 |
| 3 | GSE10325_BCELL_VS_MYELOID_UP | 3.23 | 3.64e-33 | 3.69e-30 |
| 3 | FOURATI_BLOOD_TWINRIX_AGE_25_83YO_RESPONDERS_VS_POOR_RESPONDERS_0DY_UP | 3.04 | 4.04e-31 | 3.41e-28 |
| 3 | GSE22886_TCELL_VS_BCELL_NAIVE_DN | 3.21 | 6.56e-31 | 4.75e-28 |
| 3 | GSE29618_BCELL_VS_MONOCYTE_DAY7_FLU_VACCINE_UP | 3.2 | 1.33e-30 | 7.74e-28 |
| 3 | GSE3982_BCELL_VS_CENT_MEMORY_CD4_TCELL_UP | 3.11 | 1.38e-30 | 7.74e-28 |
| 3 | GSE22886_CD8_TCELL_VS_BCELL_NAIVE_DN | 3.19 | 2.64e-30 | 1.34e-27 |
| 3 | GSE29618_BCELL_VS_MONOCYTE_UP | 3.18 | 4.43e-30 | 2.04e-27 |
| 4 | GSE10325_LUPUS_CD4_TCELL_VS_LUPUS_BCELL_UP | 3.6 | 4.36e-26 | 2.76e-23 |
| 4 | GSE26495_NAIVE_VS_PD1HIGH_CD8_TCELL_UP | 3.56 | 1.33e-24 | 6.74e-22 |
| 4 | GSE10325_CD4_TCELL_VS_MYELOID_UP | 3.47 | 8.82e-24 | 3.43e-21 |
| 4 | ZAK_PBMC_MRKAD5_HIV_1_GAG_POL_NEF_AGE_20_50YO_1DY_DN | 2.67 | 3.3e-23 | 9.84e-21 |
| 4 | GSE22886_NAIVE_CD4_TCELL_VS_MONOCYTE_UP | 3.49 | 4.12e-23 | 1.16e-20 |
| 4 | OSMAN_BLOOD_CHAD63_KH_AGE_18_50YO_HIGH_DOSE_SUBJECTS_24HR_DN | 2.38 | 6.28e-23 | 1.67e-20 |
| 4 | GSE10325_LUPUS_CD4_TCELL_VS_LUPUS_MYELOID_UP | 3.43 | 1.21e-21 | 2.45e-19 |
| 4 | GSE22886_NAIVE_CD8_TCELL_VS_NKCELL_UP | 3.47 | 1.81e-21 | 3.4e-19 |
| 4 | GSE22886_NAIVE_TCELL_VS_NKCELL_UP | 3.41 | 1.49e-20 | 2.36e-18 |
| 4 | GSE22886_NAIVE_TCELL_VS_MONOCYTE_UP | 3.37 | 3.3e-20 | 4.91e-18 |
| 5 | GSE26495_NAIVE_VS_PD1LOW_CD8_TCELL_DN | 3.34 | 3.33e-35 | 1.69e-31 |
| 5 | GSE45739_UNSTIM_VS_ACD3_ACD28_STIM_WT_CD4_TCELL_DN | 3.21 | 1.18e-28 | 2e-25 |
| 5 | GSE45739_UNSTIM_VS_ACD3_ACD28_STIM_NRAS_KO_CD4_TCELL_DN | 3.15 | 2.28e-27 | 2.89e-24 |
| 5 | GSE26495_NAIVE_VS_PD1HIGH_CD8_TCELL_DN | 3.12 | 1.05e-26 | 1.07e-23 |
| 5 | KAZMIN_PBMC_P_FALCIPARUM_RTSS_AS01_AGE_UNKNOWN_CORRELATED_WITH_PROTECTION_56DY_NEGATIVE | 3.18 | 1.33e-26 | 1.12e-23 |
| 5 | HOFT_PBMC_TICE_BCG_RBCG_AG85A_AG85B_AGE_18_40YO_CORRELATED_WITH_WHOLE_BLOOD_BACTERICIDAL_ACTIVITY_NEGATIVE | 2.96 | 4.78e-23 | 3.46e-20 |
| 5 | GSE22886_NAIVE_CD8_TCELL_VS_NKCELL_DN | 2.95 | 7.76e-18 | 4.91e-15 |
| 5 | GSE26495_PD1HIGH_VS_PD1LOW_CD8_TCELL_DN | 2.81 | 2.24e-15 | 1.26e-12 |
| 5 | GSE22886_NAIVE_TCELL_VS_NKCELL_DN | 2.77 | 3.32e-14 | 1.68e-11 |
| 5 | GSE21063_WT_VS_NFATC1_KO_16H_ANTI_IGM_STIM_BCELL_DN | 2.75 | 2.04e-13 | 9.4e-11 |
| 6 | GSE10325_LUPUS_CD4_TCELL_VS_LUPUS_BCELL_UP | 3.17 | 1.53e-25 | 2.36e-22 |
| 6 | GSE10325_CD4_TCELL_VS_BCELL_UP | 3.16 | 3.61e-25 | 3.41e-22 |
| 6 | GSE10325_CD4_TCELL_VS_MYELOID_UP | 3.14 | 1.5e-23 | 1.02e-20 |
| 6 | GSE10325_LUPUS_CD4_TCELL_VS_LUPUS_MYELOID_UP | 3.04 | 6.85e-21 | 3.15e-18 |
| 6 | GSE22886_NAIVE_CD4_TCELL_VS_MONOCYTE_UP | 3.01 | 1.16e-20 | 4.89e-18 |
| 6 | GSE22886_NAIVE_VS_MEMORY_TCELL_DN | 3.01 | 1.43e-20 | 5.56e-18 |
| 6 | GSE11057_PBMC_VS_MEM_CD4_TCELL_DN | 3.06 | 2.22e-20 | 8.03e-18 |
| 6 | GSE22886_NAIVE_CD8_TCELL_VS_MEMORY_TCELL_DN | 2.99 | 2.55e-20 | 8.17e-18 |
| 6 | GSE45739_UNSTIM_VS_ACD3_ACD28_STIM_NRAS_KO_CD4_TCELL_UP | 3 | 2.69e-20 | 8.17e-18 |
| 6 | GSE11057_NAIVE_VS_MEMORY_CD4_TCELL_DN | 2.79 | 4.04e-19 | 1.14e-16 |
| 7 | GSE10325_LUPUS_CD4_TCELL_VS_LUPUS_MYELOID_DN | 3.38 | 4.19e-51 | 2.12e-47 |
| 7 | GSE10325_BCELL_VS_MYELOID_DN | 3.36 | 7.32e-50 | 1.85e-46 |
| 7 | HOEK_NK_CELL_2011_2012_TIV_3D_VS_0DY_ADULT_3D_DN | 3.31 | 1.25e-48 | 2.11e-45 |
| 7 | GSE10325_LUPUS_BCELL_VS_LUPUS_MYELOID_DN | 3.32 | 2.65e-45 | 3.35e-42 |
| 7 | HOWARD_PBMC_INACT_MONOV_INFLUENZA_A_INDONESIA_05_2005_H5N1_AGE_19_39YO_AS03_ADJUVANT_VS_BUFFER_1DY_UP | 3.09 | 4.52e-45 | 4.58e-42 |
| 7 | NAKAYA_PBMC_FLUARIX_FLUVIRIN_AGE_18_50YO_CORRELATED_WITH_HAI_28DY_RESPONSE_AT_3DY_POSITIVE | 3.09 | 2.87e-43 | 2.42e-40 |
| 7 | GSE22886_NAIVE_CD8_TCELL_VS_MONOCYTE_DN | 3.23 | 3.85e-42 | 2.78e-39 |
| 7 | NAKAYA_PBMC_FLUMIST_AGE_18_50YO_3DY_UP | 2.97 | 1.02e-39 | 6.45e-37 |
| 7 | GSE22886_NAIVE_TCELL_VS_MONOCYTE_DN | 3.18 | 5.67e-39 | 2.87e-36 |
| 7 | GSE29618_BCELL_VS_MONOCYTE_DN | 3.17 | 4.72e-38 | 2.17e-35 |

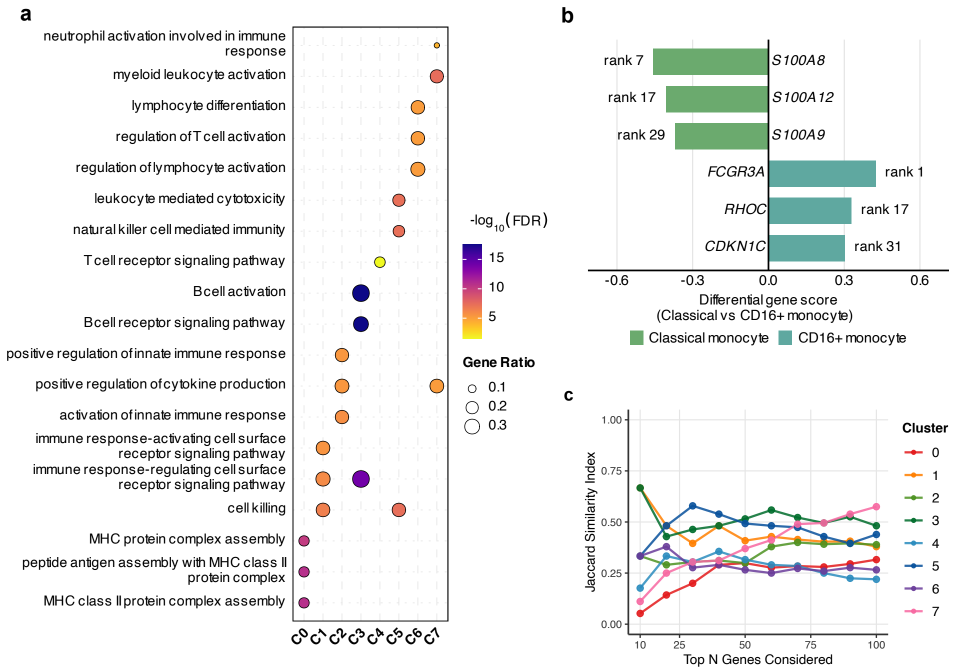


Figure S20: Further characterisation of PBMC clusters identified by scFLAME via gene scores derived from the fitted scFLAME model. **a,** Gene Ontology (GO) Biological Process enrichment of top marker genes across PBMC clusters, showing the top 3 processes per cluster. Dot plot showing over-representation analysis (ORA) of the top 100 positively scored genes per cluster. Each point represents a significantly enriched GO Biological Process term for a given cluster. Dot size indicates the gene ratio (proportion of input genes associated with the term), and colour indicates statistical significance. Enrichment patterns reflect expected lineage-specific immune functions, including cytotoxic activity in NK cells and antigen receptor signalling in B and T cells. **b,** Bar chart showing selected marker genes distinguishing classical monocytes (C2) and CD16$^+$ monocytes (C7). The x-axis shows the differential gene score, with positive values indicating enrichment in CD16$^+$ monocytes and negative values indicating enrichment in classical monocytes. Gene rank within each cluster's scored gene list is shown at the tip of each bar. **c,** Jaccard similarity index comparing the top-$N$ marker genes selected by our model against a post-hoc Wilcoxon rank-sum test using Seurat. The consistently sustained trajectories across most clusters reflect strong consensus with classical empirical testing.

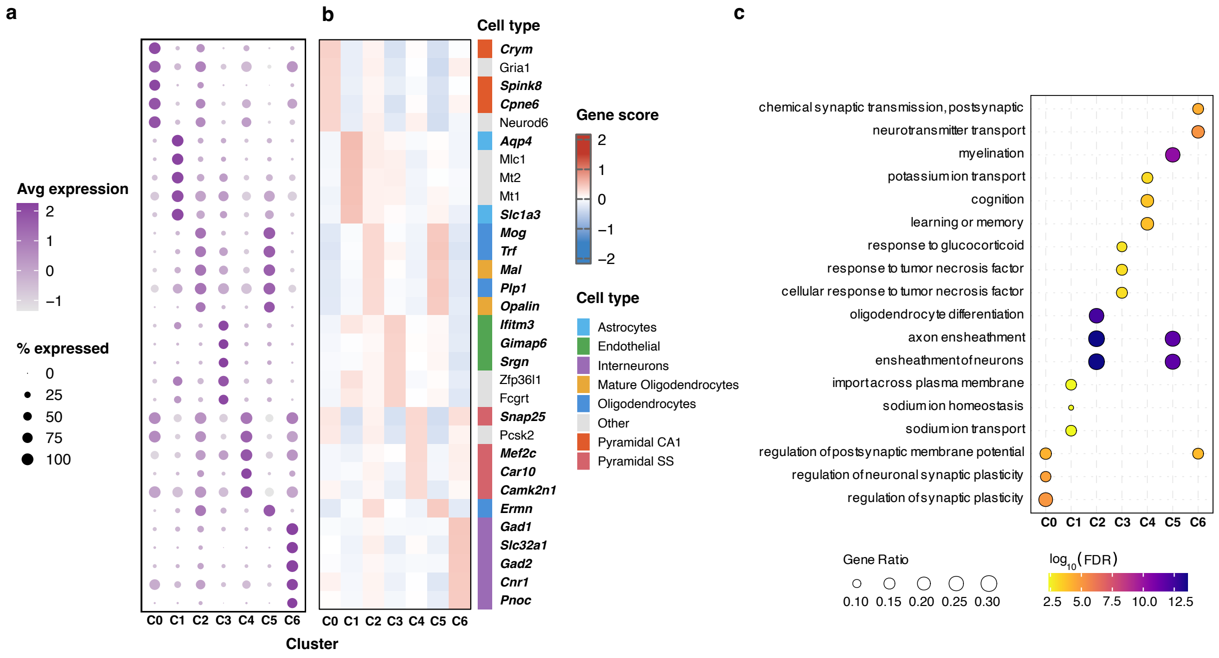


Figure S21: Characterisation of Zeisel clusters identified by scFLAME via shared scores derived from shared loading matrix. **a,** Dot plot showing expression of identified cluster marker genes across seven clusters (C0–C6). Dot size indicates the percentage of cells expressing each gene; colour indicates average scaled expression. **b,** Heatmap of gene scores (z-scored) for the top five marker genes per cluster, annotated by known cell-type identity (right). Bold italic gene names indicate canonical markers from prior literature - most are identified from the authors' own marker genes identified, with support from other sources in the literature [5–7]. Note Cluster 2 and 5 are both oligodendrocyte clusters and share many marker genes with the top 5 genes for Cluster 5 being *Trf, Mog, Plp1, Ermn* and *Opalin.* **c,** Dot plot showing GO Biological Process enrichment of the top 50 positively scored genes per cluster, showing the top 3 terms per cluster. Dot size indicates the gene ratio (proportion of input genes associated with the term), and colour indicates statistical significance. Excitatory CA1 neurons (C0) group directly with synaptic plasticity networks, whereas somatosensory (SS) neurons (C4) are associated with cognitive behaviour, and interneurons (C6) are associated with postsynaptic transmission events. Astrocytes (C1) regulate transmembrane sodium transport, and endothelial cells (C3) respond heavily to systemic inflammatory mediators such as tumor necrosis factor. Structural wrapping paths (axon ensheathment, ensheathment of neurons) are highly significant across both oligodendrocyte subsets (C2, C5).

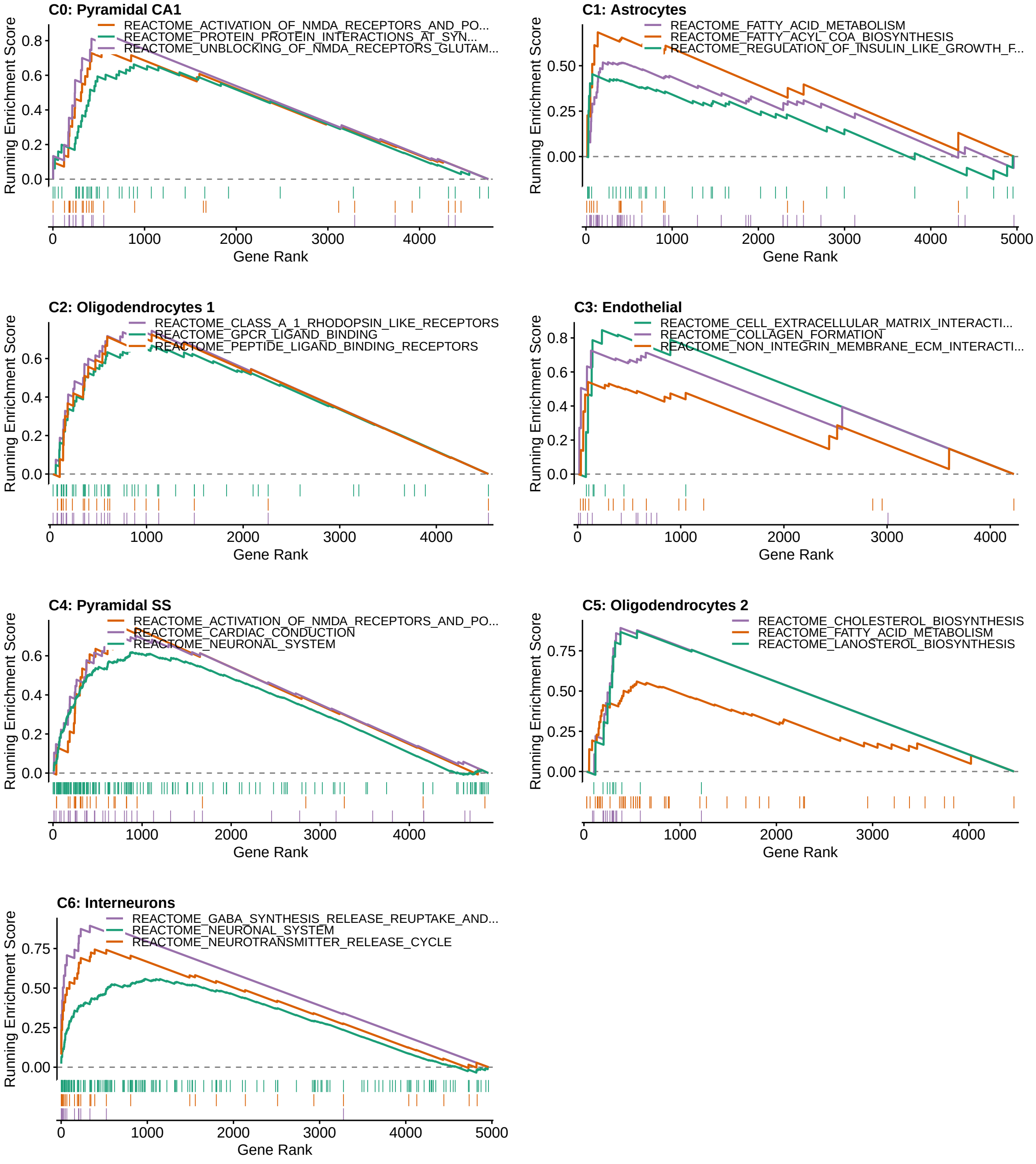


Figure S22: GSEA enrichment plots for all Zeisel clusters showing the top three enriched immunological gene sets (MSigDB M2 (Reactome) database), revealing functional networks across mouse brain cell types. All pathways are highly significant ($p_{\text{adj}} < 0.05$, NES $> 1.5$). (C0, C4) Pyramidal Neurons show strong enrichment for canonical excitatory architecture, including transmission across chemical synapses, post-synaptic signal transmission and activation of NMDA receptors. Astrocytes (C1) are characterised by upregulation in lipid processing, including fatty acid metabolism and acyl-CoA biosynthesis. Oligodendrocytes in C2 are mainly associated with GPCR and peptide ligand binding, whereas in C5, these oligodendrocytes show association with cholesterol and lanosterol biosynthesis. This could indicate these cells are actively myelinating oligodendrocytes as this biosynthesis is required for myelin sheath generation; this is further supported by the ORA dotplot in Figure S21. Endothelial cells (C3) are defined by blood-brain barrier maintenance pathways e.g. extracellular matrix (ECM) interactions and collagen formation, while interneurons (C6) are enriched for neurotransmitter (e.g. GABA) synthesis, release and reuptake cycles.

#### 4.4 scFLAME reveals hierarchical structure and cellular sub-clusters

##### 4.4.1 PBMC

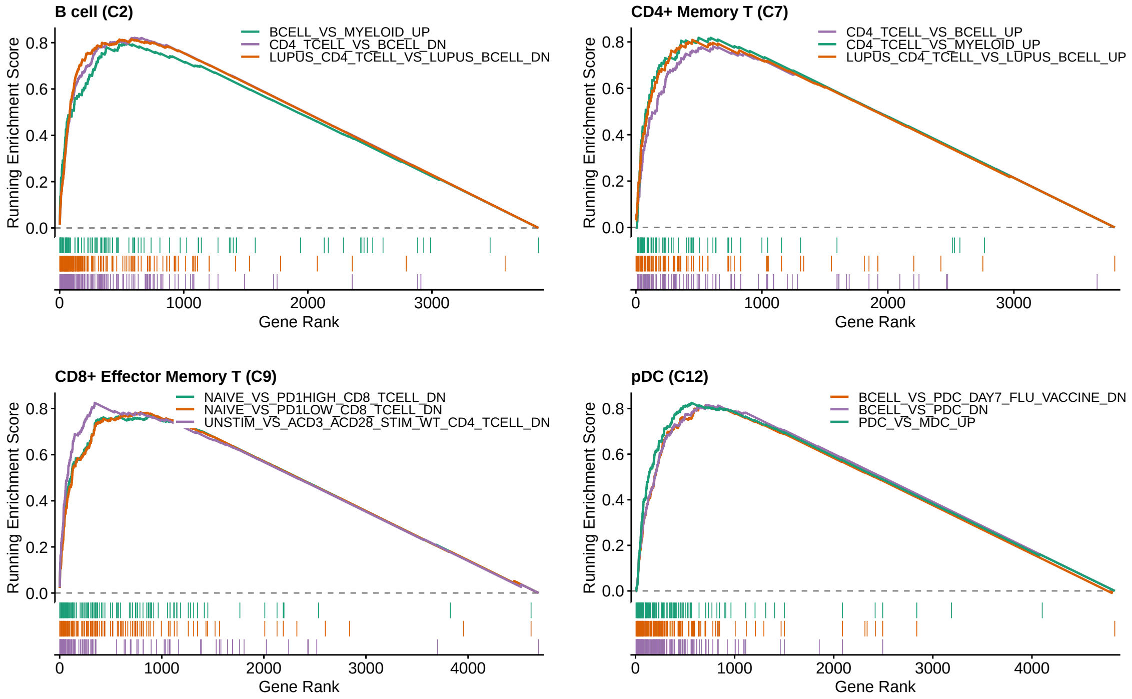


Figure S23: GSEA enrichment plots for clusters 2, 7, 9, 12 for $K = 15$ in the PBMC dataset (when initialised with $K = 15$ clusters) showing the top three enriched immunological gene sets (MSigDB C7). Running enrichment scores are shown as a function of gene rank. Even at a fine-grained resolution, scFLAME can recover a ranked list of biologically sound marker genes corresponding to known cell types.

##### 4.4.2 Baron

At $K = 5$, top marker genes of each cluster distinguish their broad identities (Figure S25c). However, we note that e.g. the gamma cell signal in Cluster 0 is suppressed by a dominance of three times as many delta cells in this shared cluster, with key marker gene *PPY* having a score of $s_{4,d} = 0.15$ (rank 43) compared to $s_{4,d} = 0.40$ for SST (rank 1). Ductal and acinar cells are grouped together at $K = 5$, separate from endothelial and stellate cells, reflecting their shared exocrine identity, distinct from the vascular and stromal compartment. At $K = 6$, there exists a beta cell sub-cluster that merges with the alpha cluster at $K = 5$ (Figure S25d); the beta identity of this population is supported by marker genes including *INS* (rank 2), but is also characterised by a stress-associated transcriptional programme including *DDIT3*, *TRIB3*, *PPP1R15A* in the top 20 genes, and canonical beta-cell markers such as *IAPP* (rank 169) and *NPTX2* (rank 1555, as opposed to rank 5 in Cluster 0) appear lower in the ranked list. The reduced representation of these markers may reduce the transcriptional similarity between Cluster 0 and Cluster 5, while the shared endocrine transcriptional programme between alpha and beta cells may facilitate the smaller beta-cell population being absorbed into the alpha-cell cluster. This suggests the merge reflects resolution-dependent clustering of a transcriptionally altered beta-cell population.

At $K = 9$, PPY distinguishes gamma cells (rank 1, $s_{4,d} = 0.43$), SST distinguishes delta cells (rank 1, $s_{7,d} = 0.36$), and REG1A distinguishes acinar cells (rank 1, $s_{2,d} = 0.69$), revealing a similar clustering structure to that discussed in the main text. We inspect marker genes for the two stellate clusters (Figure S25e), showing both sub-clusters retain positive scores across all marker genes. The largest difference in score is

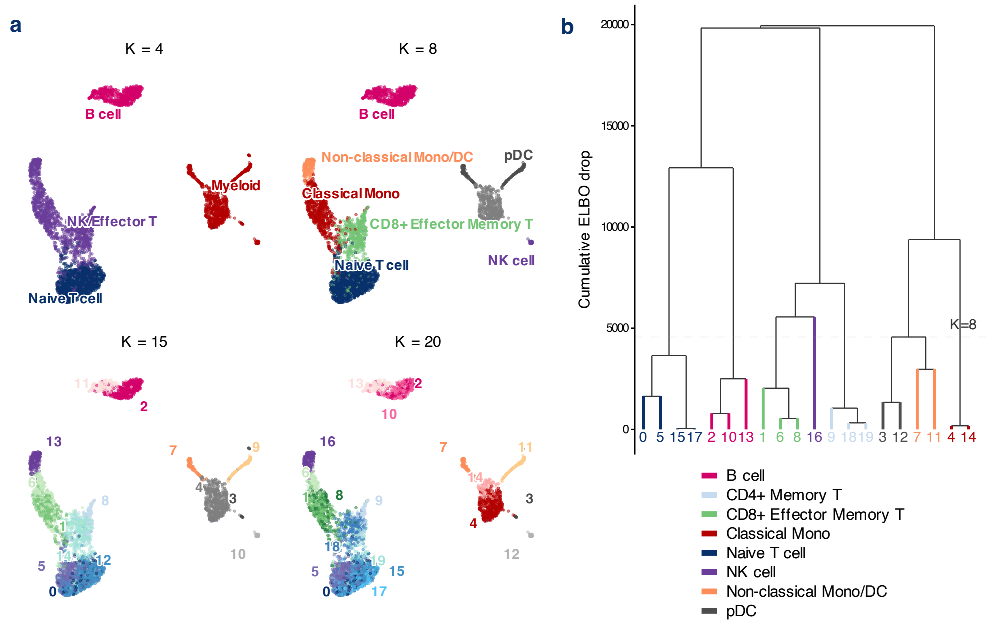


Figure S24: Visualisation of the greedy merge procedure applied to the PBMC 4K dataset with $K_{\text{init}} = 20$, showing that at a finer resolution, a biologically meaningful hierarchy is still recovered. **a,** UMAP projections of the PBMC 4K dataset coloured by cluster assignment at four resolutions of the merge hierarchy, starting from $K_{\text{init}} = 20$: $K = 4, 8, 15, 20$. Cluster identities were assigned post hoc using marker gene scores (Methods). **b,** Dendrogram showing the order of cluster merges, with branch height representing cumulative ELBO drop. Branches are coloured by their $K = 8$ parent cell type; leaves are labelled with their $K = 20$ cluster index. Dashed line indicates $K = 8$.

*DLK1* ($\Delta s = 0.54$) for the activated state, accompanied by higher scores for stromal and extracellular-matrix-associated genes including *SFRP2*, *COL6A3* and *IL11*. These differences are consistent with a more activated, matrix-remodelling stellate state, while the retention of positive scores across markers shared by both clusters suggests that the two populations remain closely related. $K = 11$ achieves the lowest DBI score in Figure S25f, reflecting the support for extra sub-clustering discussed.

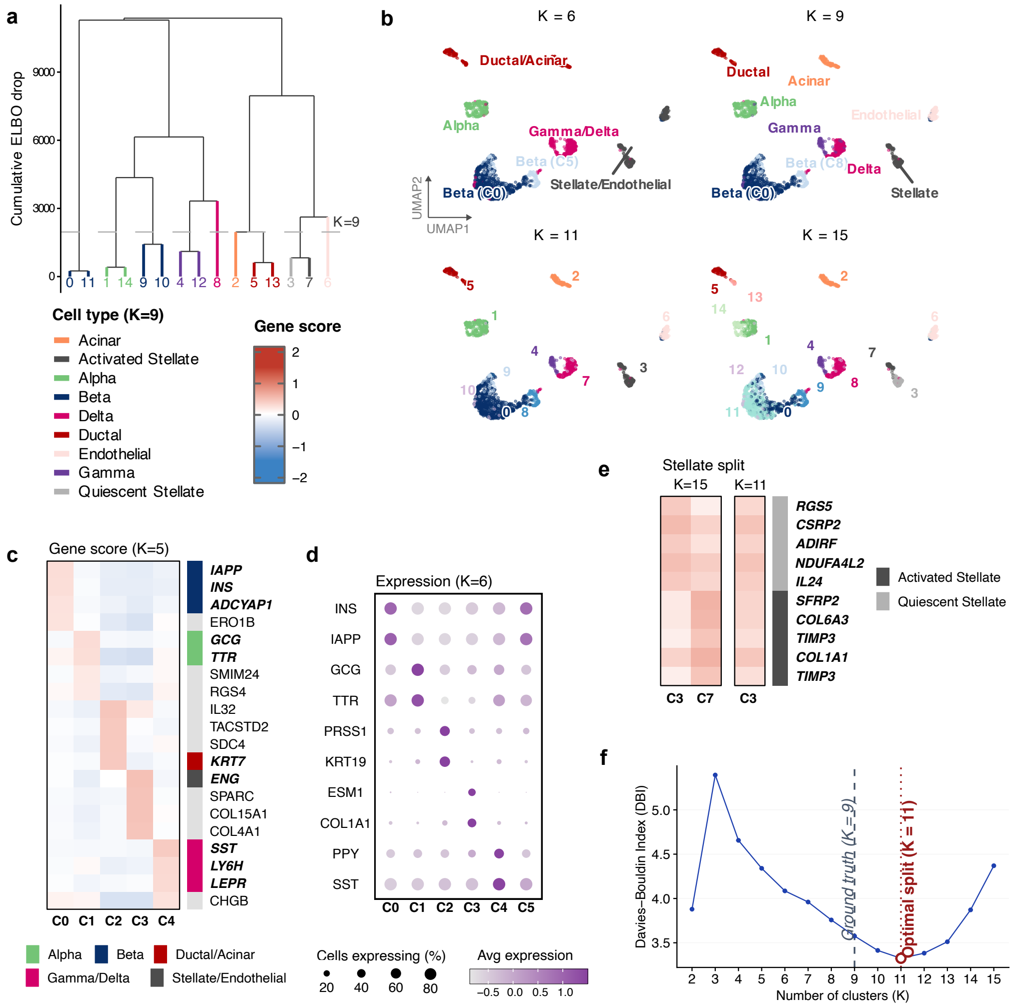

Figure S25: A greedy merge procedure recovers a biologically coherent hierarchy of cell types in pancreatic single-cell data. **a,** Dendrogram showing the order of cluster merges, with branch height representing cumulative ELBO drop. Branches are coloured by their cell type when compared to the ground truth labels; leaves are labelled with their $K = 15$ cluster index. Dashed line indicates $K = 9$. **b,** UMAP projections of the Baron pancreas dataset coloured by cluster assignment at four resolutions of the merge hierarchy, starting from $K_{\text{init}} = 15$: broad cell types ($K = 5$), cell types at the 'ground truth' number of clusters [8] ($K = 9$), and at two finer-grained resolutions ($K = 11$, the DBI defined optimal structure, and $K = 15$). Cluster identities were assigned post hoc using marker gene scores and by comparing to ground truth labels by the authors. **c,** Gene scores (Methods) for the clusters at $K = 5$, showing the top 4 genes by cluster. Known marker genes are annotated [9, 10]. **d,** Dot plot of gene expression for well-known pancreatic marker genes at $K = 6$ (*INS, IAPP*: beta; *GCG, TTR*: alpha; *PRSS1*: acinar; *KRT19*: ductal; *ESM1*: endothelial; *COL1A1*: stellate; *PPY*; gamma, *SST*: delta). **e,** Gene scores distinguishing activated stellate and quiescent stellate at $K = 15$ (clusters 3 and 7) and their merged counterpart at $K = 8$ (cluster 3). We show 5 key marker genes for each cell type in both cases from Hao et al. [9], showing those with the highest difference in score. **f,** Davies–Bouldin index across merge steps from $K_{\text{init}} = 15$ down to $K = 2$.
